# Identification of a novel MCR-1 variant displaying low level of co-resistance to β-lactam antibiotics that uncovers a potential novel antimicrobial peptide

**DOI:** 10.1101/2023.04.03.535341

**Authors:** Lujie Liang, Lan-Lan Zhong, Lin Wang, Dianrong Zhou, Yaxin Li, Jiachen Li, Yong Chen, Wanfei Liang, Wenjing Wei, Chenchen Zhang, Hui Zhao, Lingxuan Lyu, Nicole Stoesser, Yohei Doi, David L Paterson, Fang Bai, Siyuan Feng, Guo-Bao Tian

## Abstract

The emerging and global spread of a novel plasmid-mediated colistin resistance gene, *mcr-1*, threatens human health. It is accepted that MCR-1 affects bacterial fitness, and this fitness cost correlates with bacterial membrane lipid A perturbation. However, the detailed molecular mechanism remains unclear. Here, we screened out a novel MCR-1 variant, named M6, with a two-point mutation that rendered a low level of co-resistance to β-lactam antibiotics. Compared to wild-type (WT) MCR-1, this variant caused severe lipid A perturbation resulting in peptidoglycan layer remodelling and thus resulted in phenotypic co-resistance to β-lactams. Moreover, we found that a lipid A loading cavity is localized at the linker domain of MCR-1 and governs colistin resistance and bacterial membrane permeability, and the mutated pocket of M6 facilitates the binding affinity of lipid A. More importantly, synthetic peptides derived from M6 achieved broad-spectrum antimicrobial activity. These findings provide insights into a potential vulnerability that could be exploited in future antimicrobial drug design.

## Introduction

Colistin is a last-resort antibiotic against infections caused by highly drug-resistant bacteria, especially carbapenem-resistant *Enterobacterales*[1]. However, in 2016, the emergence of a novel plasmid-mediated colistin resistance gene, *mcr-1*, threatened the clinical efficacy of colistin. Since then, *mcr-1*-positive *Enterobacterales* (MCRPE) have been detected from various sources (including livestock, humans, animal food products and the environment) and have disseminated globally, spreading to >40 countries in five continents[2-4]. Colistin has been used as an additive in livestock feed to promote growth and prevent infection in China since the 1980s, and the correlation between the spread of *mcr-1* and colistin use in animal husbandry have been observed[5]. To prevent the continuous spread of plasmid-borne *mcr-1*, the Chinese government banned the use of colistin as an animal feed additive in 2017, resulting in remarkable reductions in the production and sale of colistin sulfate premix[6], and in dramatic consequent reductions in *mcr-1* prevalence. For instance, based on a 7-year regional surveillance survey, the prevalence of MCRPE in pigs and chickens gradually increased during 2015-2016, reaching a peak of 38-45%, then dropped to a trough of <2% in 2018-2019 following the banning policy[7]. However, even after the ban, a low *mcr-1* prevalence persisted in inpatients was still detected, likely associated with the approval of colistin for human clinical use in China[5, 6, 8, 9]. Notably, recent research reported that the coexistence of *mcr-1* and carbapenemase genes increased after the clinical introduction of polymyxin[10]. There is an urgent need to develop new strategies to eliminate *mcr-1* spread and prolong the use of colistin/polymixins as the last-resort antibiotic against *mcr-1*-positive carbapenem-resistant bacteria.

MCR-1 belongs to the YhjW/YjdB/YijP alkaline phosphatase superfamily and acts as an inner-membrane bound metalloenzyme that transfers the phosphoethanolamine (PEA) moiety from the phosphatidylethanolamine (PE) donor to the 1’- or 4’-phosphate group of lipid A, which is localized at the outer leaflet of the cytoplasmic membrane[11, 12]. By neutralizing the overall negative charge of the bacterial surface, MCR-1 renders the bacteria resistant to colistin. MCR-1 contains five predicted membrane-spanning α-helixes, which function as the insoluble domain and anchor in the cytoplasmic membrane. This domain is connected with a soluble catalytic fraction that is exposed in the periplasm through a flexible linker region[13, 14]. The catalytic domain of MCR-1 is highly homologous with that of the *Neisseria meningitidis* PEA transferase EptA and constitutes several β-α-β-α “sandwich” structures, which comprise an active site that must be coordinated with zinc ions[11, 15]. EptA and MCR-1 share a conserved zinc-binding pocket, which is necessary for phenotypic resistance to colistin. It is proposed that a two-step reaction, including (i) MCR-1 hydrolysing PE into diacyl glycerol and MCR-1-bound PEA as well as (ii) MCR-1 transferring the PEA moiety to lipid A, may facilitate lipid A modification with MCR-1 based on the mechanism for the enzymatic hydrolysis of EptA[16]. It is well known that MCR-1 possesses two putative functional domains for PE entry and a catalytic centre with an active site, and these two cavities are highly conserved across the entire MCR family of enzymes[17]. Although the PE-interacting cavity has been assigned[18], the precise mechanism for the interplay between MCR-1 and lipid A remains unknown.

The lipid A moiety of lipopolysaccharide (LPS), the main component of Gram-negative bacterial outer membrane (OM), is the hallmark of Gram-negative bacterial physiology and is essential for survival[19, 20]. Lipid A is synthesized on the cytoplasmic surface of the inner membrane by a conserved pathway of nine constitutive enzymes. After the attachment of the core oligosaccharide, lipid A is then flipped to the outer leaflet of the inner membrane (IM) by the ABC transporter MsbA, following by the addition of O-antigen polymer [21]. To establish OM lipid asymmetry, LPS are transported from the IM to the outer leaflet of the OM[22]. The normal distribution of lipid A is the key to maintaining the mechanical integrity and plasticity of bacterial outer membrane[23, 24]. It appears that the interaction between MCR-1 and lipid A to some extent impact bacterial physiology. Several different groups have shown that MCR-1 leads to loss in bacterial fitness[25-28]. For example, the overexpression of MCR-1 in *Escherichia coli* caused growth defect, reduced bacterial fitness and cell membrane disruption[28]. Such MCR-1 related fitness cost was also observed from hospital collected isolate[29]. The embedding of MCR-1 in the *E. coli* inner membrane and the modification of LPS with PEA contribute to this fitness loss[27]. Subsequently, it is accepted that the reduced *mcr-1* prevalence after the ban on colistin-added fodders is firmly associated with the MCR-1-mediated fitness cost in the absence of antibiotic selective pressure[6, 7, 9]. Notably, our recent work suggested that MCR-1 causes lipid remodelling that results in a defect in the outer membrane permeability, thus compromising the viability of Gram-negative bacteria[30]. These observations indicate that MCR-1 may possess a potential lipid binding pocket that disrupts lipid A distribution in the outer membrane. However, the detailed molecular mechanism by which MCR-1 disrupts lipid homeostasis remains unclear.

Here, using our previous high-throughput library, we isolated and identified an MCR-1 variant (M6) carrying a two-point mutation (P188A and P195S) at the linker domain that conferred a low level of co-resistance to β-lactams. The *E. coli* expressing this variant exhibited growth defect, cell shrinkage, peptidoglycan remodelling, and more severe membrane lipid A perturbation comparing with *E. coli* expressing WT MCR-1. Moreover, we found a lipid A loading cavity localized at the linker domain of MCR-1, and the mutated pocket of M6 may facilitate lipid A binding affinity. Strikingly, synthetic peptides derived from the M6 as “a potential cure” enabled the drug-resistant bacteria to be killed. These findings extended our understanding of the effect of MCR-1 on bacterial physiology and provided a potential strategy for eliminating drug-resistant bacteria.

## Results

### A novel MCR-1 variant, M6, caused a low level of co-resistance towards β-lactam antibiotics

The bacterial outer membrane is the first line of defence, forming a strong permeability barrier against antibiotics, whose permeabilization has been suggested to facilitate the entry of many antibiotics[31-33]. Given that recent evidence indicates that MCR-1 causes defect in the permeability of outer membranes[30, 34], we wondered if certain MCR-1 variants inducing significant OM defect may exhibit a co-resistance phenotype to other classes of antibiotics. We tested this hypothesis using our previous *mcr-1* mutant library, which included 171,769 mutation genotypes[35]. Among these genotypes, 4858/4860 (99.9%) possible single point mutations were represented. To decrease the interference of drug-induced tolerance colonies, BW25113 carrying empty vector and BW25113 carrying MCR-1 were used as controls. Screening was performed at concentrations below and above the minimal inhibitory concentration (MIC) of the antibiotics tested (0.8-2× MIC) to separate mutants that gained low- and high-level resistance (Figure S1A and Table S1). We estimated the average coverage of each single point mutation genotype in the *mcr-1* mutant library to be 10-fold. Among the colony forming unit (CFU) values from plates containing aminoglycosides (streptomycin, SM), tetracyclines (tetracycline, TET), quinolones (nalidixic acid, NAL) or glycopeptides (vancomycin, VAN), no significant difference was observed among the three strains. However, for the plates containing penicillin (ampicillin, AMP), cephalosporin (ceftazidime, CAZ) or carbapenem (imipenem, IMP) antibiotics, obvious CFUs were observed for the MCR-1 library but not for the control strains (Figure S1B). Moreover, we clarified that the MCR-1 library indeed exhibited higher viability than that of the remaining two strains after treatment with varied β-lactam antibiotics (Table S2; Figure S1C). To prevent interference caused by chromosomal mutation, 50 isolates of the MCR-1 library were selected from the plates containing 2 x MICs CAZ or AMP, and plasmids harbouring the *mcr-1* variant were first extracted from selected colonies separately, followed by transformation into a new *E. coli* BW25113 competent cell. Next, the genotypes of selected MCR-1 mutants were identified by Sanger sequencing (Table S3). In addition, the reconstructed strains were applied for MIC testing among CAZ, AMP and FOX. Several mutants displayed low-level resistance to AMP or CAZ (Table S3). Notably, a variant we called M6 was identified, carrying two mutations (P188A and P195S) localized at an α-helix of the linker domain (Table S3, Figure 1A), where exhibits low conservation among the protein structures of the MCR family (Figure S2). Compared to MCR-1-expressing cells, this mutant displayed lower sensitivity towards several β-lactam antibiotics and remained resistance towards colistin (Figure 1B and Table 1), but not for other types of antibiotics (Figure S3A). This phenotype was further confirmed by using the MCR-1 native promoter (Figure S3B). In addition, single mutation of MCR-1 with either P188A or P195S abolished the co-resistance phenotype (Figure S3C), indicating the requirement of simultaneous mutations.

**Figure 1.**
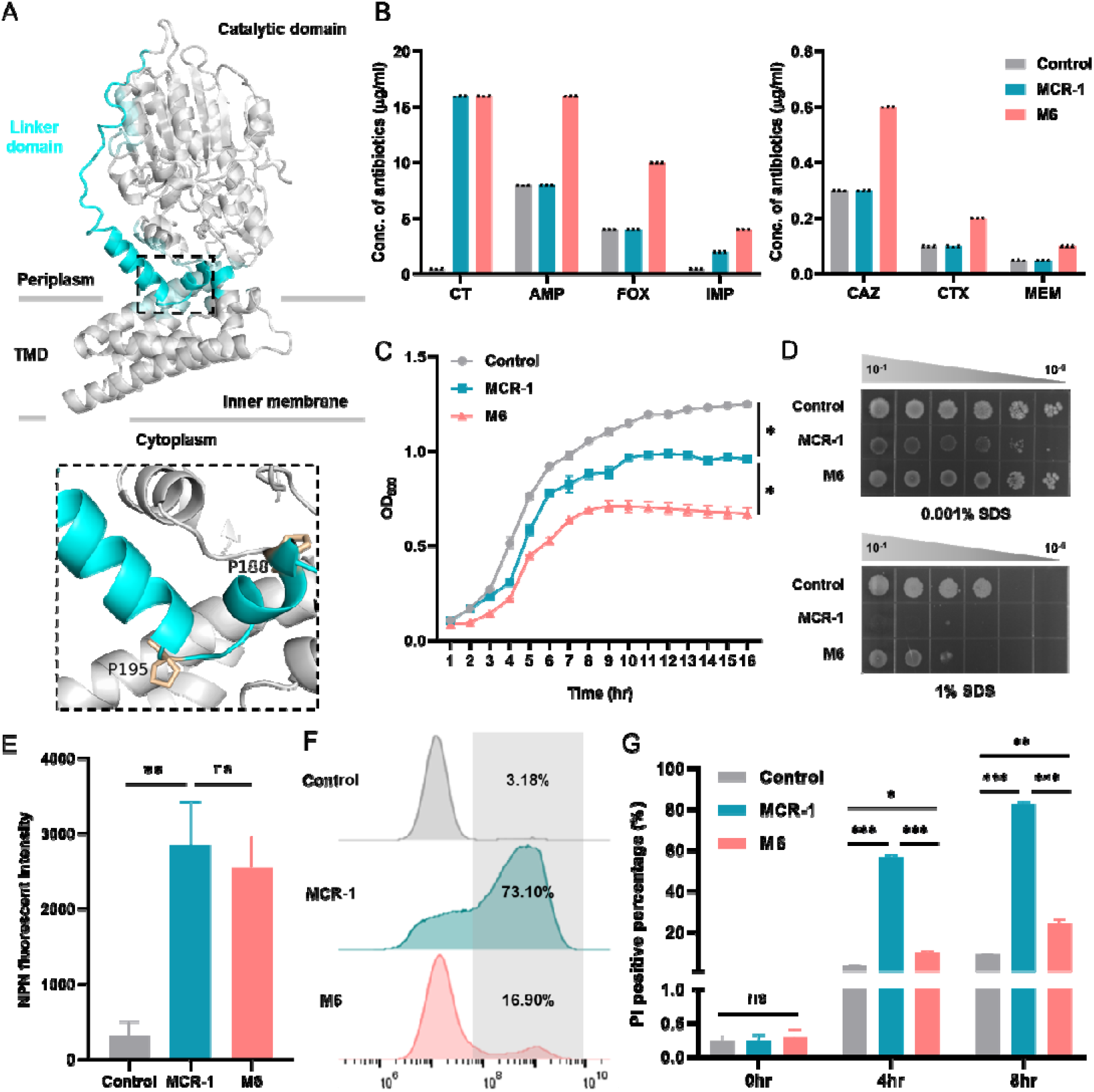
An MCR-1 mutant M6 (P188A and P195S) exhibited reduced sensitivity towards β-lactam antibiotics. **(A)** MCR-1 protein structure in cartoon. The linker domain (blue) and approximate membrane boundaries are indicated. TMD, transmembrane domain. Close-up view of two mutated residues. The mutated residues P188 and P195 are highlighted in orange. **(B)** Assessment of the sensitivity of M6-expressing cells to colistin and β-lactam antibiotics. **(C)** Growth curves of BW25113 carrying empty vector, M6, or MCR-1. The y-axes show optical densities at 600 nm (OD_600_) of broth cultures, x-axes show period of growth (hr). **(D)** Efficiency of plating assays on LB agar plates containing 1% SDS and 1 mM EDTA or 0.001% SDS and 1 mM EDTA. Ten-fold dilutions of cultures are indicated above the left plate. **(E)** The outer membrane integrity of strains carrying M6 or MCR-1 was determined by measuring NPN uptake during logarithmic phase. **(F-G)** The inner membrane permeability of cells carrying MCR-1 or M6 was evaluated by PI staining assay. Overnight cultures were sub-cultured into fresh LB broth at a ratio of 1:100 and induced with 0.2% arabinose to express MCR-1 or MCR-6. After induction for 4 hr and 8 hr, stationary and late-stationary phase cultures were collected, respectively, followed by staining with PI dye for 15 min. The PI-positive proportion was determined by flow cytometry and analysed by FlowJo version 10 software. All the above-described experiments were performed thrice with similar results. Error bars indicate standard errors of the means (SEMs) for three biological replicates. A two-tailed unpaired *t* test was performed to determine the statistical significance of the data. ns, no significant difference; *, *P*< 0.1; **, *P*< 0.01; ***, *P*< 0.001.

**Table 1.**
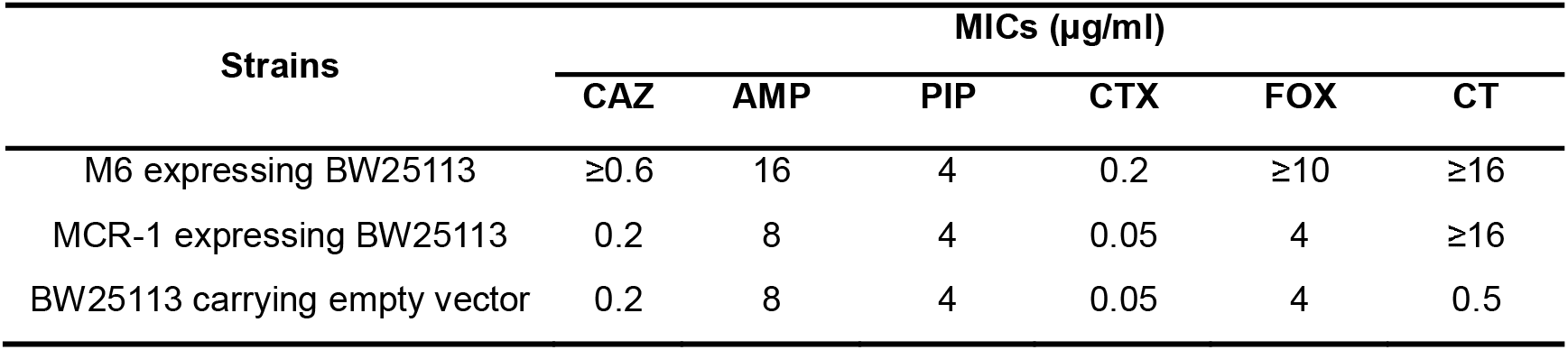
β-Lactam antibiotics MICs of M6.

Interestingly, M6 expression significantly retarded bacterial growth compared to that of *E. coli* expressing MCR-1 (Figure 1C). Based on antimicrobial susceptibility assays, we speculated that this phenotypic co-resistance may result from a decrease in the outer membrane permeability of M6. To do it, we first assessed the susceptibility of M6 to SDS/EDTA, which were external agents that can damage the cell and prevent growth by disrupting lipid stability and solubilizing the outer membrane. Consistent with our previous observation[30], MCR-1-expressing cells displayed increased sensitivity to detergents compared with that of the empty plasmid control, whereas M6 expression to some extent recovered host permeability (Figure 1D). We further quantified the degree of membrane permeability by staining the indicated strains with the small molecule n-phenyl-1-napthylamine (NPN), reflecting leakiness in the outer membrane, or propidium iodide (PI), a marker to represent the integrity of the bacterial inner membrane. As expected, an enhancement in NPN uptake was observed in the MCR-1-expressing cells, but the M6-expressing cells exhibited similar level in NPN uptake compared to that of MCR-1-expressing cells (Figure 1E). In contrast to the observation in the NPN assay, M6 expression significantly decreased the number of PI-positive cells at stationary phase compared to that of MCR-1-expressing cells (Figure 1F-G), suggesting that the M6-mediated co-resistance phenotype might be possibly related with membrane damage.

Overall, these results revealed that M6 conferred a low level of co-resistance towards β-lactam antibiotics via an unexpected mechanism affecting the inner cell membrane.

### More severe lipid A disorder caused by an M6-induced co-resistance phenotype

Next, we wanted to explore the molecular mechanism by which M6 caused a low level of co-resistance towards β-lactam antibiotics. Bacterial PG layer remodelling has been suggested to reduce the susceptibility to β-lactams, and such remodelling can be triggered due to the OM assembly defect[36-38]. To further confirm the OM integrity of M6, scanning electron microscopy (SEM) was performed for target strains. Smooth and intact surface was observed for *E. coli* BW25113 bearing empty plasmid vector but wrinkles for MCR-1-expressing cells. Significant morphological changes, however, were observed on M6-expressing cells, which showed folded membrane stacks and holes at bacillus poles (Figure 2A). This result firmly demonstrated that the expression of M6 resulted in severe OM damage. Moreover, transmission electron microscopy (TEM) revealed that both cells expressing MCR-1 and cells expressing M6 displayed cell wall shrinkage, in which an obvious interspace was formed within the periplasm at one pole of the cell (Figure 2B), indicating shrinkage of cytoplasm. To quantify the degree of bacterial membrane perturbation, a two-fluorescence reporter system was constructed to separate the bacterial periplasm and cytoplasm according to previous research[39], in which super-folded green fluorescent protein (sfGFP) was fused with the signal peptide of DsbA localized in the periplasm and mCherry was constitutively expressed in the cytoplasm (Figure S4A). Cultivated in LB broth, stationary phase cultures were analysed. The control showed an extremely low percentage of isolates carrying foci at cellular pole(s). M6-positive cells exhibited a high proportion of cells that accumulated sfGFP at bacterial pole(s) (90.7%, Figure S4B), which was higher than the proportion of MCR-1-positive cells (61.3%). Since IM shrinkage was accounted for the formation of sfGFP foci at *E. coli* bacterial pole(s)[39], this result suggested that the expression of M6 contributed to more severe membrane perturbation than MCR-1.

We next employed label-free quantitative proteomics to profile proteome changes. In total, 2287 proteins were identified, representing 55.5% coverage of the *E. coli* BW25113 proteome (Figure S5A, Table S4). With our significant criteria (|log2-fold change (M6/MCR-1)|≥1 and *p value* < 0.05), 66 proteins were differentially expressed (41 upregulated and 25 downregulated). Notably, Gene Ontology (GO) analysis revealed a significant enrichment in categories that were consistent with the key biological events related to bacterial cell membrane and envelope space (Figure S5B, Table S5). Among the top enriched processes revealed by KEGG analysis, enhanced process of peptidoglycan (PG) layer synthesis was observed (Figure S5C, Table S6). In particular, AmpH, PBP6 and LdtD, the proteins related to PG layer remodelling, were significantly upregulated (Figure 2C). A proteinJprotein interaction (PPI) analysis was performed to reveal the genes involving PG layer remodelling, and the network suggested that, besides AmpH and PBP6, LpoB and PBP1B might be also important regulators in the pathway (Figure S5D). In line with this finding, the transcriptional levels of genes involved in the envelope stress response and PG layer remodelling, such as *yiiO* (gene encoding CpxP), *yaiH* (gene encoding AmpH), *ycbB* (gene encoding LdtD) and *yaeL* (gene encoding RseP), were significantly upregulated (Figure 2D), indicating that M6 mainly influenced remodelling of the PG layer in the periplasm.

In *E. coli*, approximately 93% of cross-links in the PG layer are the 4-3 type, while 7% of the 3-3 cross-links formed between two *meso*-Dap residues of adjacent stem peptides[40]. LdtD is a kind of L, D-transpeptidases (LDTs) that contributes to PG layer remodelling by increasing the percentage of 3-3 cross-links[38, 41]. The activity of LdtD requires binding with the following partner proteins: PBP1B, a bifunctional PG synthase, and its GTPase activator LpoB, which are essential for the formation of the 3-3 cross-linking PG layer in LdtE and LdtF (growth-phase dependent LDTs) deficient *E. coli*[38]. β-lactam antibiotics mimic the D-Ala^4^-D-Ala^5^ termination of peptidoglycan precursors and inactivate the D, D-transpeptidases (DDTs) by acting as suicide substrates, which therefore prevents the formation of 4-3 cross-links and caused cell wall lysis[42, 43]. The increased percentage of 3-3 cross-links in PG layer allows the survive of Gram-negative bacteria under the treatment of β-lactam antibiotics[44]. We therefore explored the correlation between M6 and LdtD. With the addition of 3.75 mM CuSO_4_, an inhibitor of LDTs[45], the CAZ MIC of the M6-expressing strain was lower than that of the untreated group, while the AMP MIC of M6-expressing cells exhibited the same level as the empty plasmid control and WT MCR-1 (Figure S6A). Moreover, *mrcB* (gene encoding PBP1B) deficiency reduced the MICs of the M6-expressing strain to the same as those of the empty plasmid control and MCR-1-expressing cells (Figure 2E), although deleting *ycbB* (gene encoding LdtD) and *ycfM* (gene encoding LpoB) had no influence on the susceptibility of M6 towards β-lactam antibiotics (Figure S6B), demonstrating that PBP1B was dominant in the M6-mediated co-resistance phenotype. Furthermore, only the lack of PBP1B (i.e. Δ*mrcb*) in M6-expressing cells decreased the resistance to SDS detergent (Figure S6C). Strikingly, no significant difference in NPN uptake was observed between M6-expressing WT *E. coli* and *mrcB* null mutant (Figure S6D). Whereas the percentages of PI-positive cells of M6-bearing Δ*mrcB* mutant during stationary and late-stationary phases were dramatically higher than those of the WT strain, and the values were increased from 11.7% and 13.64% to 48% and 80.10%, respectively (Figure S6E-F). Given that the PG remodelling could reduce the susceptibility to SDS and the uptake of PI dye, these results suggested that the formation of 3-3 cross-links in PG layer might also contribute to the barrier function of IM. Overall, these results demonstrated that the LDTs-mediated PG remodelling caused by M6 expression resulted in phenotypic co-resistance.

Since our morphological evidence suggests that M6 exacerbated cell shrinkage and caused PG layer remodelling, we hypothesized that the phenotypes conferred by M6 might result from severe membrane lipid A disorder. By determining the protein expression level of PbgA, a periplasmic protein that senses lipid A levels and whose expression level regulates the content of OM LPS[46], we found that the PbgA level of M6-expressing cells was higher than those of both empty plasmid control and MCR-1-expressing cells (Figure 2F-G). Moreover, the overall level of LPS in M6-expressing cells was significantly lower than that in the control and MCR-1-expressing cells (Figure 2F-G). We therefore assumed that the increased permeability of MCR-1-positive cells and M6-positive cells was due to the reduced level of lipid A. To test this hypothesis, we generated strains overexpressing LpxC (an essential enzyme for lipid A biosynthesis) and an increase in the LPS expression level was observed (Figure S7). Consistently, outer membrane and inner membrane permeabilizations caused by MCR-1 or M6 expressions were disappeared with the compensation of LpxC (Figure 2H-I), indicating that the increased permeability of MCR-1 or M6 positive strains was due to the reduced membrane lipid A level. In addition, the overexpression of LpxC abolished the co-resistance phenotype conferred by M6 (Figure 2J), revealing the association between membrane lipid A level and co-resistance to β-lactam antibiotics in M6-bearing strain.

Taken together, these results demonstrate that severe lipid A defect caused by M6 expression exacerbated PG layer remodelling, which conferred reduced β-lactams sensitivity.

### Uncovering an essential lipid A binding pocket

Given the observations caused by this gain-of-function mutation, we hypothesized that the linker region at which the point mutations occurred might play a role in the function of the MCR-1 enzyme. To test this hypothesis, we generated a mutant of *E. coli* BW25113 MCR-1 lacking P188-P195 at the linker domain. As expected, ΔP188-P195 mutation abolished the colistin resistance of *mcr-1*-bearing strain (Figure S8A) and resumed the susceptibility to β-lactams (Figure S8B) and SDS tolerance (Figure S8C). Next, we wondered whether this region might be the potential lipid A loading pocket of MCR-1. The native conformation of lipid A was subjected to molecular docking with MCR-1. A putative lipid A binding cavity that localized at the region where the point mutations occurred was proposed. Lipid A appeared to lie in this cavity of MCR-1, and a total of 18 residues are involved in the formation of this substrate cavity for lipid A binding (Figure 3A). It seems like that four regions were critical for anchoring lipid A in the loading cavity (Figure 3B and Figure S9), since there is an intact lipid A molecule with six flexible fatty acyl chains (C12–C14). To confirm this region that interacted with lipid A, structure-guided, site-directed mutagenesis was performed. Functional assays informed us that these mutants exhibited different roles in the context of colistin resistance (Figure 3B). For example, a hydrogen network was observed within K211, K204 and another phosphorate group of lipid A, and the two-point mutation K211A+K204A of MCR-1 significantly weakened the binding of lipid A. Notably, a large hydrophobic region consisting of L64, I65, L68, L69, I165 and I168 may be responsible for the stabilization of the flexible fatty acid tails of lipid A. Consistently, the MCR-1 mutant harbouring L64A+I65A+L68A+L69A mutations completely inactivated the function of MCR-1. Since our previous work suggested that the MCR-1 lipid A binding ability is closely linked with cell envelope permeability[30], we hypothesized that mutation in the putative lipid A cavity might to some extent mitigate the membrane permeabilization. Strikingly, the mutations that abolished the colistin resistance ability of MCR-1 also resumed the integrities of both OM and IM to the same as those of control (Figure 3B). Moreover, the colistin susceptibility phenotype for these mutants was highly correlated with the integrity of the inner membrane (ρ=0.93, *P*=0.00026) or outer membrane (ρ=0.93, *P*=0.00026) (Figure 3C). In short, these results strongly suggested that the linker region of MCR-1 where the P188A+P195S mutations occurred was the lipid A binding pocket, which governed phenotypic colistin resistance and bacterial membrane integrity.

**Figure 2.**
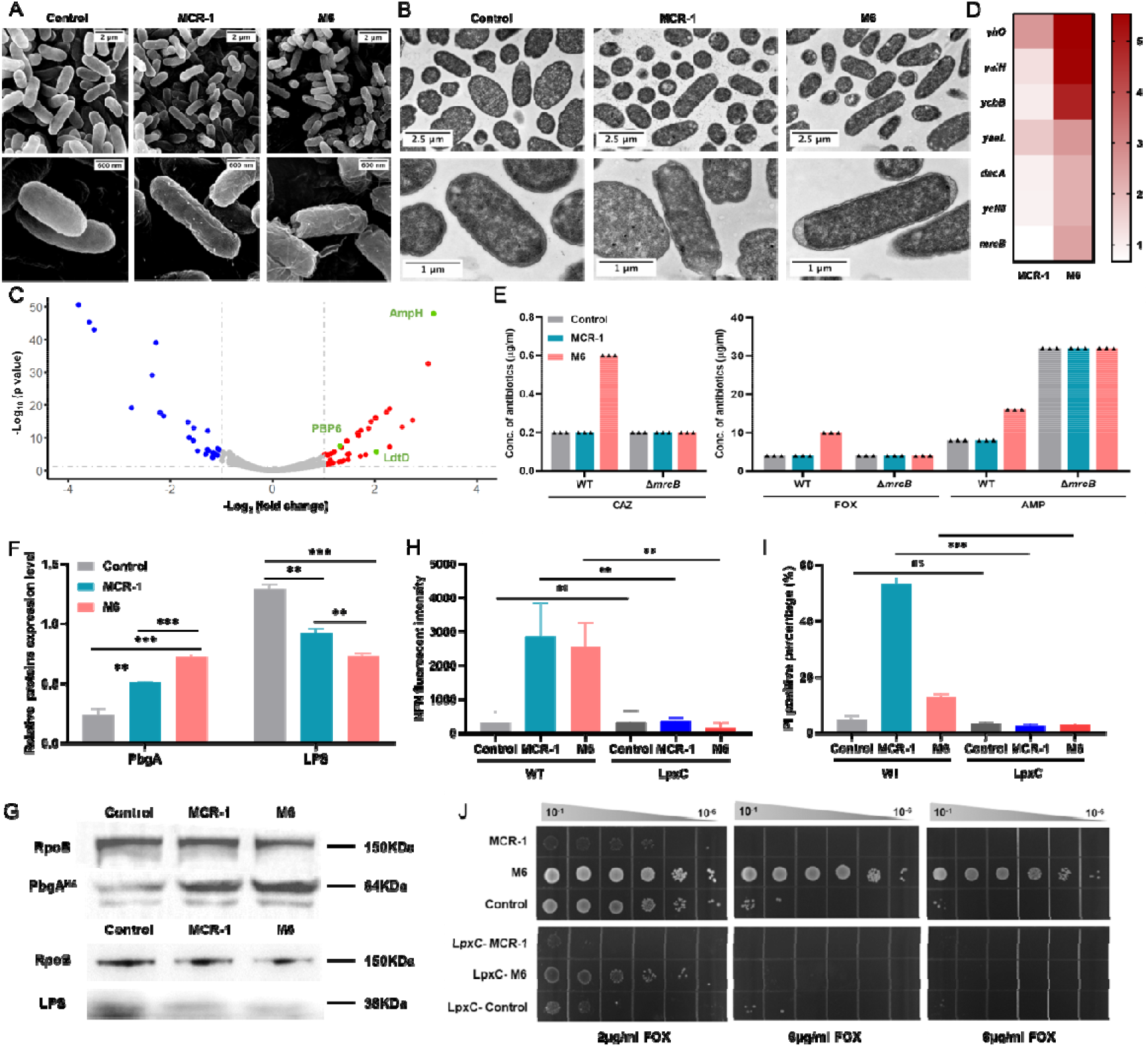
M6 caused peptidoglycan (PG) remodelling via the induction of severe lipid disorders. **(A)** SEM micrographs of BW25113 carrying MCR-1 or M6. Overnight cultures of the above strains were sub-cultured into fresh LB broth at a ratio of 1:100 and induced with 0.2% arabinose for 2 hr before sample preparation. **(B)** TEM micrographs of BW25113 carrying MCR-1 or M6. Overnight cultures of the above strains were sub-cultured into fresh LB broth at a ratio of 1:100 and induced with 0.2% arabinose for 2 hr before sample preparation. **(C)** The volcano diagram of the differentially expressed protein between BW25113 carrying MCR-1 and M6 determined by label-free quantitative proteomics. The differential expression threshold was set as log_2_ fold change >1 and *p* value <0.05. Red dots represent significantly upregulated genes, while blue dots represent significantly downregulated genes. The genes related to PG layer remodelling are highlighted in green. **(D)** The transcriptional level of differentially expressed genes was measured by qRT_JPCR, which was normalized to the transcript level of the housekeeping gene *rpoB* and quantified with ΔΔCT analysis. The heatmap represents the fold change in target gene transcriptional levels for MCR-1-expressing cells or M6-expressing cells compared with those of the empty plasmid control. **(E)** Evaluation of the impact of *mrcB* on the M6-mediated co-resistance phenotype. The antibiotic susceptibility of WT or Δ*mrcB* carrying empty plasmid (control), MCR-1 or MCR-6 was determined by agar dilution MIC tests. **(F-G)** The expression levels of PbgA and LPS were determined by western blot, along with BW25113 carrying empty plasmid (control) as a control. **(H)** Evaluation of the impact of LpxC overexpression on M6-mediated OM permeability. The outer membrane integrity was determined by measuring NPN uptake during logarithmic phase. **(I)** Evaluation of the impact of LpxC overexpression on M6-mediated IM integrity. The inner membrane permeability was evaluated by PI staining assay. **(J)** Assessment of the impact of LpxC on the M6-mediated co-resistance phenotype. The FOX MICs of WT or LpxC overexpression strains carrying empty plasmid (control), MCR-1 or M6 were determined by agar dilution MICs tests. All the above-described experiments were performed three times with similar results. Error bars indicate standard errors of the means (SEMs) for three biological replicates. A two-tailed unpaired *t* test was performed to determine the statistical significance of the data. ns, no significant difference; **, *P*< 0.01; ***, *P*< 0.001.

**Figure 3.**
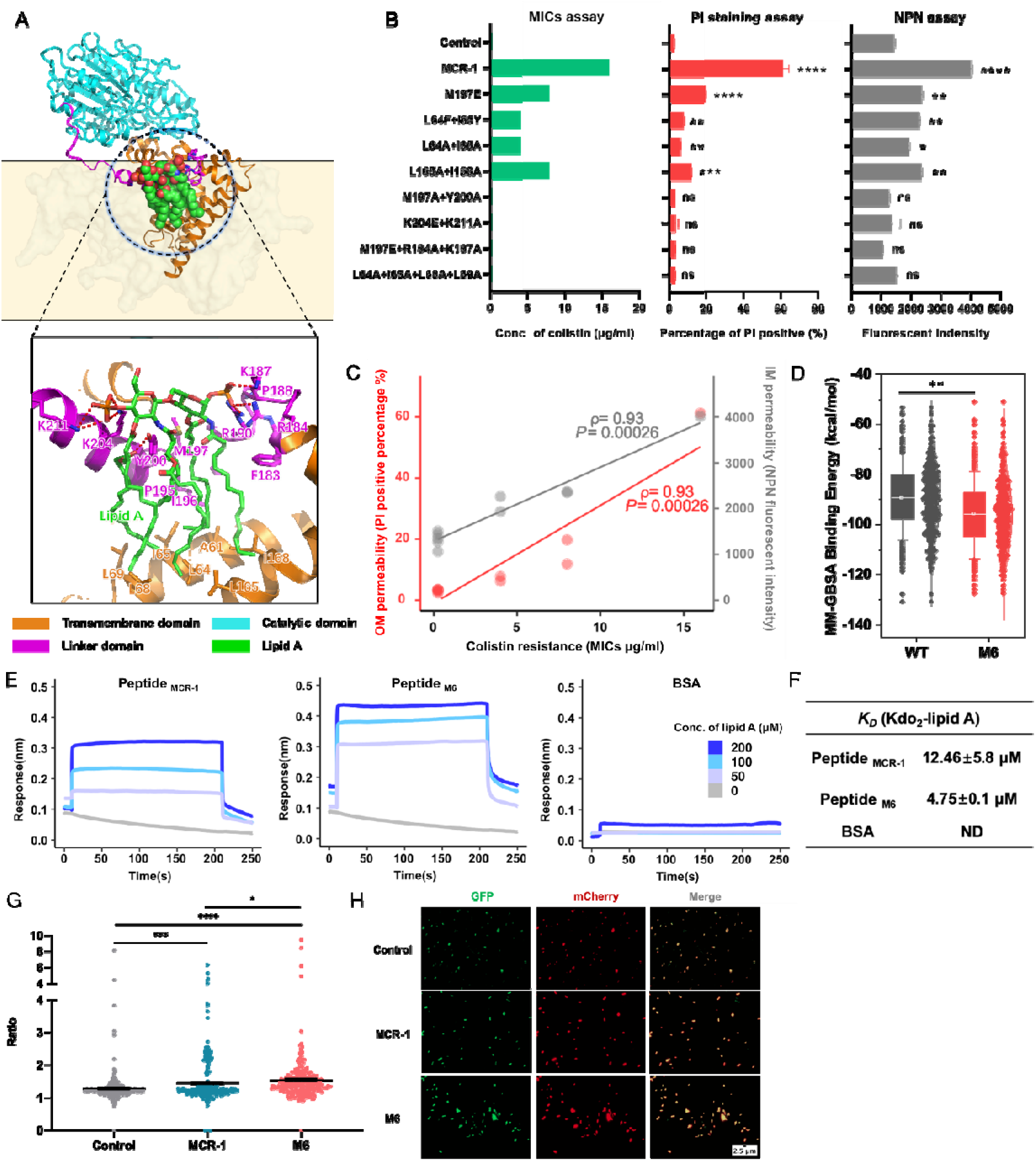
Discovery and functional definition of the lipid A loading cavity in MCR-1. **(A)** Protein structure of MCR-1 and close-up view of the LPS binding cavity at the linker domain. The catalytic domain, linker domain and transmembrane domain are in cyan, magenta and orange, respectively, with LPS is shown as green sticks and salt bridges for interaction as red dashes. **(B)** Verification of the key residues in the lipid A binding cavity that interacts with lipid A. To evaluate the influence of mutations on colistin resistance activity, the colistin MICs of the indicated mutants were determined by agar dilution MIC tests. In addition, logarithmic phase cultures of target mutants were applied to evaluate the permeability of the inner membrane and outer membrane by NPN assay and PI staining assay, respectively. **(C)** Evaluating the correlation between cell membrane integrity and colistin resistance. ρ is Spearman’s rank correlation coefficient. The related *P* value and regression line (blue) are shown. **(D)** Estimations of the binding free energies of Lipid A against WT MCR-1 and M6 were calculated. The calculation was performed over the last 200 ns trajectory after reaching equilibrium by using MM-GBSA. **(E-F)** Synthetic biotinylated MCR-1-derived lipid A-binding peptides were transferred into different concentrations of lipid A, and the affinity between synthetic peptides and lipid A was determined by interferometry measurements. **(G)** The fluorescence ratio indicates the relative membrane voltage in indicated *E. coli* cells by using a genetically-encoded voltage sensor Vibac2. **(H)** Representative images of GFP responds to membrane voltage and the mCherry used to normalize protein expression were as shown. All the above-described experiments were performed three times with similar results. Error bars indicate standard errors of the means (SEMs) for three biological replicates. A two-tailed unpaired *t* test was performed to determine the statistical significance of the data. ns, no significant difference; *, *P*< 0.1; **, *P*< 0.01; ***, *P*< 0.001; ****, *P*< 0.0001.

Next, we hypothesized that M6 might exhibit higher affinity for lipid A, since expression of M6 significantly perturbed the membrane lipid A homeostasis. To this end, we performed molecular dynamics simulations over M6 and MCR-1 to obtain probable binding conformations and evaluate their binding stabilities (Figure S10A). The binding affinities between lipid A and WT MCR-1 or M6 were evaluated by using the MM-GBSA method, and the results showed that the binding affinity of lipid A to M6 was slightly higher than that of WT MCR-1 (Figure 3D and S10B). Moreover, as illustrated in Figure S10C-D, the frequencies of the interactions between lipid A and certain residues of M6 were different from those of MCR-1. For example, the mutations P188A+P195S increased the interaction frequency between K204 and lipid A but weakened the binding between R190 and lipid A, two hydrogen bonds were generated between R190 and lipid A in M6 (Figure S10E), which were not observed in MCR-1. Finally, synthetic peptides encompassing the lipid A-binding cavity from MCR-1 or M6 were utilized to interact with LPS, with dissociation constant (*K*_D_) values of approximately 12.46 nM and 4.75 nM (Figure 3E-F), respectively, further demonstrating that a potential lipid A binding pocket at the linker domain of MCR-1 and this pocket of M6 facilitated the lipid A binding affinity. Furthermore, using a sensitive bacterial membrane voltage V_m_ sensors[47], this two-point mutation appeared to increase the membrane voltage of *mcr-1* bearing *E. coli* (Figure 3G-H).

### Synthetic peptides derived from the MCR-1 variant achieve broad-spectrum activity

Inspired by the previous research of Thomas *et a*l[46], we postulated that peptides derived from the lipid A binding cavity of MCR-1 or M6 might be utilized as antimicrobial peptides (AMPs). To do this, we first designed two 24AA short peptides based on the lipid A binding cavity of WT MCR-1 and M6, termed 24AA-WT and 24AA-2M, respectively. Peptide 24AA-WT showed no antimicrobial activity. In contrast, peptide 24AA-2M was active against *E. coli* ATCC25922 and maintained its activity during the first 6 hr (Figure 4A). However, the viability of 24AA-2M treated strain resumed to the same as that of the 24AA-WT treated one at 24 hr after initiation, suggested inactivation of the synthetic peptide. To increase the stability of indicated peptide, we generated a derivative of 24AA-2M by adding a five-tryptophan (5W) tag at the C-terminus, which was named 19AA-2M-tag, since the tryptophan C-terminal end-tagging modification generated AMPs against protease and increased peptide stability[48-51]. Promising results were obtained with the 19AA-2M-tag, which showed potent activity against ATCC25922 and maintained its activity at least up to 24 hr. And no obvious antimicrobial activity was observed for peptide consisted with five tryptophan (5W) only. Using SEM, we observed that the treatment of 24AA-2M to *E. coli* caused severe OM shrinkage, while the surface of ATCC25922 treated with 19AA-2M-tag was wrinkled and had crevasses (Figure 4B), indicating that both 24AA-2M and 19AA-2M-tag kill bacteria by damaging cell membrane. Moreover, the treatments with 24AA-2M or 19AA-2M-tag both increased the PI-positive proportion of *E. coli* ATCC25922 in a dose-dependent manner, while 19AA-2M-tag exhibited higher antimicrobial activity (Figure 4C-D). Furthermore, comparing with 24MM-2M, no general membrane lytic activity was observed towards mouse red blood cells treated with 19AA-2M-tag, indicating that 19AA-2M-tag might be a promising antimicrobial peptide (Figure S11).

**Figure 4.**
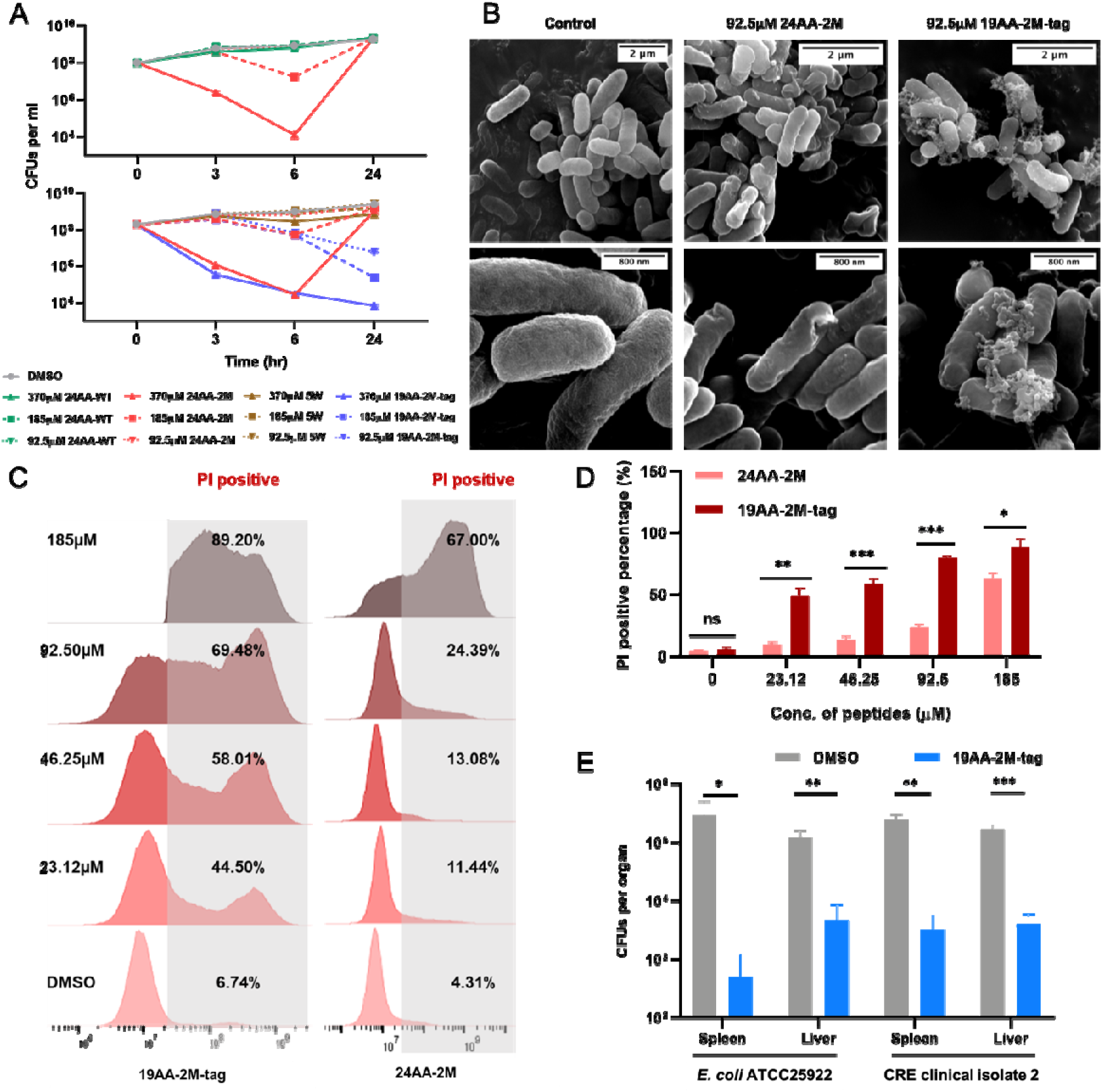
Synthetic peptides derived from the MCR-1 variant displayed antimicrobial activity *in vitro* and *in vivo*. **(A)** Time-dependent killing of *E. coli* ATCC25922 by 24AA-WT, 24AA-2M, or 19AA-2M-tag. An exponential culture of *E. coli* ATCC25922 was challenged with the corresponding peptides. Cultures treated with DMSO or 5W were set as control. **(B)** Scanning electron microscopy analysis of *E. coli* ATCC25922 treated with 0.25× MIC 24AA-2M or 19AA-2M-tag. **(C-D)** Evaluation of the bacterial membrane permeability with the treatment of 24AA-2M or 19AA-2M-tag *in vitro* by PI staining assay. **(E)** Evaluation of the bactericidal ability of 19AA-2M-tag *in vivo*. C57BL/6 mice were first infected with 1×10^7^ CFUs of *E. coli* ATCC25922 or CRE isolated in the clinic through intraperitoneal injection for 1 hr, followed by treatment with 200 μg of the indicated peptides. The infected individuals treated with DMSO were used as controls. Total mouse liver or spleen bacterial burdens were determined 16 hours after treatment by spotting serial dilutions of tissue homogenates on LB plates. The average CFUs values of each organ with (blue) or without (grey) 19AA-2M-tag treatment were counted 16 hr after incubation at 37 °C. All the above-described experiments were performed thrice with similar results. Error bars indicate standard errors of the means (SEMs) for three biological replicates. A two-tailed unpaired *t* test was performed to determine the statistical significance of the data. *, *P*< 0.1; **, *P*< 0.01; ***, *P*< 0.001.

We tested the susceptibility of 19AA-2M-tag among three *Escherichia coli* strains and clinically relevant pathogens, including *Pseudomonas aeruginosa*, *Salmonella* Typhimurium, *Klebsiella pneumoniae* and *Acinetobacter baumannii* (Table 2). MICs of 185-370 μM were obtained against these Gram-negative strains. Moreover, we examined the efficiency of 19AA-2M-tag towards clinical isolates obtaining drug resistance through different mechanisms. When comprising amikacin-resistant *A. baumannii*, carbapenem-resistant Enterobacterales, and even colistin-resistant *E. coli* (*mcr-1*^+^), all these clinical strains revealed susceptibility to the synthetic peptide with MICs ranging from 92.5 to 370 μM. For most Gram-negative strains, the IC_50_ values were similar, ranging from 27.56 to 47.19 μM, suggesting that 19AA-2M-tag, the peptide derived from the colistin resistance gene, exhibited effective antimicrobial activity against antibiotic-resistant isolates (Figure S12). Interestingly, 19AA-2M-tag displayed synergetic activity with colistin (CT) against *E. coli* ATCC 25922 (FICI=0.312) and a carbapenem-resistant *Enterobacteriaceae* (CRE) strain collected from the clinic (FICI=0.156, Figure S13A). And the addition of 46.25 μM target peptide improved the antimicrobial activity of colistin against *E. coli* ATCC25922 (Figure S13B). Finally, the *in vivo* efficacy of the 19AA-2M-tag was demonstrated in mouse models of peritonitis against the *E. coli* ATCC25922 and CRE clinical isolate. By counting CFUs in the liver and spleen 16 hr after treatment, 19AA-2M-tag treatment resulted in a significant reduction in the bacterial burden than the untreated group (Figure 4E).

**Table 2.** 19AA-2M-tag exhibits broad-spectrum antibacterial activity.

| Strains | MICs ($\mu\text{M}$) |
| --- | --- |
| *Escherichia coli* BW25113 | 185 |
| *Escherichia coli* MG1655 | 370 |
| *Escherichia coli* ATCC25922 | 370 |
| *Pseudomonas aeruginosa* ATCC27853 | 185 |
| *Salmonella enterica* serovar Typhimurium SL1344 | 185 |
| *Klebsiella pneumoniae* ATCC13883 | 185 |
| *Acinetobacter baumannii* ATCC19606 | 185 |
| *A. baumannii* clinical isolate 1 | 185 |
| *A. baumannii* clinical isolate 2 | 185 |
| *E. coli* CRE clinical isolate 1 | 92.5 |
| *K. pneumoniae* CRKP clinical isolate 1 | 185 |
| *E. coli mcr-1*<sup>+</sup> clinical isolate 1 | 370 |
| *E. coli mcr-1*<sup>+</sup> clinical isolate 2 | 370 |

Overall, these results suggested that synthetic peptides derived from the M6 are a potential cure for killing drug-resistant bacteria both *in vivo* and *in vitro*.

## Discussion

In this study, we described the isolation and characterization of a MCR-1 mutant with two-point mutation localized at the linker domain that specifies the disturbance of lipid homeostasis. Unlike MCR-1, this mutant renders a low level of co-resistance to β-lactam antibiotics and generates severe lipid A perturbation in the cell membrane, as indicated by a decrease in the level of LPS. This disorder eventually results in severe growth arrest, OM permeabilization, and PG remodelling. Moreover, we further identified a lipid A binding cavity that is critical for colistin resistance and bacterial membrane integrity, and the mutated cavity of M6 facilitates lipid A binding affinity, which might be responsible for the β-lactams co-resistance phenotype. Perhaps the most striking finding is that antimicrobial peptides modelled on the drug resistance gene itself are “a potential cure” to overcome drug resistance.

To date, ten variants of the *mcr* gene (*mcr-1* to *mcr-10*) have been identified in different bacteria[52, 53]. Among the MCR variants, MCR-1 is considered the largest lineage and exhibits the highest prevalence, followed by MCR-3[54, 55]. Although a two-step reaction for enzymatic hydrolysis has been proposed, the step of interplay between MCR-1 and lipid A remains unclear[17]. Recently, work in an MCR-3 homologue isolated from *Aeromonas hydrophila* suggested that a linker region consisting of 59 residues (Linker 59) guaranteed the formation of a phosphatidylethanolamine (PE) substrate-binding cavity and governed the interaction with a lipid A substrate, which was exposed to the bacterial periplasm[56]. Unlike this observation in MCR-3, the lipid A binding pocket of MCR-1 verified in our research was within the cytoplasmic membrane region. Multiple lines of evidence support our hypothesis that the lipid A binding cavity of MCR-1 is localized at the linker domain. First, our previous work demonstrated that WT MCR-1 caused OM permeabilization, resulting in bacterial fitness costs, especially in the stationary phase. The increased OM permeability was resulted from MCR-1-conferred lipid A disorder[30]. In this study, we have identified an MCR-1 variant (M6) that exhibited a set of special phenotypes, such as growth retardation, PG remodelling, and a low level of co-resistance to β-lactam antibiotics. Moreover, this variant induced a co-resistance phenotype that can be ameliorated by overexpressing LpxC, which catalyzes the first step in lipid A synthesis. This situation prompted us to hypothesize a link between membrane lipid A perturbation and phenotypic co-resistance of M6. Second, this two-points mutation localized at the linker domain but not at the catalytic domain, suggesting that the effects caused by the mutations were less possibly related with the change on catalytic process. Third, combining molecular dynamics (MD) simulation with AlphaFold structural predictions, we confirmed that a lipid A binding cavity is present at the linker domain of MCR-1. We also introduced mutations that interfered the interplay between the cavity and lipid A, and these mutants to some extent decreased colistin resistance and resumed bacterial membrane integrity. Moreover, a strong correlation between the cellular permeability of these mutations and colistin resistance was observed, which is in line with our previous finding. Based on our observations, we propose an unusual “loading-transferring” mode for this pleiotropic enzyme (Figure S14), which involves the following main steps: during the first-half reaction, the Apo state MCR-1 interacts with the PE donor at the outer leaflet of the inner membrane and accepts a PEA group in the catalytic domain, transforming MCR-1 into an active state and preparing it for interactions with LPS. Next, LPS binds with the MCR-1 loading cavity at the linker domain, which is inserted into the lipid bilayer, followed by transfer to the catalytic domain that is exposed at the interface between the inner membrane and periplasm. During the second-half reaction, the PEA group at the catalytic domain added to the 1’- or 4’-phosphate group of LPS to form PEA-LPS, which was then released back into the inner membrane, and MCR-1 switched to the Apo state again for the next cycle of LPS modification.

We do not yet understand why MCR-1 impairs cell envelope integrity. It is well established that the cytoplasmic membrane protein PbgA senses LPS in the periplasm and directs proteolytic regulation, which maintains the membrane LPS homeostasis [57]. PbgA directly interacts with its downstream regulator, the inner membrane proteins PlsY and LapB, which in turn promotes FtsH degradation of LpxC, the enzyme that catalyzes the second reaction and the first committed step in lipid A biosynthesis[58]. We found that PbgA was increased after MCR-1 expression, indicating an aberrant distribution of LPS in *E. coli*. Consistently, a reduced level of LPS was observed in MCR-1-positive cells. We therefore anticipated that MCR-1 increased bacterial permeability by reducing periplasmic LPS and further disrupting the overall LPS level, which was enhanced by the expression of M6 and resulted in PG layer remodelling (Figure 5). Consistently, the results of membrane voltage measured by Vibac2 sensor showed that the membrane voltage of M6-expressing *E. coli* was higher than that of the *mcr-1* bearing strain[47]. Given the modification mediated by MCR-1 reduces the negative potential of LPS on the OM surface, our data indicated the increase in extracellular positive potential as well as the enhanced enzymatic activity of M6. However, the molecular mechanism by which MCR-1 impacts LPS transport remains unknown.

**Figure 5.**
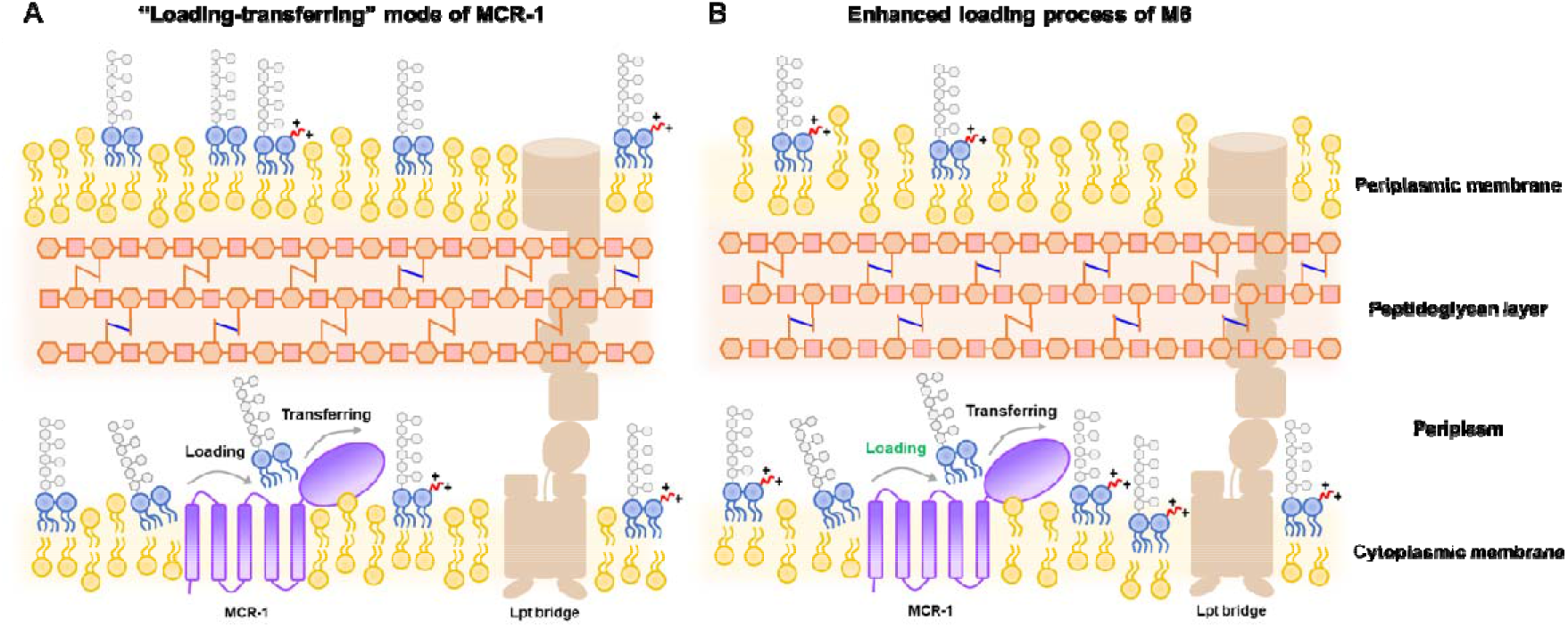
Mode of cell wall perturbation caused by MCR-1 or M6. The lipid A PEA modification mediated by MCR-1 involves the following main steps: loading of lipid A substrate in the binding cavity at the linker region; transferring to the catalytic domain to accept the PEA group; and releasing lipid A-PEA from MCR-1. (**A**) The expression of wild-type MCR-1 can modify LPS at the outer leaflet of the inner membrane, followed by transporting lipid A-PEA to the outer membrane. Although the increased positive charge at the outer membrane enables bacteria to resist colistin, it may also disturb the LPS trans-envelope transportation and cause membrane lipid A perturbation. The damaged cell wall induces remodelling of the PG layer, which increases the proportion of 3-3 cross links (blue). **(B)** The enhanced lipid A substrate loading process of M6 (highlighted in green) might promote LPS modification mediated by MCR-1, which further prevents the transportation of LPS and promotes the formation of a 3-3 cross-linking PG layer, remarkably reducing bacterial sensitivity towards β-lactam antibiotics.

There are a few caveats of our study that are worth discussing. First, our results demonstrated that severe bacterial membrane damage caused by M6 resulted in PG layer remodelling, decreasing the susceptibility of *E. coli* to β-lactam antibiotics. Deleting PBP1B (*mrcB*), a transpeptidase involved in PG cross linking, abolished the co-resistance phenotype conferred by M6. It is known that most β-lactam antibiotics target the peptidoglycan layer in the bacterial periplasm, and the mode of action of this kind of antibiotic is achieved by acting as an analogue of D-alanine at the terminal position of the tetrapeptide stem or by tightly binding to the active site of the transpeptidase (PBP) to prevent the formation of 4-3 cross links between two monomers and damage cell wall stiffness[59]. Several different groups have shown that PG layer remodelling is associated with increased OM permeability[38, 45, 60, 61]. For example, Niccolò et al. proved that the high percentage of 3-3 crosslinks in the peptidoglycan layer increased the robustness of the bacterial cell envelope in response to defect in outer membrane assembly[38]. Katharina et al. demonstrated that the deletion of all 6 LDT proteins significantly increased the SDS susceptibility of *E. coli* BW25113, emphasizing the importance of the peptidoglycan layer structure to the barrier function of bacterial cell wall[45]. Second, it should be noted that the sequence of the linker domain is diverse among the ten *mcr* variants (Figure S2) and different MCR variants have different abilities in conferring colistin resistance. For example, unlike MCR-1, MCR-9 and MCR-10 confer poor resistance to colistin[62, 63]. In fact, our data revealed that the catalytic domain is relatively conserved among ten *mcr* variants (Figure S2), which strongly suggests that the linker domain governs the diverse phenotypic polymyxin resistance of the *mcr* family. Exploring these correlations would be an interesting direction for future research. Third, it appears that MCR-1 has a pleiotropic effect. Under colistin selection, acquisition of *mcr-1* guaranteed benefit for competing among bacterial community. However, in the absence of colistin, the lipid A binding cavity of MCR-1 that disrupts lipid homeostasis confers a cost to bacterial fitness. Very recently, Pramod et al. provided evidence that *lpxC* mutations provide a fitness advantage by compensating the costs of MCR expression and that pre-existing LpxC mutations in pathogenic *E. coli* potentiate the evolution of antibiotic resistance by *mcr-1*-bearing plasmid acquisition[64]. Given the co-resistance phenotype conferred by the M6 mutant, this kind of gain-of-function mutation in the clinic is of great concern. The identification and surveillance of MCR-1 variants would also be an interesting direction of future research. Finally, although we suggest that MCR-1 or M6 expression may lead to reduction in OM LPS, disrupting lipid homeostasis and directly demonstrating LPS distribution in the IM and OM will be challenging and which require the development of new methodologies.

Over the years, the use of antibiotics now underpins many areas of medicine. Unfortunately, antibiotic treatment is accompanied with the evolution of drug resistance, resulting in poor patient outcomes. Addressing the ongoing antibiotic crisis requires the design of more effective treatment strategies that can treat drug-resistant infections. We identified an antimicrobial peptide derived from the drug resistance gene itself as “a potential cure” to kill drug-resistant bacteria. Antimicrobial peptides (AMPs) are short peptides tightly binding with the LPS on the bacterial membrane surface, which is matched with the high affinity of the lipid A binding motif of transmembrane protein for protein-substrate interactions. A recent study demonstrated that a synthetic peptide based on the lipid A binding motif of PbgA also displayed broad-spectrum activity[46]. Moreover, the introduction of an LPS binding motif into the C-terminus of temporin produces a broad spectrum of antibacterial activity[65]. Recently, AMP development has been considered an effective strategy to overcome urgent antibiotic resistance problems, and several strategies have been carried out to advance the efficiency of AMPs, including library screening, peptide structural modification and machine deep learning techniques[66, 67]. Nevertheless, our findings extend the understanding of AMPs design and provide a potential strategy for eliminating drug-resistant bacteria.

## Materials and Methods

### Bacterial strains and growth conditions

The *E. coli* strains used in this research were ATCC25922 and K-12 derivatives, including BW25113 and DH5-α. BW25113 cells were utilized to assay the influence of MCR-1/M6 (the co-resistant *mcr-1* mutant conferring reduced susceptibility to β-lactams) on bacterial drug sensitivity, viability and membrane permeability, while DH5-α cells acted as a cloning host for plasmid construction. The ATCC25922 strain was routinely used to evaluate the *in vitro* bacteriostasis activity of synthetic peptides and murine *in vivo* infection. In addition, *Pseudomonas aeruginosa* ATCC27853, *Salmonella enterica* serovar Typhimurium SL1344, *Klebsiella pneumoniae* ATCC13883 as well as *Acinetobacter baumannii* ATCC19606 were also used to test the antimicrobial effect of the 19AA-2M-tag. All the above strains were cultivated in Luria Bertani (LB) broth at 37°C, and antibiotics and promoter inducers were supplemented as follows when necessary: 100 μg/ml for ampicillin (AMP), 30 μg/ml for chloramphenicol (CHL) and 0.2% arabinose. All the strains used in this study were listed in Table S7.

### Strain construction

We generated three mutant strains (Δ*mrcB*, Δ*ycbB*, and Δ*ycfM*) as previously described[30]. Briefly, three 20-bp spacer fragments targeting the indicated gene (i.e. *mrcB*, *ycbB* or *ycfM*) were digested with *BsmB*I (Thermo Scientific) and inserted into pgRNA (Addgene #44251). To construct the donor DNA, two 500-bp homologous arms were amplified separately and fused together with fusion PCR. Electrocompetent cells containing pREDCas9 (Addgene #71541) were generated as previously described[68]. After electroporation, cultures were incubated for 3 h at 30 °C and then plated on LB agar plates containing 100 µg/ml spectinomycin, 100 µg/ml ampicillin, and 50 µg/ml bleomycin. The mutant strains were confirmed with Sanger (DNA) sequencing. All the plasmids used in this study were listed in Table S8.

### Plasmid construction

To express MCR-1 and related variants, target DNA fragments were cloned into the plasmid backbone of pACYCDuet-1. The introduction of mutations for constructing MCR-1 mutants was carried out through overlapping PCR with appropriate primers (Table S9). The process for constructing vectors expressing MCR-1 or related mutants under the regulation of the inducible arabinose promoter was as follows. Two target fragments were inserted into the multiple cloning site (MCS) region of pACYCDuet-1 through a one-step cloning strategy. The plasmid backbone pACYCDuet-1 was digested with *HindIII/PstI* (Fast digest, Thermo Fisher). The DNA fragment encoding the arabinose promoter was amplified with the primers araC-HindIII-F/ParaBAD-*mcr-1*-R, which carried 15-20 bp overlapping arms that were homologous to the 3’-end of the restricted plasmid and the 5’-end of the downstream fragment. The DNA fragment encoding WT MCR-1 or related mutants was amplified with the primers ParaBAD-*mcr-1*-F/Terminator-PstI-R, which carried 15-20 bp overlapping arms homologous to the 3’-end of the upstream fragment and 5’-end of the restricted plasmid. The purified plasmid and target fragments were mixed in a molar ratio of 1:3, followed by adding 2× ClonExpress^®^ Mix (ClonExpress Ultra One Step Cloning Kit, Vazyme) into the reaction and incubating in a thermocycler at 50 °C for 15 minutes. Finally, the well-reacted products were transformed into *E. coli* DH5-α competent cells for enrichment. All target constructs were confirmed by Sanger sequencing. The well-constructed plasmids were transformed into *E. coli* BW25113 cells through electroporation. All the plasmids and primers used in this study were listed in Table S8 and S9.

### Homology protein modelling

To analyse the protein structure of MCR-1 and related variants, homology models for MCR-1, M6, P188A and P195S were generated by SWISS-MODEL based on the crystal structure of *Neisseria meningitides* lipid A phosphoethanolamine (PEA) transferase (PDB entry: 5FGN). The PDB files of target proteins were visualized by PyMOL version 2.6 software.

Furthermore, in this study, either truncation or mutations on the hinge-like linker, which connects the TM domain with the catalytic domain in any certain MCR member, were found to be sensitive to the enzymatic action of MCR-1. This is consistent with previous findings for MCR-3, specifically, the linker domain determines the enzymatic action [56]. This indicates that the area of the hinge-like linker could be a candidate binding site for lipid A.

To predict the binding mode of Lipid A to MCR-1, a workflow was designed, as shown in Figure S15A. Small molecules always bind with proteins in their lowest energy binding conformations. Hence, we extracted a conformation of Lipid A from RCSB (https://www.rcsb.org/) (PDB code: 5IJD) for our molecular docking simulation. In the experimental complex structure, lipid A is in complex with TLR4/MD-2. Interestingly, we found that despite interacting with different proteins, the lipid A analogues, i.e., Eritoran and Palmitoyllpid A, present similar molecular conformations with the extracted lipid A conformation. Structural analysis of lipid A and its analogues revealed that the lipid tails bunched together in the centre (Figure S15B). This further confirms that the conformation of lipid A obtained for docking is a reasonable initial conformation for our docking. The structure of MCR-1 was downloaded from AlphaFold DB, which is a protein database that is computationally predicted and uses the deep learning-based protein structure prediction method Alphafold[69]. SiteMap of Maestro[70] was used to search on the surface of MCR-1 to identify the potential binding sites for any ligand. As a result, a large potential binding site located around the area of the hinge-like linker was discovered with a Site Score as high as 0.964 (Figure S15C). Then, the coordinates of the residues of this site, i.e., I168, F183, R190, I196 and K204, were used to determine the centre of the binding pocket. An inner grid with a size of 30×30×30 Å and a larger outer box with a size of 35×35×35 Å were set. Then, the mode of SP in Glide of Maestro[71] was chosen to perform docking simulation. Both the ligand and protein were taken as rigid. The radius of van der Waals, initial pose, and cut-off of the pose energy filter were set as 0.5, 10000 and 500 kcal/mol, respectively. The top-ranked binding pose was further structurally optimized by using Prime. This structure was used as the starting structure for molecular dynamics simulation to obtain a more reliable binding pose of Lipid A against MCR-1. Six independent simulations (noted as A-F) with a length of 200 ns were performed by using Desmond[72] of Schrödinger2021-2 under the OPLS4 force field. The POPC membrane was assembled to MCR-1 by referencing the positions of the membrane in the homologue protein EptA. The complexed structures were explicitly solvated with TIP3P water molecules under cubic periodic boundary conditions for a 15 Å buffer region. The overlapping water molecules were deleted, and 0.15 M NaCl were added, and the systems were neutralized by adding Na^+^ as counter ions. Brownian motion simulation was used to relax these systems into local energy minimum states separately. An ensemble (NPT) was then applied to maintain the constant temperature (310 JK) and pressure (1. 01325 bar) of the systems, and the simulations start with two random initial velocities. The produced trajectories were clustered by all atoms of the MCR1 protein using the Desmond trajectory cluster analysis module and produced five clusters per trajectory. Then, the clustered conformers from all six trajectories were further clustered by heavy atoms of lipid A using Maestro13.2. Finally, the largest cluster was identified as the representative conformation of MCR1-lipid A. The root-mean-square deviations (RMSDs) of the heavy atoms in all simulated trajectories were calculated by Maestro13.2, and the RMSD plot was plotted by Origin version 2021 (Figure S15D).

To assess the impact of mutations of MCR-1 on their binding strength with lipid A, M6 were designed and compared with the WT of MCR-1. The initial binding complex of lipid A and mutant MCR-1 was constructed via the Residue Scanning tool of Maestro based on WT MCR-1. The residue conformations were correspondingly optimized after mutations. The generated binding complex of the mutant with lipid A was taken as the input structure of MD simulations for further structure optimization. The membrane setting was the same as the above simulations, and the parameter settings were also the same as the simulations for the WT complex.

### Growth curve measurements

Fresh single colonies of *E. coli* BW25113 carrying MCR-1, M6 or empty vector were inoculated into fresh Luria Bertani (LB) broth containing 30 μg/ml CHL and cultivated overnight at 37 °C. The overnight cultures were then adjusted to an OD_600_ ranging from 0.5 to 0.6 with saline and diluted at a ratio of 1:10. The 20 µl diluted cultures were inoculated into 180 µl LB broth containing 0.2% arabinose and 30 µg/ml CHL in a 96-well plate. Three replicates were carried out for each strain. A nanophotometer (NP80, IMPLEN) was utilized to measure the optical density at 600 nm (OD_600_) in each well per hour. Growth curves were plotted with Prism 9 software.

### Agar dilution MIC tests

To evaluate the antibiotic susceptibility of *E. coli* BW25113 carrying MCR-1, M6 or empty plasmid, an MIC assay was performed by the agar dilution method, which was conducted as previously established[73]. To prepare agar plates that contained antibiotics in a series of concentrations, antibiotics were added to Mueller–Hinton agar (MHA) containing 30 µg/ml chloramphenicol (CHL) and 0.2% arabinose to maintain the existence of target plasmids and induce the expression of MCR-1 or M6. Fresh single colonies of the above target strains were inoculated into fresh Mueller–Hinton broth (MHB) containing 30 µg/ml CHL and cultivated overnight at 37 °C. The overnight cultures were then adjusted with saline to an OD_600_ that ranged from 0.5 to 0.6. The well-adjusted cultures were diluted in six gradients, ranging from 10^-1^ to 10^-6^, with saline. For each dilution, 3 µl of culture was spotted onto MHA plates containing antibiotics and incubated at 37 °C overnight. Growth was determined by counting colony-forming units (CFUs). Three replicates were carried out for each strain. The MIC values were defined as the concentration at which bacterial growth was absolutely inhibited. The following antibiotics and concentrations (highest to lowest) were utilized for the assay: ampicillin (AMP, 2 to 32 μg/ml), cefoxitin (FOX, 2 to 10 μg/ml), imipenem (IMP, 0.5 to 2.5 μg/ml), ceftazidime (CAZ, 0.2 to 0.6 μg/ml), cefotaxime (CTX, 0.1 to 0.5 μg/ml) and meropenem (MEM, 0.01 to 0.1 μg/ml). The results were plotted with Prism 9 software.

### Determination of the outer membrane integrity

A 1-*N*-phenylnaphthylamine (NPN) uptake assay was performed to determine the outer membrane integrity of the following strains: *E. coli* BW25113 MCR-1/M6, BW25113 Δ*mrcB*/Δ*mrcB* M6 and BW25113 MCR-1/M6 carrying pBAD24-LpxC. NPN is a kind of probe that emits strong fluorescent signals in phospholipid environments, thus, NPN can verify the permeability of the outer membrane for Gram-negative bacteria[74]. The procedures were performed as previously described[75]. Briefly, fresh single colonies of target strains were inoculated into fresh LB broth containing 30 µg/ml CHL and cultivated overnight at 37 °C. The overnight cultures were then adjusted to an OD_600_ ranging from 0.5 to 0.6 with saline and subcultured into 1 ml of LB broth containing 30 µg/ml CHL and 0.2% arabinose with shaking at 37 °C until the cultures reached an OD_600_ = 0.5. The cells were harvested by centrifugation (4,000 rpm for 3 min), washed twice with assay buffer (5 mM HEPES, 5 mM glucose, pH=7.2) and resuspended in assay buffer to a final OD_600_ = 1. Then, 100 μl of washed cultures and 100 μl of assay buffer containing 20 μM NPN were mixed and added to a 96-well half area black opaque plate (Greiner Bio), in which the washed culture that was replaced with saline was set as a control to remove the background signal. The fluorescence for each well was immediately monitored with a microplate reader (BioTek) at an excitation wavelength of 350 nm and emission wavelength of 420 nm.

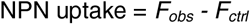

*F_obs_* represents the NPN uptake of different strains, and *F_ctrl_* represents the background signal without the addition of bacterial culture. Three replicates were carried out for each strain. The results were plotted with Prism 9 software.

### Determination of the inner membrane integrity

A PI (propidium iodide) staining assay was performed to determine the inner membrane integrity of the following strains: *E. coli* BW25113 MCR-1/M6, BW25113 Δ*mrcB/*Δ*mrcB* M6 and BW25113 MCR-1/M6 carrying pBAD24m-LpxC. Acting as a dye crossing compromised bacterial membranes and binding with DNA and RNA inside of damaged cells[76], PI was utilized to identify dead cells or those with irreversibly damaged membranes. Therefore, the uptake of PI can reflect the permeability of the bacterial inner membrane. The process for PI staining was performed as follows: fresh single colonies of the above target strains were inoculated into fresh LB broth containing 30 µg/ml CHL and cultivated overnight at 37 °C. The overnight cultures were then adjusted to an OD_600_ ranging from 0.5 to 0.6 with saline and were subcultured into 2 ml of LB broth containing 30 μg/ml CHL and 0.2% arabinose with shaking at 37 °C, and the samples were collected at 4 hr and 8 hr after induction. Cultures were harvested by centrifugation (4,000 rpm for 3 min), washed twice with saline and resuspended in 95 µl buffer A (PI staining kit, Sangon Biotech). Then, 5 µl of PI dye was added to each resuspended culture and incubated in the dark for 15 min. The percentage of the PI-positive population was verified by a flow cytometer (Gallios10, Beckman). The sample without staining with PI was set as a control for gating. The data were analysed with FlowJo version 10 software and visualized with Prism 9 software.

### Screening of the MCR-1 mutant library against a variety of antibiotics

The mutant library of MCR-1 was applied for screening on LB agar plates containing antibiotics as shown in Table S1, in which the BW25113 carrying MCR-1 or empty vector were set as controls. The main procedures are shown in Figure S1A. In brief, the frozen stocks of the MCR-1 mutation library, BW25113 carrying MCR-1 or empty vector control were thawed at room temperature and inoculated into 10 ml of fresh LB broth containing 30 μg/ml CHL, followed by cultivation at 37 °C until the cultures reached an OD_600_ = 0.5. The logarithmic-phase cultures were then induced with 0.2% arabinose for 2 hr. To ensure that all the mutation genotypes in the MCR-1 mutant library were present on each plate at least once, the coupon collector problem was applied as follows[77]:

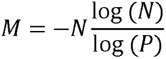

where *M* represents the number of cells on each plate, *N* is the number of total mutation genotypes and *P* is the probability of coverage for all the genotypes.

For P = 99.9%:

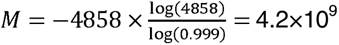

Therefore, approximately 1×10^11^ cells were plated on each agar plate to achieve 99.9% coverage of all the single-point mutation genotypes. After incubation at 37 °C for 16 hr, CFUs were counted to evaluate viability.

### Time killing assay to assess colistin susceptibility

Time killing assays were performed to evaluate the sensitivity of the BW25113 harbouring MCR-1/M6/empty vector towards colistin (CT). Similar to the process mentioned in previous research[78], overnight cultures of the above target strains were diluted with LB broth to a density of ∼10^6^ CFUs/ml and aliquoted for treatment with antibiotics as follows: CT (4, 8, 16 and 32 μg/ml), and the samples without drug treatment were set as controls. For each treatment, 100 µl of culture was sampled at time intervals of 0 hr (T0) and 2 hr (Tn) and diluted in six gradients, ranging from 10^-1^ to 10^-6^, with saline. Then, 100 μl of diluted culture was plated on LB agar plate. Each treatment was repeated in triplicate. After incubation at 37 °C for 18 hr, CFUs were counted to evaluate the antimicrobial effect.

### Morphological analysis by scanning electron microscopy (SEM) to evaluate the bacterial cell wall surface

Scanning electron microscopy was performed to observe the change in bacterial surface. The main process was similar to that previously described[79]. For BW25113 expressing MCR-1 or M6, exponential phase cultures were collected, and BW25113 carrying the empty plasmid vector was set as a control; for ATCC25922 treated with 24AA-2M / 19AA-2M-tag, cultures were grown to exponential phase and then treated with 92.5μM synthetic peptides for 1 hr, and ATCC25922 treated with DMSO was set as a control. All the samples were harvested by centrifugation (4,000 rpm for 3 min) and fixed with 2% glutaraldehyde (Servicebio) for at least 4 hr at room temperature, followed by dehydration with a graded ethanol series and air drying. The dried powder was loaded on a rotating stage, sputtered with gold using a vacuum coater (Leica EM ACE200) and coated with a 0.1 mm gold layer. Microscopy was performed with a desktop field emission scanning electron microscope (Phenom Pharos G2, Thermo Fisher Scientific). Images were collected using a secondary electron detector, and the acceleration voltage was adjusted to 10 kV. SEM images were recorded at magnifications ranging from 20,000× to 100,000×.

### Morphological analysis by transmission electron microscopy (TEM) to evaluate cell wall morphology

Transmission electron microscopy was performed to explore the cell wall morphological change in MCR-1-expressing cells or M6-expressing cells. Exponential phase cultures were collected and harvested by centrifugation (4,000 rpm for 3 min), followed by fixation with 2% glutaraldehyde (Servicebio) for at least 4 hr at room temperature. The sample pellets were embedded in a 1 mm cube containing 4% low melting point agarose and postfixed with osmium tetroxide for 2 hr. The specimens were then dehydrated in a graded series of ethanol, transferred to propylene oxide and embedded in Epon according to standard procedures. Thin (80 nm) sections were cut and collected on copper grids. After staining with uranyl acetate and lead citrate, the specimens were subsequently examined with a field emission transmission electron microscope (JEM-2100, JEOL) operated at an accelerating voltage of 80 kV. TEM images were recorded at magnifications of 20,000× to 80,000×.

### Quantitative real-time PCR to assess transcription of envelope and transpeptidase genes

To verify the change in the transcriptional level of genes related to envelope responses and LDTs (L, D-transpeptidases), mRNA was extracted for quantitative real-time PCR (q-PCR). Overnight cultures of target strains were sub-cultured into 1 ml fresh LB broth containing 30 µg/ml CHL and 0.2% arabinose. After induction for 2 hr, exponential phase cultures were collected and harvested by centrifugation (4,000 rpm for 3 min). After removing the supernatant, the cell pellet was resuspended in RNA-easy Isolation Reagent (Vazyme). Total RNA was precipitated by adding isopropanol and was collected by centrifugation (37 °C, 12,000 rpm for 10 min). The supernatant was discarded, and the mRNA pellet was washed with 75% ethanol. The mRNA pellet was dehydrated through air drying and dissolved in RNase-free H_2_O. The contaminating genomic DNA was digested with gDNA wiper Mix (Vazyme). cDNA was prepared from purified mRNA with HiScriptII qRT SuperMix II (Vazyme) through reverse transcription. The cDNA levels of target genes were then quantified by quantitative real-time PCR (qRTJPCR) on a CFX96 cycler (BIO RAD) by using AceQ Universal SYBR qPCR Master Mix (Vazyme) according to the manufacturers’ protocol. All qPCR primers were determined to be >95% efficient, and the cDNA masses tested were experimentally validated to be within the linear dynamic range of the assay. Signals were normalized to those of the transcript of the housekeeping gene *rpoB* and quantified with ΔΔCT analysis.

### Expression level quantification of PbgA and LPS

To verify the influence of MCR-1/M6 expression on bacterial LPS homeostasis, immunoblot analysis was performed to evaluate the expression levels of PbgA (a periplasmic lipid A sensor, labelled with HA-tag) and LPS. Briefly, fresh single colonies of target strains were inoculated into 5 ml fresh Luria Bertani (LB) broth containing 30 µg/ml CHL and 0.2% arabinose. After induction for 2 hr, exponential phase cultures were collected and harvested by centrifugation (4,000 rpm for 3 min). Then, 1 ml RIPA lysis buffer (Beyotime) was added to resuspend the cell pellet, and the bacillus was further broken down with an ultrasonic processor (SONICS, VCX 130). After centrifugation (10,000 rpm for 1 min), the supernatant of the lysate was stored, and the protein concentration of each sample was quantified by using the Bradford Protein Assay Kit (Beyotime) according to the manufacturer’s protocol. The samples were standardized by protein concentration, mixed with SDSJPAGE Sample Loading Buffer (Beyotime), and electrophoresed on a 12% Bis-Tris SDS-polyacrylamide gel. After electrophoresis, proteins and LPS were transferred to a polyvinylidene difluoride (PVDF) membrane (Thermo Fisher) using the Trans-Blot Turbo Transfer System (BIO RAD). Primary antibodies against RpoB (BioLegend), LPS core (Hycult Biotech) and HA-tag (Cell Signalling) were used at dilutions of 1:100,000, 1:1,000 and 1:10,000, respectively. Goat anti-rabbit horseradish peroxidase (HRP) conjugate (Zen Bioscience) and rabbit anti-mouse HRP conjugate (Dingguo Biology) secondary antibodies were each used at a 1:10,000 dilution. After processing of chemiluminescent detection with a ChemiDoc Touch Imaging System (BIO RAD), the protein levels were quantified with Fuji software, in which the expression level of RpoB was regarded as a reference. Three replicates were performed for each sample.

### Fluorescent imaging to assess membrane shrinkage

Since membrane shrinkage of *E. coli* was accompanied by the formation of inner membrane foci[39], we labelled the periplasm with super-folded GFP (sfGFP) and the cytoplasm with mCherry to construct a two-fluorescent reporter system. For each strain, stationary phase culture was collected after induction with 0.2% arabinose, and 5 μl of bacterial culture was placed on a gel pad containing 1% agarose and covered with a coverslip. The GFP and mCherry signals were recorded with a fluorescence microscope (Olympus BX63) on phase contrast equipped with a 100× oil immersion objective and a xenon lamp. The images merged with the two fluorescent signals were processed with Fuji software.

### Fluorescent imaging to assess membrane voltage

The construction of genetically-encoded membrane voltage sensor Vibac2 was as described in previous research[47]. ViBac2 is a double-channel fusion protein that emits green and red fluorescence. The fluorescence intensity of GFP (Ex=488 nm, Em=512 nm) responds to membrane voltage, and the mCherry (Ex=561 nm, Em=610) is used to normalize protein expression. Thus, the fluorescence ratio indicates the relative membrane voltage in *E. coli* cells. The plasmids encoding MCR-1 or M6 were transformed into *E. coli* BW25113 competent cell expressing Vibac2. Strain harbouring pACYCDuet-1 empty plasmid was set as control. For each strain, exponential phase culture was collected after induction with 0.2% arabinose, and 5 μl of bacterial culture was placed on a gel pad containing 1% agarose and covered with a coverslip. The GFP and mCherry signals were recorded with a fluorescence microscope (Olympus BX63) on phase contrast equipped with a 100× oil immersion objective and a xenon lamp. The images merged with the two fluorescent signals were processed with Fuji software.

### Label-free quantitative proteome analysis

To profile the proteomic characteristics of the M6-positive strain, a label-free quantitative proteome analysis was performed to compare the differentially expressed proteins between BW25113-carrying MCR-1 and BW25113-carrying M6. The process for sample preparation was similar to that mentioned previously[80]. In brief, fresh single colonies of target strains were inoculated into fresh Luria Bertani (LB) broth containing 30 μg/ml CHL and 0.2% arabinose, and three replicates were carried out for each strain. After induction for 2 hr, exponential phase cultures were collected and harvested by centrifugation (4,000 rpm for 3 min). The cell pellets were sent to APTBIO Co. (Shanghai, China) for proteomic analysis. A volcano map was constructed with R studio (version 3.6.1) to show the comprehensive changes induced by the expression of M6 in *E. coli* compared with the *mcr-1*^+^ strain. The differentially expressed proteins were mapped to the KEGG database (http://geneontology.org/) and Gene Ontology (GO) terms (with Blast2GO software) for enrichment analysis, and the results were plotted by R studio (version 3.6.1). Additionally, the studied proteins were subjected to proteinJprotein interaction (PPI) analysis through STRING (http://string-db.org/), and the results were plotted by Cytoscape software.

### SDS sensitivity assay to evaluate cell wall permeability

The sodium dodecyl sulfate (SDS)–ethylenediaminetetraacetic acid (EDTA) sensitivity assay was performed with an agar dilution strategy to evaluate the cell wall permeability. To prepare agar plates containing SDS in series concentrations, SDS was added to Mueller–Hinton agar (MHA) containing 30 µg/ml chloramphenicol (CHL) and 0.2% arabinose. EDTA was also added to the agar to a final concentration of 100 µM to increase the permeability of the bacterial cell wall. Fresh single colonies of the above target strains were inoculated into fresh Mueller–Hinton broth (MHB) containing 30 µg/ml CHL and were cultivated overnight at 37 °C. The overnight cultures were then adjusted to an OD_600_ ranging from 0.5 to 0.6 with saline. The well-adjusted cultures were diluted with saline in six gradients, ranging from 10^-1^ to 10^-6^. For each dilution, 3 µl of culture was spotted onto MHA plates containing SDS-EDTA and were incubated at 37 °C overnight. Growth was determined by counting CFUs. Three replicates were carried out for each strain. The inhibition values were defined as the concentration at which bacterial growth was absolutely inhibited.

### Biolayer interferometry

An *in vitro* interaction assay was carried out to verify the affinity between lipid A and synthetic peptides. Both Kdo_2_-lipid A (Avanti Polar Lipids) and biotinylated peptide (GenScript) stock powders were dissolved in DMSO and diluted with PBST. All assays were performed at 25 °C in PBST buffer containing 10% DMSO. The biotin-blocked reference streptavidin (SA) biosensor biosensors (Sartorius) were soaked with PBST+10% DMSO buffer for 10 min, and biotinylated peptides were loaded onto SA biosensors to a response of approximately 0.5 nm. Target peptides were bound to Kdo_2_-lipid A in series of concentrations (150, 100, 50, 25 and 10 μM) with 300 s association and dissociation steps. Assays were performed in triplicate on an Octet Red384 (Sartorius). Dissociation constants for 16AA-WT and 16AA-M6 interactions with Kdo_2_-lipidA were estimated by plotting response values at equilibrium as a function of concentration and fit to a global specific binding with the Hill slope model in Prism 9.

### Growth inhibition assay

To verify the bacteriostatic activity of the synthetic peptides towards *E. coli* ATCC25922, a growth inhibition assay was performed. The density of the target strain overnight culture was adjusted with saline to OD_600_=0.5 and further diluted at a ratio of 1:10. The diluted culture was aliquoted for treatment with synthetic peptides as follows: 24AA-WT, 24AA-2M and 19AA-2M-tag. All the peptides were in concentrations of 370, 185 and 92.5 μM. The samples without drug treatment were used as controls. For each treatment, 100 µl of culture was sampled 0, 3, 6 and 24 hr after the addition of target peptides and diluted with saline in six gradients, ranging from 10^-1^ to 10^-6^. Then, 3 μl of each dilution was spotted onto LB agar plates. Each treatment was repeated in triplicate. After incubation at 37 °C for 18 hr, CFUs were counted to evaluate the antimicrobial effect. All the synthetic peptides used in this study were listed in Table S10.

### Checkerboard assay

Synergy measurement by checkerboard analysis was utilized to determine the interactions of colistin (CT) and 19AA-2M-tag on *E. coli* ATCC25922 inhibition in comparison to their individual effects alone. The checkerboard assay was set up in a 96-well plate. Briefly, Columns 2 to 12 contained 2-fold serial dilutions of colistin (ranging from 10 to 0.0098 μg/ml), and Rows A to G contained 2-fold serial dilutions of 19AA-2M-tag (ranging from 92.5 to 1.45 μM). Column 1 contained a serial dilution of 19AA-2M-tag alone (ranging from 92.5 to 1.45 μM), while Row H contained a serial dilution of colistin alone (ranging from 10 to 0.0098 μg/ml). These two groups were used as controls to determine the MIC value for colistin or 19AA-2M-tag of the target strain. *E. coli* ATCC25922 was cultured to exponential phase and adjusted to a density of OD_600_=0.5 by diluting with LB broth. After dilution at a ratio of 1:10, 20 µl of diluted culture was added to 180 µl of LB broth containing colistin and/or 19AA-2M-tag in a 96-well plate. A nanophotometer (NP80, IMPLEN) was utilized to measure the optical density at 600 nm (OD_600_) in each well before and after incubation at 37 °C for 16 hr. The fractional inhibitory concentration index (FICI) was calculated as follows:

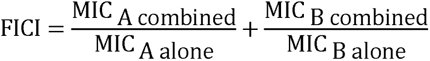

where MIC _A combined_ and MIC _B combined_ are the MICs of each drug in combination, while MIC _A alone_ and MIC _B alone_ are the MICs of each drug individually.

### Murine red blood cell lysis assay

To verify the cytotoxicity of 24AA-2M and 19AA-2M-tag towards human red blood cells, the lactate dehydrogenase (LDH) assay was utilized to measure cell permeation. Each peptide was diluted with saline to a final concentration of 185 µM. Mouse blood samples were added to a 96-well plate (3000 cells/well), and 100 µl of diluted peptide was added in triplicate to different wells of the plate. The LDH-based TOX-7 kit (SigmaJAldrich) was used for quantification of LDH release from the cells. The results represent the mean values from triplicate measurements and were plotted by Prism 9.

### Murine infection model

The drug combination of colistin and 19AA-2M-tag was used to test the efficiency against *in vivo* infection. Ten-week-old C57BL/6 mice were infected with 1×10^7^ CFUs of *E. coli* ATCC25922 or CRE through intraperitoneal injection. Two hours after infection 200 μg 19AA-2M-tag was supplemented to the infected mouse by intraperitoneal injection as treatment. The individuals without treatment were set as controls, and each condition was carried out in triplicate. Mice were sacrificed 24 hr after infection, and bacterial burdens in the liver and spleen were determined by spotting serial dilutions of tissue homogenates on LB plates. After incubation at 37 °C for 18 hr, CFUs were counted to evaluate the antimicrobial effect.

### Statistical analysis

Statistical analysis was performed using Prism (version 9, GraphPad Software). Data were analysed using the paired Student’s t test, and in the comparisons of data from three or more conditions, analysis of variance (ANOVA) was used. A p value of 0.05 or less was considered statistically significant.

## Acknowledgment

This work was supported by the National Natural Science Foundation of China (grant number 81830103, 82061128001 to G.-B T., grant number 82002173, 82272378 to S. F.), Guangdong Natural Science Foundation (grant number 2017A030306012 to G.-B T.), and Project of High-level Health Teams of Zhuhai in 2018 (The Innovation Team for Antimicrobial Resistance and Clinical Infection to G.-B T.). The plasmid encoding bacterial membrane voltage sensor Vibac2 was a generous gift from Prof. Bai Fan’s lab.

## Author Contributions

LJL, LLZ, and LW contributed equally to this study. GBT, SYF, and FB conceived the idea, designed and supervised the study. LJL, SYF, LW, DRZ, YC and LXL conducted the experiments and produced the tables and figures. YXL, WFL, WJW, CCZ, NS, and HZ searched the literature. JCL, YD and DLP contributed to data analysis and interpretation. LJL, SYF, LW, FB and GBT wrote the manuscript. All authors reviewed, revised, and approved the final report.

## Supplementary figures

**Figure S1.**
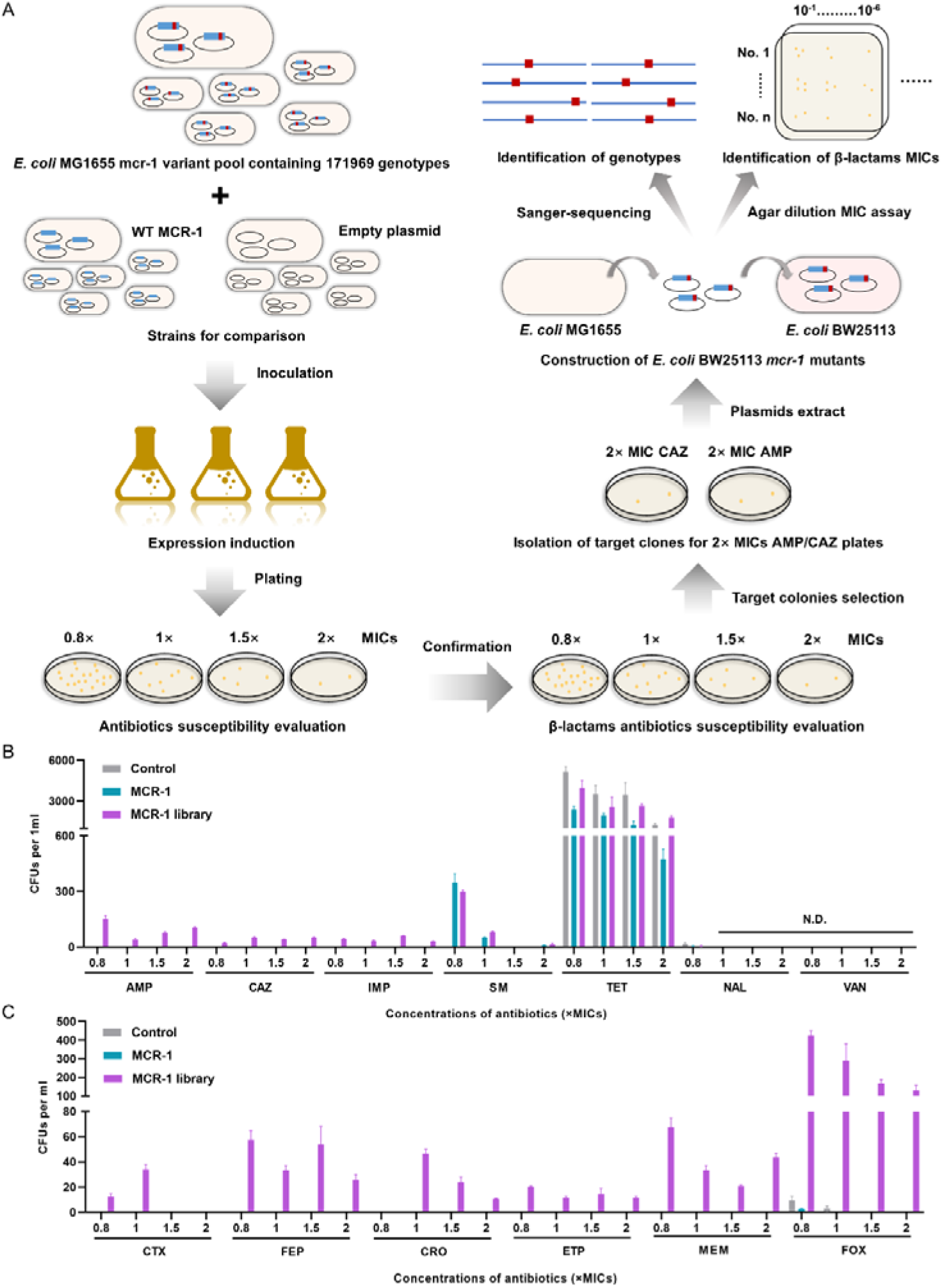
Identification of isolates with reduced sensitivity towards β-lactam antibiotics from the MCR-1 mutant library. **(A)** The schematic illustration shows the process of screening MCR-1 mutants with reduced susceptibility towards β-lactams. The variant library containing 171969 genotypes was cloned into the medium-copy plasmid pACYDuet-1 to create the BW25113 strain pool. Along with *E. coli* BW25113, strains carrying WT MCR-1 or empty plasmid were applied for testing sensitivities towards different antibiotics. Logarithmic-phase cultures of the three target strains were first induced with 0.2% arabinose for 2 hr, followed by plating on LB agar plates containing AMP, CAZ, IMP, SM, TET, NAL or VAN. All antibiotics were at MICs of 0.8×, 1×, 1.5× and 2×. After incubation at 37 °C for 16 hr, the number of surviving bacilli was counted to evaluate antibiotic susceptibility **(B)**. It appears that certain MCR-1 variants exhibited reduced sensitivity to β-lactam antibiotics. To confirm the sensitivity of β-lactam antibiotics among the three strains, a similar process was performed, and well-induced cultures were plated on LB agar plates containing CTX, FEP, CRO, ETP, MEM or FOX. CFUs were counted after incubation at 37 °C for 16 hr **(C)**. Fifty colonies of the *mcr-1* library in the background of *E. coli* BW25113 were selected from the plates containing CAZ or AMP at concentrations of 2× MICs. For each isolate, the pACYCDuet-1 plasmid carrying *mcr-1* variants was extracted, followed by transformation into *E. coli* BW25113 electroporation competent cells to eliminate influence caused by chromosomal mutation. Next, reconstructed strains were subjected to agar dilution MICs test to verify susceptibility to CAZ, AMP or FOX. In addition, the *mcr-1* genotypes of the target isolates were verified through Sanger sequencing. Both Panels **(B)** and **(C)** were visualized with Prism 9 software. All the above-described experiments were performed three times with similar results. Error bars indicate standard errors of the means (SEMs) for three biological replicates.

**Figure S2.**
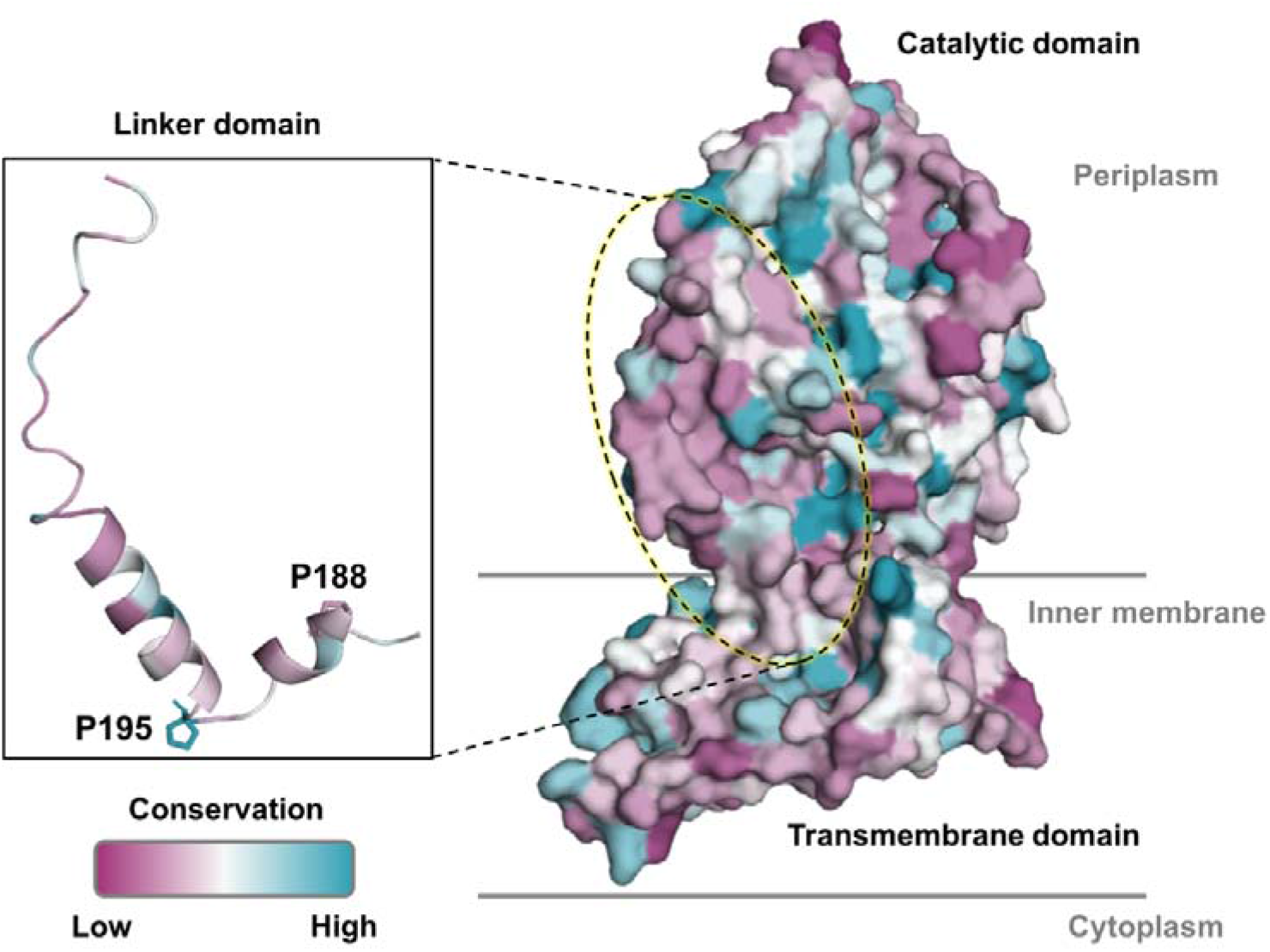
Conservation of the protein structure of the MCR family. A surface representation shows the conservation analysis calculated across 106 MCR family proteins, in which red represents low homologous positions and green represents high conservation. The linker region and the mutated residues are shown in cartoon and sticky representations, respectively. The protein structure was analysed and plotted by PyMOL 2.6 software.

**Figure S3.**
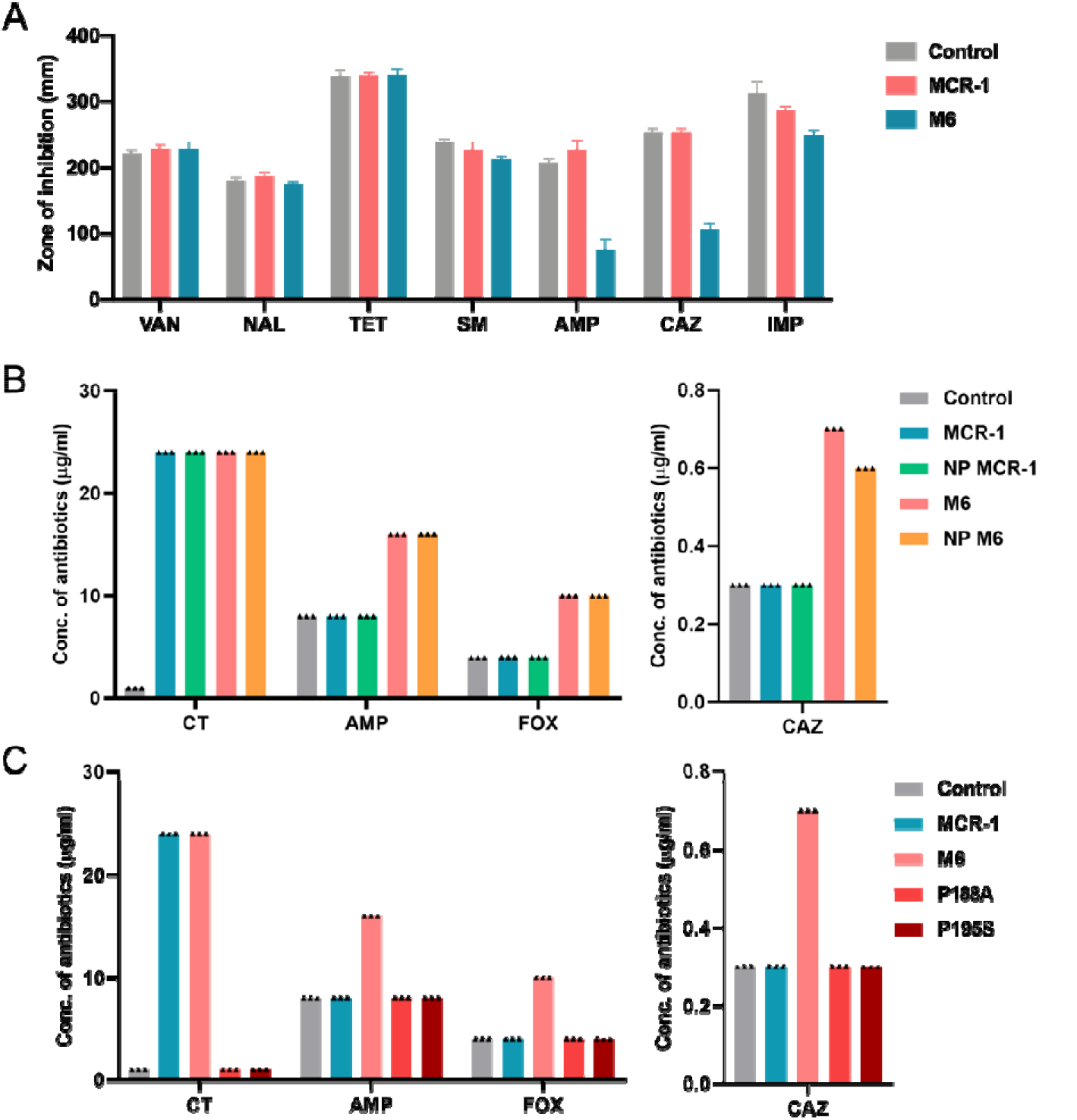
Verification of the β-lactam antibiotic co-resistance rendered by M6. **(A)** Zone of inhibition of *E. coli* BW25113 harbouring empty plasmid, MCR-1 or M6 generated by disk diffusion method. **(B)** Evaluation of the impact of different levels of MCR-1 expression on the co-resistance phenotype. pACYCDuet-1 carrying MCR-1 or M6 under the regulation of the MCR-1 native promoter (NP MCR-1 or NP M6) was generated, and a similar approach was applied to evaluate the sensitivity towards CT and β-lactam antibiotics. Each triangle represents an independent experiment. The experiments were performed three times with similar results. NP, native promoter. **(C)** Different levels of antibiotic sensitivity in *E. coli* expressing either wild-type MCR-1, M6 or its single point mutants.

**Figure S4.**
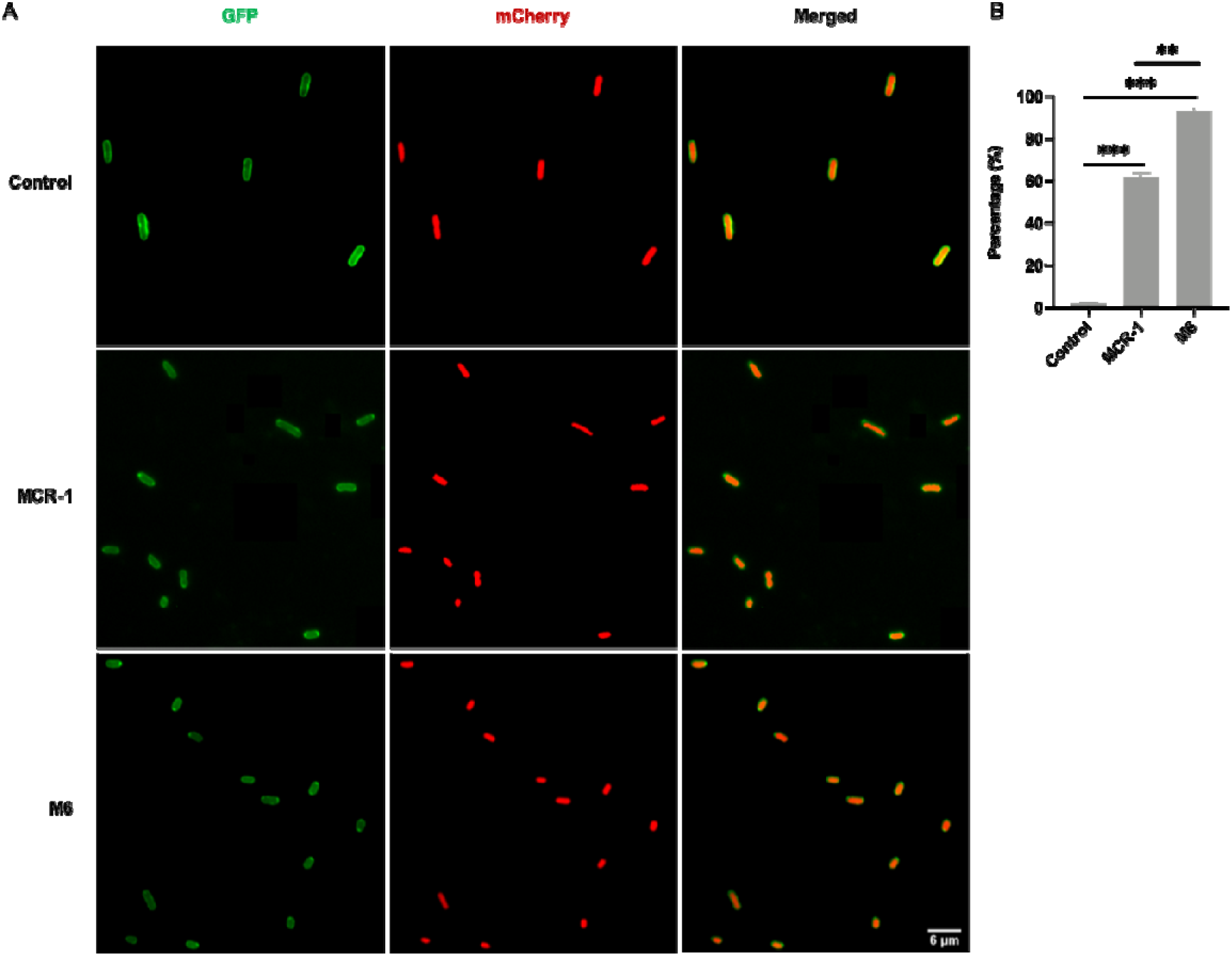
Expression of M6 causes *E. coli* membrane shrinkage. **(A)** Representative periplasmic GFP and cytoplasmic mCherry images of BW25113 cells expressing MCR-1 or M6 during stationary stage. Shrinkage of the cytoplasm was evident by the bright periplasmic GFP signal. (B) The percentage of cells exhibiting shrinkage at pole(s) were calculated. The graph was visualized with Prism 9 software. All the above-described experiments were performed three times with similar results. Error bars indicate standard errors of the means (SEMs) for three biological replicates. A two-tailed unpaired *t* test was performed to determine the statistical significance of the data. **, *P*< 0.01; ***, *P*< 0.001.

**Figure S5.**
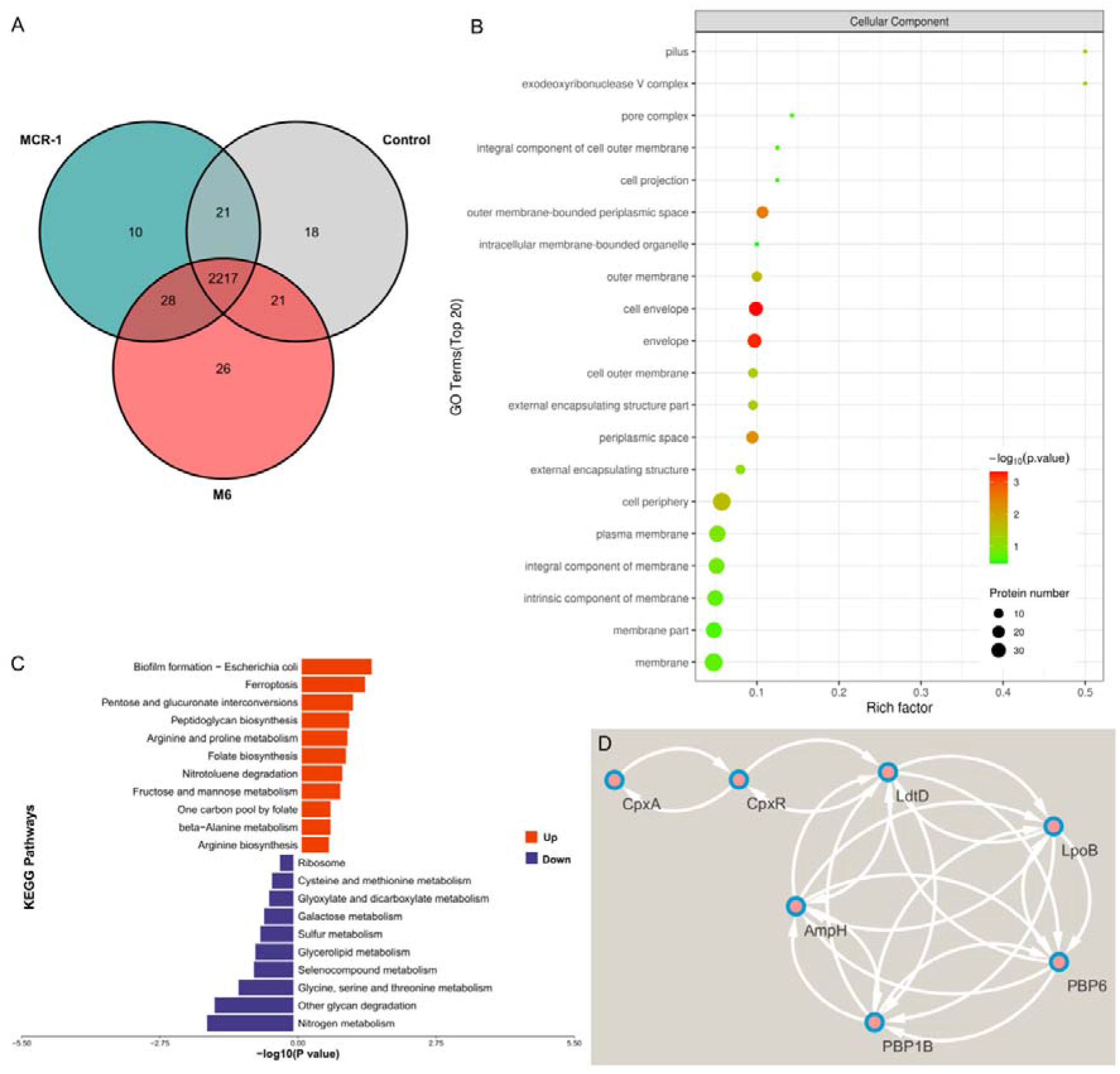
Comprehensive proteome profiles of M6 using label-free quantitative proteomics. **(A)** Venn diagram representing the differentially expressed proteins of BW25113 that harboured empty vector (control), MCR-1 and M6 during the exponential phase with the induction of 0.2% arabinose. **(B)** Top 20 pathways according to GO enrichment. The size and colour of the points represent the number of target proteins and the q-values, respectively. The rich factor showed the enrichment degree in the GO pathway. **(C)** Visualization of the top KEGG enrichment pathway in M6-expressing cells compared with MCR-1-expressing cells. The upregulated pathway is labelled in red, whereas the downregulated pathway is labelled in blue. **(D)** Protein–protein interaction (PPI) analysis of the peptidoglycan layer remodelling pathway in M6 based on proteomics profiles, which were analysed by STRING and visualized with Cytoscape software. Two biological replicates for each strain were used for the analysis.

**Figure S6.**
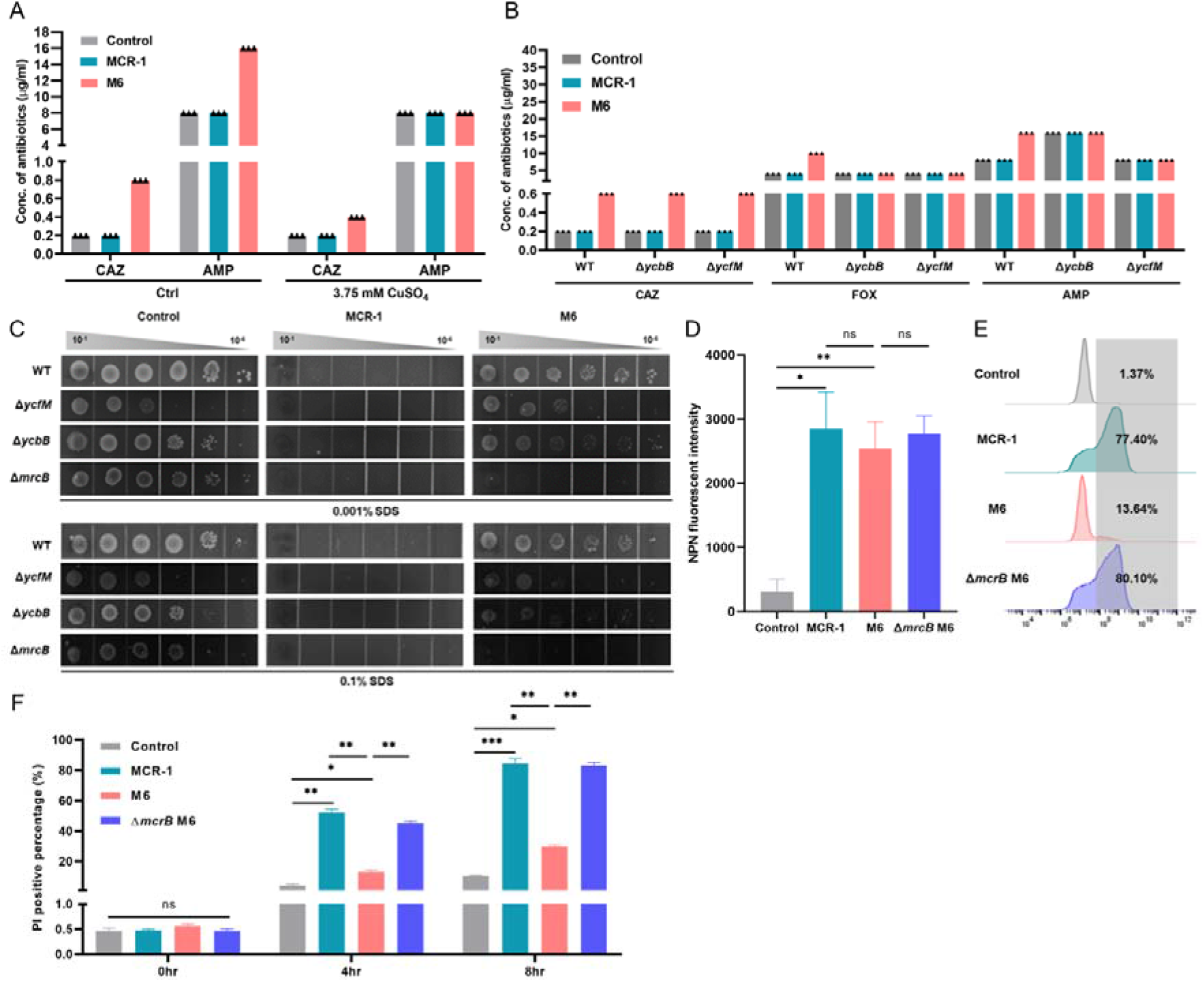
β-Lactams and SDS susceptibilities for LDT-defective strains. To test the role of LDTs in the M6-mediated phenotype, LdtD, PBP1B or LpoB mutant strains carrying empty vector (control), WT MCR-1 or M6 were generated. **(A)** Testing the antibiotic sensitivity of M6-expressing cells and MCR-1-expressing cells in the presence of copper. Overnight cultures of strains were sub-cultured into fresh LB broth with or without 3.75 mM CuSO_4_ at a ratio of 1:100 and induced with 0.2% arabinose for 2 hr. Next, the logarithmic-phase cultures were collected and adjusted to OD_600_=0.6, followed by spotting serial dilutions on LB agar plates containing CAZ or AMO, together with or without the addition of 3.75 mM CuSO_4_, and incubation at 37 °C for 16 hr to determine related MICs. Each triangle represents an independent experiment. **(B)** Testing the role of LDTs on β-lactam antibiotic susceptibility of M6. Overnight cultures of the indicated strains were sub-cultured into fresh LB broth at a ratio of 1:100 and induced with 0.2% arabinose for 2 hr. The logarithmic-phase cultures were collected and adjusted to OD_600_=0.6, followed by spotting serial dilutions on LB agar plates with target antibiotics and incubation at 37 °C for 16 hr. Each triangle represents an independent experiment. The experiments were performed three times with similar results. **(C)** Evaluation of the impact of LDTs on the sensitivity to SDS detergent. **(D)** *mrcB* deletion failed to reverse the OM permeability defect caused by M6. NPN uptake is represented by the background subtracted fluorescence at an excitation wavelength of 350 nm and emission wavelength of 420 nm. **(E-F)** *mrcB* deletion abolished M6-mediated inner membrane integrity. The inner membrane permeability was evaluated by PI staining assay. Overnight cultures were sub-cultured into fresh LB broth at a ratio of 1:100 and induced with 0.2% arabinose to express WT MCR-1 or M6. After induction for 4 hr and 8 hr, stationary and late-stationary phase cultures were collected, respectively, followed by staining with PI dye for 15 min. The PI-positive proportion was determined by flow cytometry and analysed by FlowJo version10 software. Representative results of three independent experiments are shown in **(E)**. All the above-described experiments were performed three times with similar results. Error bars indicate standard errors of the means (SEMs) for three biological replicates. A two-tailed unpaired *t* test was performed to determine the statistical significance of the data. ns, no significant difference; *, *P*< 0.1; **, *P*< 0.01; ***, *P*< 0.001.

**Figure S7.**
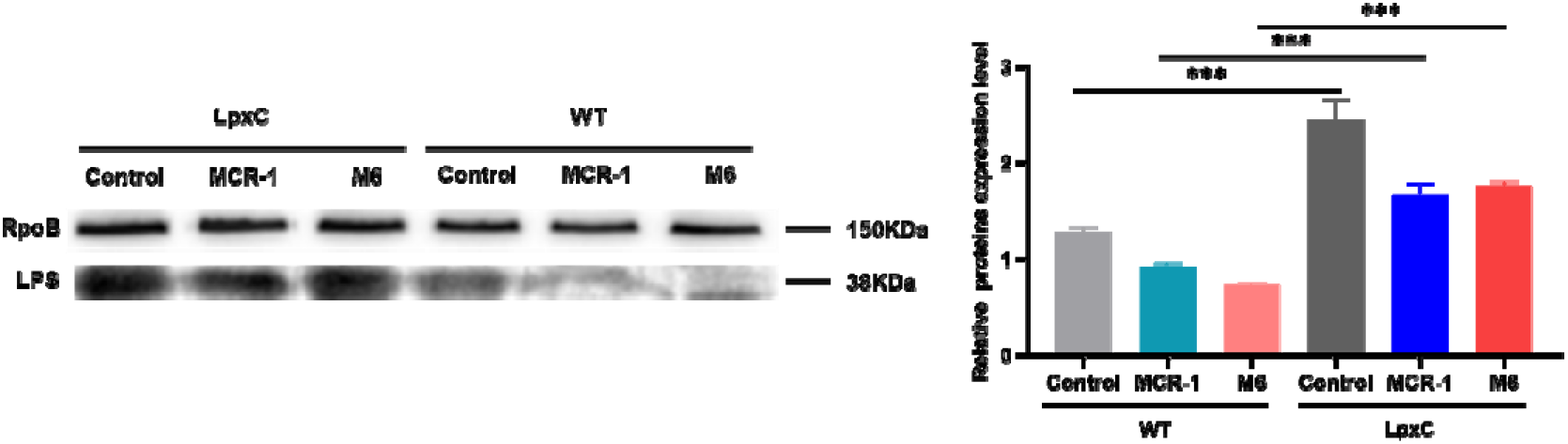
Construction of LpxC overexpression strains. The expression level of LPS was determined by western blot. pBAD24-LpxC was transformed into *E. coli* BW25113 carrying empty plasmid (control), MCR-1 or M6 separately. Error bars indicate standard errors of the means (SEM) for triple biological replicates. A two-tailed unpaired *t* test was performed to determine the statistical significance of the data. ***, *P*< 0.001. The bar graph was visualized with Prism 9 software.

**Figure S8.**
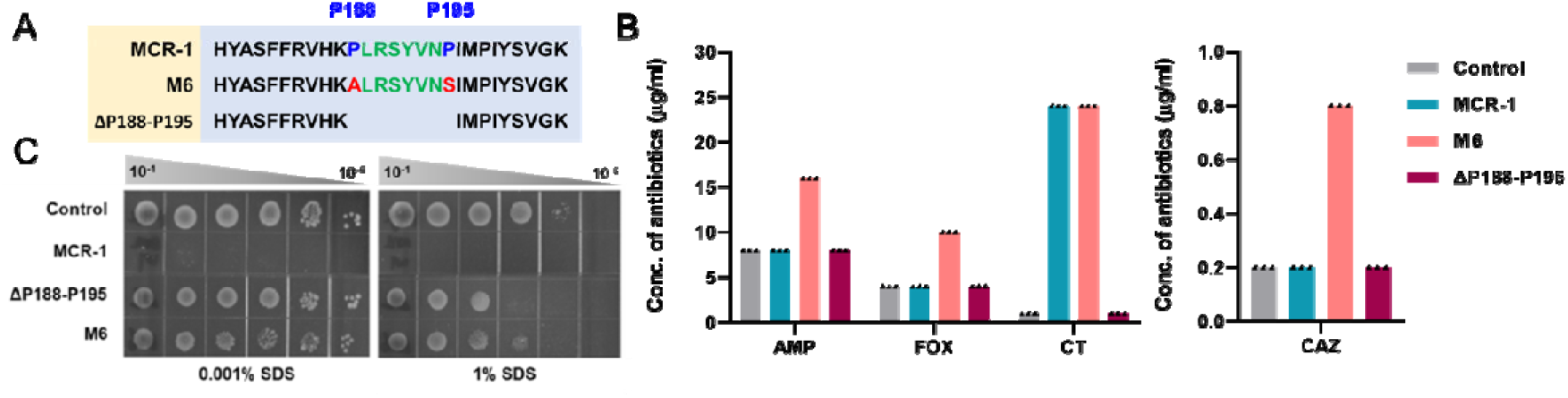
Necessity of P188-P195 for MCR-1 activity. **(A)** Deletion of the region that includes P188-P195. *E. coli* BW25113 carrying ΔP188-P195 was generated, and the susceptibilities towards colistin (CT) and β-lactam antibiotics (AMP, FOX and CAZ) were evaluated by agar dilution MIC tests **(B)**. **(C)** Deficiency of P188-P195 recovered cell wall permeability.

**Figure S9.**
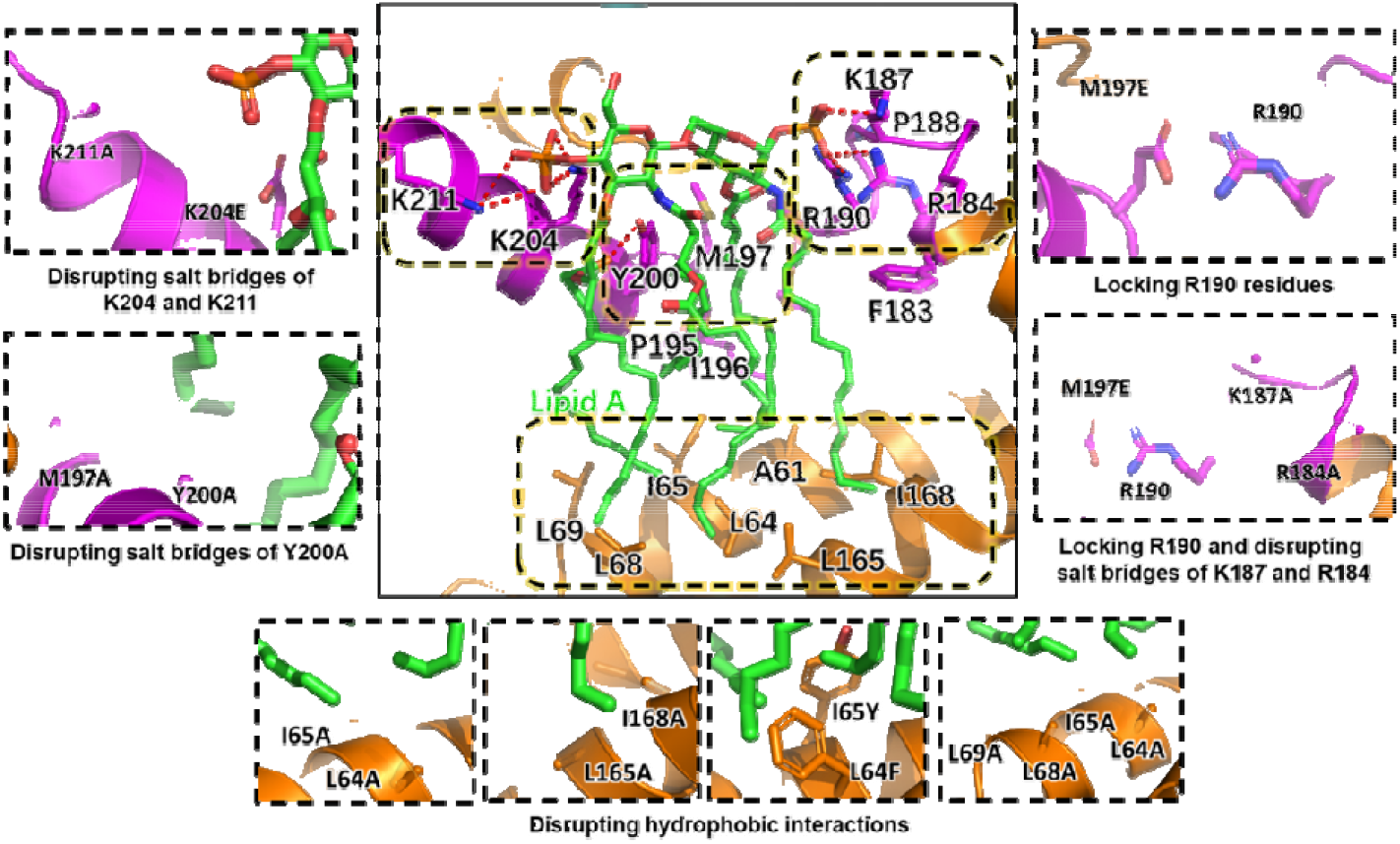
Mutations of essential regions for forming the lipid A binding cavity. The close-up view shows the MCR-1 lipid A binding cavity with LPS and the four regions responsible for anchoring LPS. The linker domain and transmembrane domain are in magenta and orange, respectively. LPS binding with MCR-1 is represented as a stick mode and coloured in green. The salt bridges for the interaction with LPS are shown in red. The change in structure with mutated residues in each region is shown in detail.

**Figure S10.**
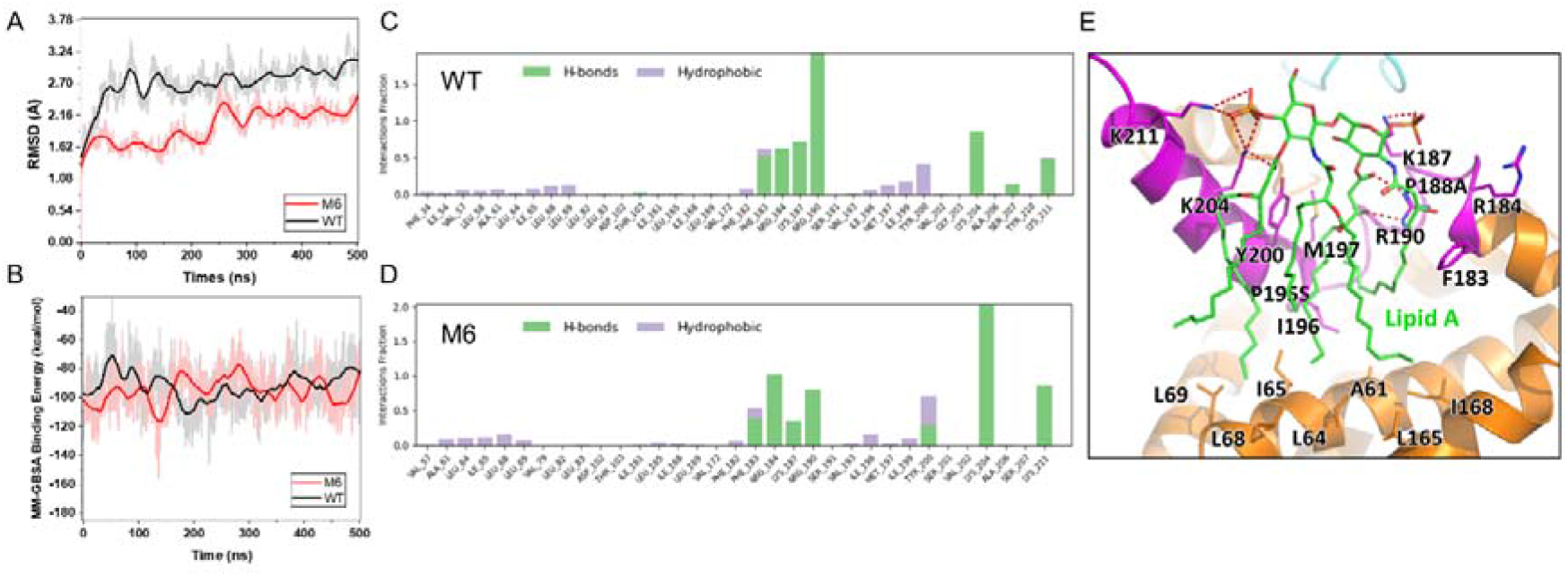
Detailed molecular interactions captured along the trajectories of MD simulations. **(A)** represents the RMSD fluctuations for WT and M6 of MCR1-lipid A. The variations in MCR1 MM-GBSA binding energy with WT and M6 during the MD trajectories are shown in **(B). (C)** and **(D)** show the statistics for the observed molecular interactions between lipid A and MCR-1 over the simulated trajectories of WT and the M6 mutant (the last 200-ns trajectories used for binding free energy calculations). **(E)** The predicted binding mode of lipid A to the M6 mutant.

**Figure S11.**
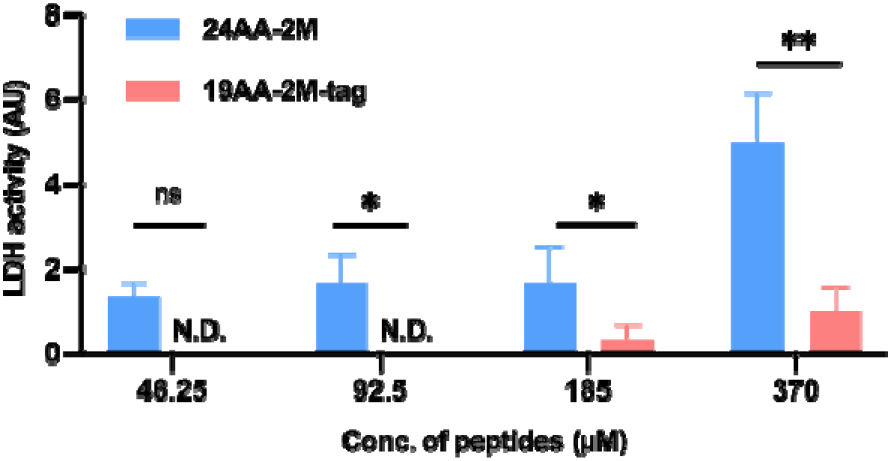
Permeabilization of synthetic peptides on mouse blood cells. To identify the cytotoxicity of 24AA-2M and 19AA-2M-tag, the cell permeabilizing effects of the indicated peptides on mouse blood cells were determined by an LDH-based TOX-7 kit (Sigma). Fresh healthy mouse blood was treated with the above peptides at concentrations of 46.25, 92.5, 185 or 370 µM. LDH activity was evaluated to determine LDH released from mousse cells. No LDH release was measured without peptide treatment. Error bars indicate standard errors of the means (SEMs) for three biological replicates. A two-tailed unpaired *t* test was performed to determine the statistical significance of the data. ns, no significant difference; *, *P*< 0.1; **, *P*< 0.01. The bar graph was visualized with Prism 9 software. N.D., not detected.

**Figure S12.**
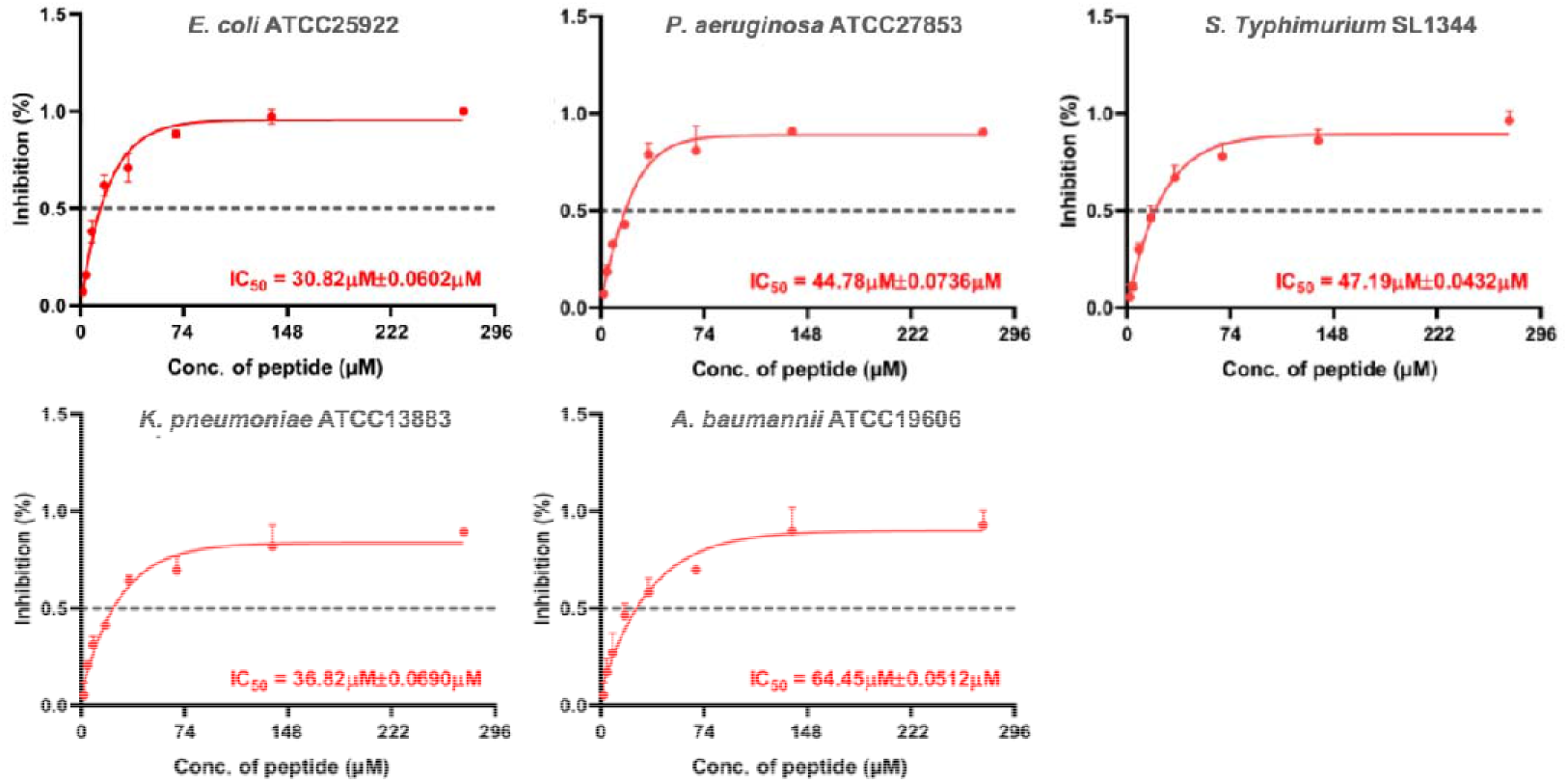
Inhibitory rate of 19AA-2M-tag among Gram-negative and Gram-positive strains. By treating *E. coli* ATCC25922, *P. aeruginosa* ATCC27853, *S. Typhimurium* SL1344, *K. pneumonic* ATCC13883, and *A. baumanii* ATCC19606 with 19AA-2M-tag in a series of concentrations, dose_Jresponse curves were generated. The *in vitro* inhibitory rate and IC_50_ value of various strains treated with 19AA-2M-tag are presented.

**Figure S13.**
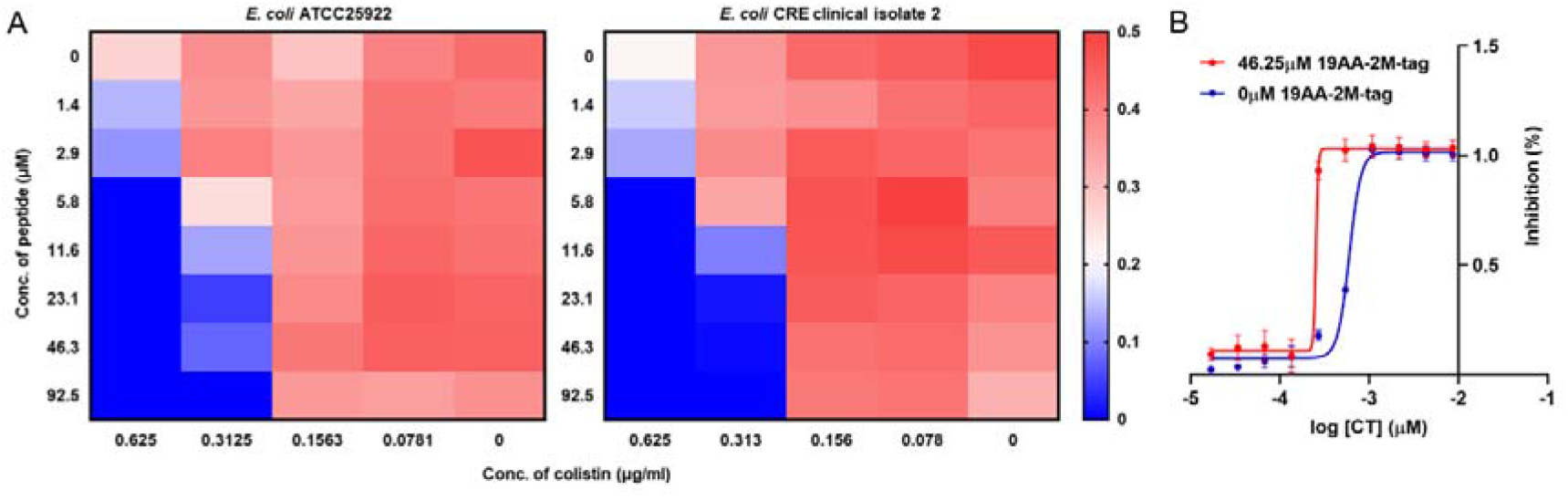
Synergistic inhibition of *E. coli* growth by 19AA-2M-tag and colistin. **(A)** *E. coli* ATCC25922 and a carbapenem-resistant *E. coli* isolated in the clinic were used to test the efficacy of a drug combination consisting of 19AA-2M-tag and colistin through a checkerboard assay. A total of 1× 10^3^ CFUs of the indicated strains were inoculated at initiation (T0), followed by treatment with drug combinations at various concentrations at 37 °C for 16 hr (Tn). The OD_600_ was measured before and after the treatment. The graphs show the value of OD_600 T0_ minus the OD_600 Tn_ of each well for both strains. A colour gradient heat map, with hot (red: OD_600_ = 0) to cool (blue: OD_600_ = 0.5) colours indicating low to high values The synergetic effect was further determined by calculating the FICI. **(B)** shows the inhibition curves of colistin with (red) or without (blue) the addition of 19AA-2M-tag. The dose_Jresponse curves of *E. coli* ATCC25922 were generated by measuring the inhibitory rate after treatment with a series of concentrations of colistin with or without co-treatment with 92.5 µM 19AA-2M-tag.

**Figure S14.**
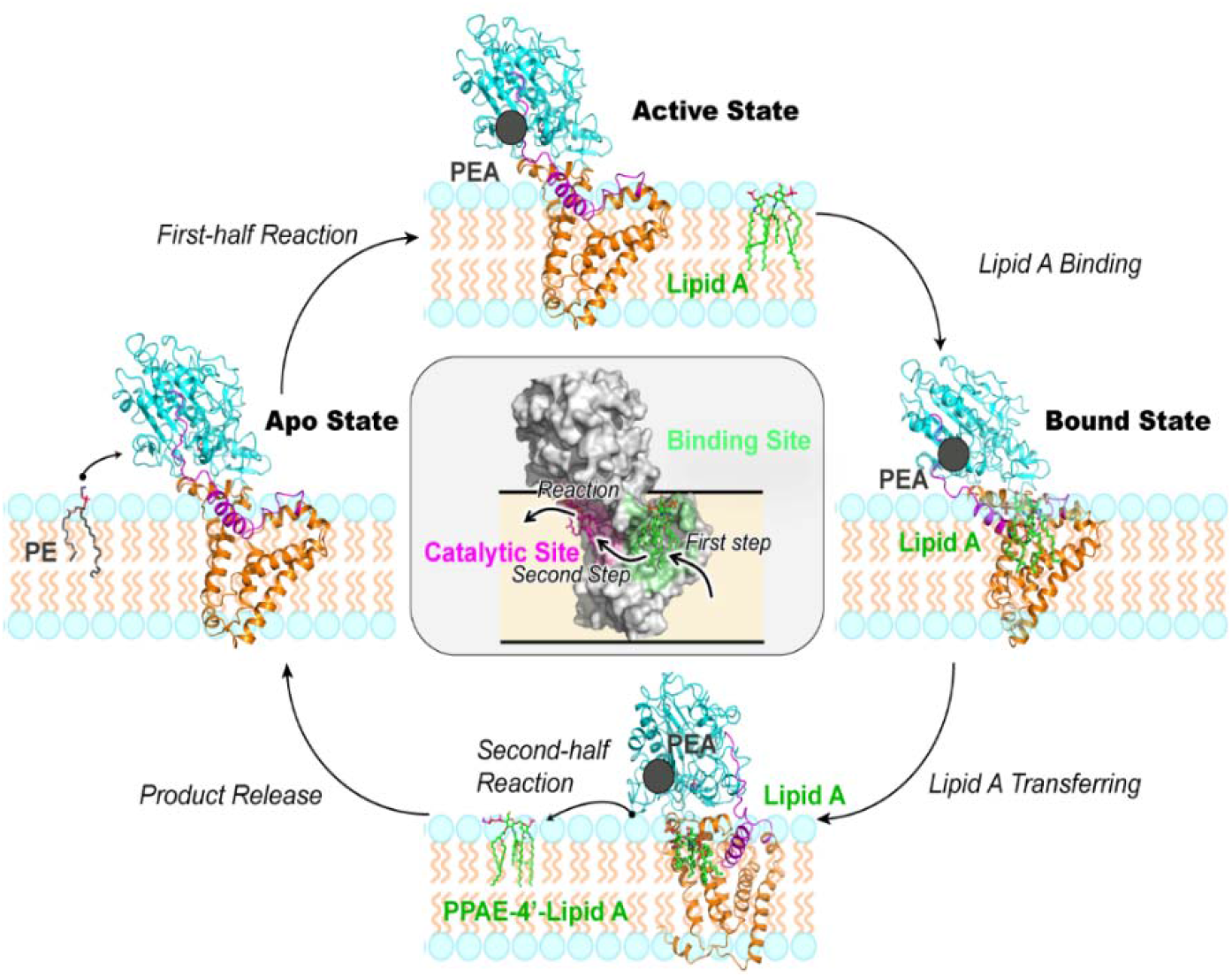
The “loading-transferring” mode of MCR-1. Modification of the LPS substrate of MCR-1 might be guaranteed by a two-step process, which requires the LPS substrate to be loaded and transferred to the catalytic domain to add the PEA group. During the first-half reaction, the Apo state MCR-1 interacts with the PE donor at the outer leaflet of the inner membrane and accepts a PEA group in the catalytic domain, transforming into the active state to interact with LPS. Next, LPS binds with the MCR-1 loading cavity at the linker domain, which is inserted into the lipid bilayer, followed by transfer to the catalytic domain exposed at the interface between the inner membrane and periplasm. During the second-half reaction, the PEA group at the catalytic domain added to the 1’- or 4’-phosphate group of LPS to form PEA-LPS, which was then released back into the inner membrane, and MCR-1 switched to the Apo state again.

**Figure S15.**
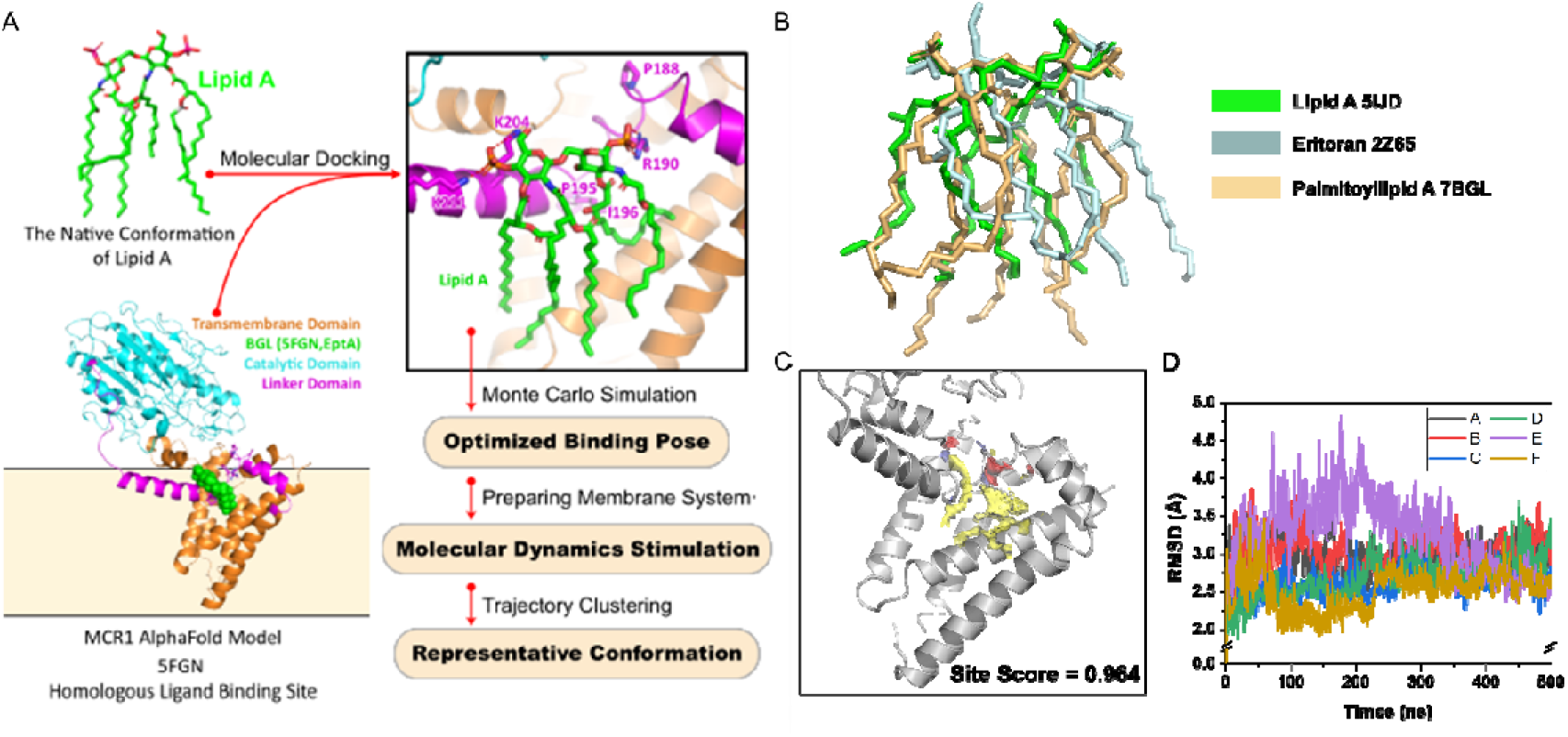
Analysis of the molecular dynamics simulation and binding energy. Molecular dynamic stimulation was utilized to discover the potential LPS binding cavity on the surface of MCR-1, and the workflow was designed as shown in **(A)**. Structural comparison within lipid A (PDB code: 5IJD) and its analogues: Eritoran (PDB code: 2Z65) and Palmitoyllpid A (PDB code: 7BGL) are shown in **(B)**. **(C)** The potential lipid A binding site on the surface of MCR-1 was discovered by SiteMap of Maestro. **(D)** shows the RMSD fluctuations for multiple trajectories of MCR1-lipid A.

**Supplementary Table 1.** Concentrations of antibiotics used for screening the MCR-1 library.

| Antibiotics | MIC |  |  |  |
| --- | --- | --- | --- | --- |
|  | 0.8x | 1x | 1.5x | 2x |
| AMP | 25.6 | 32 | 48 | 64 |
| CAZ | 0.4 | 0.5 | 0.75 | 1 |
| IMP | 0.2 | 0.25 | 0.375 | 0.5 |
| SM | 25.6 | 32 | 48 | 64 |
| TET | 6.4 | 8 | 12 | 16 |
| NAL | 6.4 | 8 | 12 | 16 |
| VAN | 204.8 | 256 | 384 | 512 |

Concentrations are in µg/ml.

**Supplementary Table 2.** Concentrations of β-lactams used for screening the MCR-1 library.

| Antibiotics | MIC |  |  |  |
| --- | --- | --- | --- | --- |
|  | 0.8x | 1x | 1.5x | 2x |
| CTX | 0.4 | 0.5 | 0.75 | 1 |
| FEP | 1.6 | 2 | 3 | 4 |
| CRO | 0.8 | 1 | 1.5 | 2 |
| ETP | 0.1 | 0.125 | 0.1875 | 0.25 |
| MEM | 0.025 | 0.031 | 0.047 | 0.063 |
| FOX | 3.2 | 4 | 6 | 8 |

Concentrations are in µg/ml.

**Supplementary Table 3.** Verification of MCR-1 isolates through Sanger sequencing.

| No. | Mutations | MICs (µg/ml) |  |  |
| --- | --- | --- | --- | --- |
|  |  | CAZ | AMP | FOX |
| 1 | V22D, A23D, A30T | 0.2 | 8 | 4 |
| 2 | S171A | 0.2 | 8 | 4 |
| 3 | P171L, A174V | 0.4 | 16 | 4 |
| 4 | L165P | 0.4 | 16 | 8 |
| 5 | WT | 0.2 | 8 | 4 |
| 6 | P188A, P195S | ≥0.6 | 16 | ≥10 |
| 7 | H186Q, P188T | 0.4 | 16 | 4 |
| 8 | P188R | 0.4 | 8 | 4 |
| 9 | A123E, R128L | 0.2 | 8 | 4 |
| 10 | WT | 0.2 | 8 | 4 |
| 11 | T53K | 0.2 | 8 | 4 |
| 12 | V512L, T514S, H515Y | 0.2 | 8 | 4 |
| 13 | P493L | 0.2 | 8 | 4 |
| 14 | D46V | 0.2 | 8 | 4 |
| 15 | WT | 0.2 | 8 | 4 |
| 16 | T558A, C562A | 0.2 | 8 | 4 |
| 17 | C563G | 0.2 | 8 | 4 |
| 18 | S177R | 0.4 | 8 | 4 |
| 19 | T29P | 0.2 | 8 | 4 |
| 20 | WT | 0.2 | 8 | 4 |
| 21 | A20S | 0.2 | 8 | 4 |
| 22 | V162G, V227E | 0.2 | 8 | 4 |
| 23 | V202G | 0.2 | 8 | 4 |
| 24 | F495C, F496F, D499G | 0.2 | 8 | 4 |
| 25 | WT | 0.2 | 8 | 4 |
| 26 | K212R, K500R, Q501R | 0.2 | 8 | 4 |
| 27 | Q265H | 0.2 | 8 | 4 |
| 28 | W8G | 0.2 | 8 | 4 |
| 29 | G87C, T96I, G98G | 0.2 | 8 | 4 |
| 30 | D366G, G368V | 0.2 | 8 | 4 |
| 31 | V19A, R157L, V533F | 0.2 | 8 | 4 |
| 32 | V19A, A342S, D346Y | 0.2 | 8 | 4 |
| 33 | WT | 0.2 | 8 | 4 |
| 34 | G368G (WT) | 0.2 | 8 | 4 |
| 35 | E418D, Q425R, A430D | 0.2 | 8 | 4 |
| 36 | WT | 0.2 | 8 | 4 |
| 37 | E418D, Q425R, A430D | 0.2 | 8 | 4 |

