## Supplementary material for "Identification of a novel MCR-1 variant displaying low level of co-resistance to β-lactam antibiotics that uncovers a potential novel antimicrobial peptide": Table S4-10: Table S4-6. The differentially expressed proteins .pdf

**Table S4. The differentially expressed proteins between *E. coli* BW25113 carrying WT MCR-1 and M6**

| Protein_ID | Fc.value | P.value | gene |
|---|---|---|---|
| WP_000830741.1 | 8.87473 | 1.2E-48 | *ampH* |
| WP_000471889.1 | 8.27719 | 2.2E-33 | *mgtA* |
| WP_000190655.1 | 6.69928 | 4.7E-16 | *yfhb* |
| WP_000732512.1 | 5.8143 | 5.8E-14 | *rstB* |
| WP_001295285.1 | 4.90911 | 6.5E-08 | *pmrD* |
| WP_000820410.1 | 4.89458 | 1.6E-19 | *agn43* |
| WP_001092508.1 | 4.6609 | 2E-18 | *rstA* |
| WP_001265471.1 | 4.043 | 9.1E-17 | *phoP* |
| WP_001380339.1 | 3.81865 | 1.4E-15 | *KXD89_14405* |
| WP_000705211.1 | 3.67125 | 1.4E-13 | *ynfB* |
| WP_001273658.1 | 3.54056 | 1.8E-05 | *ymdF* |
| WP_000433476.1 | 3.32089 | 7.1E-13 | *maeA* |
| WP_000983871.1 | 3.30813 | 4.9E-05 | *trpCF* |
| WP_000883034.1 | 3.22982 | 2.7E-11 | *glsA* |
| WP_000597196.1 | 3.20858 | 3E-12 | *slyB* |
| WP_000045315.1 | 3.19218 | 3.7E-12 | *hemL* |
| WP_000066707.1 | 2.82914 | 9.2E-06 | *fumB* |
| WP_000594599.1 | 2.80513 | 1.1E-05 | *lpxT* |
| WP_000689228.1 | 2.79104 | 0.0005 | *ytfK* |
| WP_000024560.1 | 2.76806 | 1.4E-05 | *cbpM* |
| WP_000753946.1 | 2.74409 | 1.3E-09 | *degP* |
| WP_001157540.1 | 2.62654 | 0.00107 | *fetA* |
| WP_001138581.1 | 2.55387 | 1E-07 | *speG* |
| WP_000953361.1 | 2.54374 | 0.00156 | *nikA* |
| WP_000828648.1 | 2.51459 | 0.00179 | *rmf* |
| WP_001300708.1 | 2.48639 | 3.8E-08 | *dacC* |
| WP_000491520.1 | 2.46353 | 0.00226 | *yedP* |
| WP_001292679.1 | 2.30525 | 2.2E-06 | *mdoB* |
| WP_000436922.1 | 2.27911 | 0.0053 | *ycbJ* |
| WP_001059922.1 | 2.2013 | 7.8E-06 | *hypE* |
| WP_000190854.1 | 2.1995 | 0.00078 | *ycgB* |
| WP_001253696.1 | 2.17048 | 0.00875 | *envZ* |
| WP_000394140.1 | 2.12624 | 1.9E-05 | *ansB* |
| WP_000624375.1 | 2.11382 | 2.3E-05 | *folA* |
| WP_000989419.1 | 2.09378 | 0.01246 | *astA* |
| WP_000925969.1 | 4.0901 | 1.7E-06 | *ldtD* |
| WP_000459630.1 | 2.08903 | 0.01273 | *tatD* |
| WP_001336277.1 | 2.05056 | 1.2E-05 | *bglA* |
| WP_001297409.1 | 2.03771 | 0.01611 | *tadA* |
| WP_000735412.1 | 2.01658 | 0.01775 | *phoQ* |
| WP_000081588.1 | 2.01513 | 1.8E-05 | *rpoS* |
| WP_001274963.1 | 2.00813 | 0.01845 | *pspB* |
| WP_000426427.1 | 0.48755 | 5.6E-07 | *dsdA* |
| WP_000845104.1 | 0.47602 | 2.3E-07 | *asnA* |

| | | | |
|---|---|---|---|
| WP_000037608.1 | 0.47527 | 2E-05 | *yhbO* |
| WP_001350521.1 | 0.46479 | 1.1E-05 | *yedQ* |
| WP_000758655.1 | 0.46019 | 8.4E-06 | *ygdR* |
| WP_000178044.1 | 0.4483 | 4.2E-06 | *mliC* |
| WP_001300387.1 | 0.44345 | 9.6E-05 | *garK* |
| WP_000890001.1 | 0.42062 | 6.7E-07 | *truC* |
| WP_001308392.1 | 0.41722 | 9.2E-13 | *ivy* |
| WP_001112301.1 | 0.41315 | 4E-07 | *galP* |
| WP_001045689.1 | 0.40299 | 1.3E-05 | *sbp* |
| WP_000646173.1 | 0.36176 | 1.1E-06 | *ybdR* |
| WP_000498253.1 | 0.34248 | 9.8E-14 | *osmB* |
| WP_000430057.1 | 0.34202 | 7.6E-10 | *fiu* |
| WP_000761225.1 | 0.32266 | 8.8E-11 | *malS* |
| WP_001309184.1 | 0.3172 | 1.6E-15 | *nanM* |
| AIN31628.1 | 0.22799 | 2.4E-17 | *BW25113_1171* |
| WP_000458809.1 | 0.2195 | 3.6E-18 | *hcp* |
| WP_000281320.1 | 0.21767 | 2.3E-18 | *ygaC* |
| WP_000868187.1 | 0.2058 | 9.9E-40 | *rpmJ* |
| WP_001295684.1 | 0.19484 | 8.7E-30 | *ydeN* |
| WP_000459379.1 | 0.14779 | 6.8E-20 | *malY* |
| WP_000726974.1 | 0.08873 | 8.6E-44 | *ymgG* |
| WP_000990333.1 | 0.0831 | 4.2E-46 | *amiB* |
| WP_001295734.1 | 0.07192 | 2.2E-51 | *nanC* |
| WP_000681354.1 | 0.57528 | 0.00779 | *nhaA* |
| WP_001166395.1 | 0.72617 | 0.02402 | *ispH* |
| WP_000597260.1 | 0.75275 | 0.03661 | *carA* |
| WP_000736007.1 | 0.61402 | 0.01873 | *mutT* |
| WP_000124438.1 | 0.69599 | 0.00576 | *fhuA* |
| WP_001158931.1 | 0.61302 | 0.01834 | *fhuC* |
| WP_000845394.1 | 1.8128 | 0.04469 | *clcA* |
| WP_001291990.1 | 0.73816 | 0.02397 | *gpt* |
| WP_001019920.1 | 0.61516 | 0.00524 | *DXE50_03530* |
| WP_001102108.1 | 0.65099 | 0.01359 | *KXD89_16980* |
| WP_000023927.1 | 0.63207 | 0.00837 | *ECMP0215528_0369* |
| WP_000052206.1 | 1.4751 | 0.02913 | *prpB* |
| WP_001275859.1 | 1.52746 | 0.00731 | *HMPREF1599_03159* |
| WP_000413677.1 | 0.7623 | 0.04746 | *ddl* |
| WP_001295332.1 | 1.64166 | 0.03482 | *psiF* |
| WP_001295328.1 | 0.64444 | 0.00063 | *DXE50_04210* |
| WP_001326929.1 | 0.6311 | 0.0264 | *KXD89_16745* |
| WP_000844874.1 | 0.64375 | 0.03352 | *priC* |
| WP_000738500.1 | 1.45019 | 0.01722 | *bor* |
| WP_001201845.1 | 1.42015 | 0.02377 | *ompT* |
| WP_001224604.1 | 1.62802 | 0.03801 | *KXD89_15970* |
| WP_000381303.1 | 0.58105 | 0.00896 | *A1UI_00656* |
| WP_000278509.1 | 1.75096 | 0.00052 | *A1UI_00672* |
| WP_000503931.1 | 1.94949 | 0.00445 | *tatE* |
| WP_000271153.1 | 1.86486 | 0.00043 | *nagA* |

| | | | |
|---|---|---|---|
| WP_001113989.1 | 0.6236 | 0.02278 | *nei* |
| WP_000254365.1 | 0.74511 | 0.03761 | *sdhD* |
| WP_000568275.1 | 0.66499 | 0.00418 | *DXE50_05885* |
| WP_000053415.1 | 0.65211 | 0.0009 | *galK* |
| WP_000767389.1 | 1.55538 | 0.01297 | *ECMP0215528_0917* |
| WP_000100800.1 | 1.93417 | 5.2E-05 | *dps* |
| WP_001295296.1 | 1.5718 | 0.0044 | *ompX* |
| WP_000815337.1 | 1.42197 | 0.04861 | *poxB* |
| WP_001295343.1 | 0.67762 | 0.00625 | *lolA* |
| WP_000850303.1 | 1.61106 | 0.04235 | *dmsA* |
| WP_000165879.1 | 1.44322 | 0.01857 | *A1UI_00969* |
| WP_000977920.1 | 0.73533 | 0.02199 | *ompF* |
| WP_000048252.1 | 1.60609 | 0.04371 | *yccX* |
| WP_001019197.1 | 0.60146 | 0.00349 | *DXE50_07850* |
| WP_000420621.1 | 1.42312 | 0.04808 | *cbpA* |
| WP_001151437.1 | 1.61626 | 0.00262 | *wrbA* |
| AIN31510.1 | 1.80798 | 0.04566 | - |
| WP_001144615.1 | 0.55664 | 0.00077 | *trhO* |
| pdb\|6H58\|00 | 0.73734 | 0.02338 | - |
| WP_000800153.1 | 0.55094 | 0.00415 | *bhsA* |
| WP_000284280.1 | 1.89919 | 0.00628 | *KXD89_12840* |
| WP_000702650.1 | 1.40496 | 0.02791 | *narH* |
| WP_000718995.1 | 0.69111 | 0.00478 | *galU* |
| WP_001360141.1 | 0.63673 | 0.0294 | *tonB* |
| WP_001056490.1 | 1.64087 | 0.035 | *KXD89_12400* |
| WP_001298828.1 | 0.50575 | 9.1E-05 | *hslJ* |
| WP_001163872.1 | 1.35219 | 0.04805 | *patD* |
| WP_000841554.1 | 1.5223 | 0.01819 | *DXE50_16450* |
| WP_000246019.1 | 1.70277 | 0.02342 | *gadC* |
| WP_000358930.1 | 1.77542 | 0.01448 | *O3K_13030* |
| WP_000257409.1 | 0.64259 | 0.03281 | *glsB* |
| WP_000214712.1 | 1.386 | 0.03402 | *ydfZ* |
| WP_001300836.1 | 1.85322 | 0.03727 | *A1UI_01776* |
| WP_001043339.1 | 1.75819 | 0.00047 | *A1UI_01805* |
| WP_000483353.1 | 1.47956 | 0.01249 | *hdhA* |
| WP_001296943.1 | 0.55992 | 5.1E-05 | *slyA* |
| WP_000069375.1 | 0.69236 | 0.00502 | *ppsA* |
| WP_001154168.1 | 1.47744 | 0.02847 | *btuE* |
| WP_000267650.1 | 1.60056 | 0.00809 | *DXE50_14915* |
| WP_000077872.1 | 1.40442 | 0.02807 | *katE* |
| WP_000994973.1 | 1.86296 | 0.03567 | *astB* |
| WP_000081983.1 | 1.56581 | 0.00471 | *astC* |
| WP_001019882.1 | 1.48369 | 0.01193 | *DXE50_14615* |
| WP_000394983.1 | 0.56835 | 0.00655 | *DXE50_14440* |
| WP_000010115.1 | 0.50815 | 0.00115 | *rlmA* |
| WP_000916763.1 | 0.70901 | 0.0478 | *DXE50_14305* |
| WP_000936927.1 | 0.61509 | 0.01915 | *ptrB* |
| WP_001350516.1 | 1.512 | 0.02018 | *yebF* |

| | | | |
|---|---|---|---|
| WP_000891621.1 | 1.70638 | 0.00258 | *A1UI_02070* |
| WP_001245695.1 | 1.59433 | 0.00864 | *amyA* |
| WP_000740106.1 | 1.38884 | 0.03303 | *msrP* |
| WP_001350529.1 | 1.60797 | 0.0432 | *KXD89_08150* |
| WP_000490679.1 | 0.75665 | 0.04077 | *DXE50_12405* |
| WP_000182899.1 | 0.76364 | 0.04917 | *gatA* |
| WP_000153067.1 | 1.43572 | 0.04266 | *O3K_08910* |
| WP_000422182.1 | 0.67197 | 0.02223 | *yojI* |
| WP_000566471.1 | 1.78703 | 0.0134 | *ECMP0215528_2665* |
| WP_000748271.1 | 1.40828 | 0.02695 | *A1UI_02525* |
| WP_000991370.1 | 0.50596 | 9.2E-05 | *evgA* |
| WP_000497400.1 | 0.6686 | 0.0019 | *A1UI_02594* |
| WP_000021036.1 | 0.50089 | 0.0009 | *cysA* |
| WP_001003709.1 | 1.65615 | 0.00163 | *tal* |
| WP_000087280.1 | 1.49552 | 0.0238 | *WG3_03045* |
| WP_001300814.1 | 0.71333 | 0.01732 | *A1UI_02650* |
| WP_000133592.1 | 1.66697 | 0.00143 | *pepB* |
| WP_000883122.1 | 0.71352 | 0.0174 | *hmp* |
| WP_000986023.1 | 0.60834 | 0.01661 | *acpS* |
| WP_000178456.1 | 1.47859 | 0.01263 | *raiA* |
| WP_001203437.1 | 1.61073 | 0.04244 | *bamE* |
| WP_000271909.1 | 1.38588 | 0.03406 | *lhgO* |
| WP_000522415.1 | 1.59041 | 0.00354 | *kbp* |
| WP_000777969.1 | 0.5006 | 0.00089 | *nrdF* |
| WP_001199973.1 | 1.87674 | 0.00037 | *queE* |
| WP_000626422.1 | 0.68335 | 0.00352 | *sdaB* |
| WP_000016907.1 | 1.98765 | 0.00342 | *DXE50_18860* |
| WP_001326497.1 | 0.69386 | 0.01014 | *loiP* |
| WP_000994920.1 | 0.64399 | 0.03367 | *yggU* |
| WP_000013149.1 | 1.40691 | 0.02734 | *dkgA* |
| WP_000712658.1 | 1.74598 | 0.00055 | *ygiW* |
| WP_000735278.1 | 1.70086 | 0.00095 | *tolC* |
| WP_000046281.1 | 0.68739 | 0.0084 | *A1UI_03332* |
| WP_001125331.1 | 0.72625 | 0.02406 | *DXE50_20220* |
| WP_001066494.1 | 0.69351 | 0.03524 | *A1UI_03348* |
| WP_000121433.1 | 1.42144 | 0.04885 | *A1UI_03359* |
| WP_000031415.1 | 1.69309 | 0.00104 | *ECMP0215528_3607* |
| WP_000785722.1 | 0.52762 | 8.2E-06 | *DXE50_20455* |
| WP_000548347.1 | 0.56752 | 0.00642 | *tdcB* |
| AIN33471.1 | 0.59149 | 0.00024 | - |
| WP_001058209.1 | 0.58686 | 0.0022 | *garL* |
| WP_001273753.1 | 0.64948 | 0.00247 | *garD* |
| pdb\|6H58\|oo | 0.74589 | 0.03012 | - |
| WP_001300352.1 | 0.6827 | 0.00343 | *gltB* |
| WP_000081674.1 | 0.67112 | 0.0051 | *gltD* |
| WP_000979882.1 | 0.61511 | 0.00523 | *ECMP0215528_3729* |
| WP_000209011.1 | 0.72079 | 0.02099 | *nanK* |
| WP_000224714.1 | 0.6379 | 0.00045 | *nanA* |

| | | | |
|---|---|---|---|
| WP_000695690.1 | 1.98787 | 0.02023 | *O3K_02775* |
| WP_000354622.1 | 1.38054 | 0.03599 | *accB* |
| WP_001219652.1 | 1.90568 | 0.02941 | *dusB* |
| WP_001157751.1 | 1.44043 | 0.01914 | *ompR* |
| WP_000856737.1 | 1.65146 | 0.03269 | *greB* |
| WP_000444342.1 | 0.69685 | 0.01104 | *malQ* |
| WP_000081909.1 | 0.56447 | 5.3E-06 | *malP* |
| WP_000906961.1 | 0.64756 | 0.00231 | *malT* |
| WP_001283723.1 | 1.49657 | 0.02356 | *glgB* |
| WP_000108330.1 | 0.73424 | 0.02921 | *gntK* |
| WP_001350553.1 | 1.88711 | 0.00033 | *slp* |
| WP_000372240.1 | 1.60953 | 0.00283 | *ECMP0215528_4059* |
| WP_001296794.1 | 1.9492 | 0.02414 | *ECMP0215528_4067* |
| WP_001295233.1 | 0.66721 | 0.01996 | *HMPREF1589_01249* |
| pdb\|6H58\|11 | 0.62209 | 0.0002 | - |
| WP_001243431.1 | 0.65765 | 0.00327 | *ibpB* |
| WP_001243437.1 | 0.56797 | 6.8E-06 | *ibpA* |
| WP_001291268.1 | 1.6522 | 0.00466 | *yieF* |
| WP_000763724.1 | 0.72424 | 0.02289 | *EL76_4667* |
| WP_000281668.1 | 0.66034 | 0.04499 | *A1UI_04118* |
| WP_000591375.1 | 0.7014 | 0.00704 | - |
| WP_000940106.1 | 0.71048 | 0.04915 | *DXE50_25690* |
| WP_001252058.1 | 0.60827 | 0.01659 | *malG* |
| WP_001297290.1 | 0.55203 | 0.00064 | *malF* |
| WP_000695387.1 | 0.55722 | 3.1E-06 | *malE* |
| WP_000179165.1 | 0.6329 | 0.00035 | *malK* |
| WP_000973663.1 | 0.58528 | 2.2E-05 | *lamB* |
| WP_001326641.1 | 0.67306 | 0.0023 | *malM* |
| WP_000002907.1 | 0.70491 | 0.04419 | *DXE50_25955* |
| WP_001014565.1 | 1.89183 | 0.0066 | *EKN05_018200* |
| WP_001131339.1 | 1.81878 | 0.04351 | *HMPREF1602_03064* |
| WP_001298520.1 | 1.66727 | 0.00396 | *HMPREF1599_00791* |
| WP_001295074.1 | 1.62661 | 0.00232 | *lysS* |
| WP_001350567.1 | 1.95734 | 0.02326 | *KXD89_19795* |
| WP_000853753.1 | 0.72311 | 0.01493 | *fbp* |
| WP_001336303.1 | 0.67935 | 0.00299 | *G711_01352* |
| WP_001407733.1 | 0.75656 | 0.04067 | *treB* |
| WP_001181307.1 | 0.59136 | 0.01141 | *G711_01354* |
| WP_000148581.1 | 0.71385 | 0.01095 | *pyrI* |
| WP_000013046.1 | 0.65255 | 0.00092 | *pyrB* |
| WP_000583469.1 | 0.75447 | 0.03841 | *EKN05_017700* |
| WP_001309160.1 | 1.69988 | 0.02386 | *G894_04371* |
| WP_000611568.1 | 0.68585 | 0.00802 | *A13A_04926* |
| WP_000879155.1 | 0.56818 | 0.00117 | *HMPREF1599_03219* |
| WP_000188283.1 | 0.51124 | 2.9E-06 | *HMPREF1599_03218* |
| WP_000563040.1 | 0.60878 | 0.01677 | *G925_04733* |
| WP_001338221.1 | 0.66377 | 0.0477 | *O3K_21575* |
| WP_001264707.1 | 0.81635 | 0.1628 | *A13A_05080* |

| | | | |
|---|---|---|---|
| WP_000241662.1 | 0.74519 | 0.09024 | *thrB* |
| WP_000781074.1 | 0.95181 | 0.93496 | *thrC* |
| WP_000906197.1 | 1.03389 | 0.88064 | *yaaA* |
| WP_000130185.1 | 0.92334 | 0.73266 | *tal* |
| WP_001295414.1 | 0.97072 | 0.79965 | *EKN05_016655* |
| WP_000516135.1 | 0.84478 | 0.27015 | *dnaK* |
| WP_001118476.1 | 0.95135 | 0.93171 | *dnaJ* |
| pdb\|6H58\|tt | 0.88207 | 0.46355 | - |
| WP_000767329.1 | 0.91887 | 0.62089 | *ribF* |
| WP_001286857.1 | 0.90312 | 0.59533 | *ileS* |
| WP_000004655.1 | 0.84705 | 0.23238 | *fkpB* |
| WP_001239142.1 | 1.08333 | 0.67653 | *rihC* |
| WP_000543604.1 | 1.21412 | 0.28519 | *dapB* |
| WP_001126348.1 | 0.77589 | 0.06712 | *carB* |
| WP_000377098.1 | 1.17225 | 0.60238 | *kefC* |
| WP_000257192.1 | 1.05322 | 0.79791 | *apaH* |
| WP_000610901.1 | 1.31047 | 0.36755 | *apaG* |
| WP_001065381.1 | 0.88888 | 0.38893 | *rsmA* |
| WP_000241277.1 | 0.88659 | 0.48433 | *pdxA* |
| WP_000800457.1 | 0.98514 | 0.88549 | *surA* |
| WP_000746151.1 | 1.04462 | 0.62887 | *lptD* |
| WP_001200579.1 | 1.13847 | 0.58471 | *djlA* |
| WP_000525176.1 | 0.85565 | 0.44053 | *rluA* |
| WP_001117011.1 | 0.92429 | 0.7393 | *rapA* |
| WP_001300811.1 | 0.9002 | 0.57628 | *araC* |
| WP_001301364.1 | 1.20395 | 0.4322 | *HMPREF1589_03362* |
| WP_000818228.1 | 0.85399 | 0.25514 | *leuD* |
| WP_001140652.1 | 0.90179 | 0.44603 | *leuC* |
| WP_000042353.1 | 1.04881 | 0.81646 | *leuB* |
| WP_000082850.1 | 0.93193 | 0.5925 | *leuA* |
| WP_001115472.1 | 0.68064 | 0.0628 | *A1UI_04836* |
| WP_001295534.1 | 0.8865 | 0.48395 | *A1UI_04837* |
| WP_001300383.1 | 0.94017 | 0.74674 | *A1UI_04838* |
| WP_000762401.1 | 0.92887 | 0.77145 | *cra* |
| WP_001295533.1 | 1.13733 | 0.67605 | *mraZ* |
| WP_000970479.1 | 0.96734 | 0.78113 | *rsmH* |
| WP_000625658.1 | 0.87573 | 0.50867 | *ftsL* |
| WP_000642196.1 | 1.09243 | 0.71129 | *ftsI* |
| WP_000775093.1 | 0.95324 | 0.94522 | *murE* |
| WP_000626685.1 | 0.98404 | 0.87311 | *murF* |
| WP_000964131.1 | 1.12282 | 0.70836 | *mraY* |
| WP_000796481.1 | 1.01371 | 0.97007 | *murD* |
| WP_001295532.1 | 0.83355 | 0.37032 | *ftsW* |
| WP_000016560.1 | 1.20449 | 0.43107 | *murG* |
| WP_001096049.1 | 1.01332 | 0.97184 | *murC* |
| WP_000130056.1 | 1.01358 | 0.97067 | *ddl* |
| WP_000075748.1 | 0.75121 | 0.16423 | *ftsQ* |
| WP_000588474.1 | 0.94555 | 0.66349 | *ftsA* |

| | | | |
|---|---|---|---|
| WP_000462776.1 | 1.0943 | 0.45192 | *ftsZ* |
| WP_000595482.1 | 1.48293 | 0.18594 | *lpxC* |
| WP_000905789.1 | 0.8173 | 0.1658 | *secA* |
| WP_000005042.1 | 1.12315 | 0.70761 | *yacG* |
| WP_001194734.1 | 0.89325 | 0.51151 | *zapD* |
| WP_001269520.1 | 0.85384 | 0.43456 | *coaE* |
| WP_001217338.1 | 0.95184 | 0.93517 | *guaC* |
| WP_001135174.1 | 0.93232 | 0.59449 | *nadC* |
| WP_000923721.1 | 1.01528 | 0.97549 | *ampD* |
| WP_000172005.1 | 0.99652 | 0.96341 | *ampE* |
| WP_000331776.1 | 0.99487 | 0.9684 | *pdhR* |
| WP_000003820.1 | 0.91737 | 0.69133 | *aceE* |
| WP_000963518.1 | 0.97162 | 0.94895 | *aceF* |
| WP_000102485.1 | 0.90815 | 0.62873 | *lpdA* |
| WP_001307570.1 | 1.01333 | 0.75858 | *acnB* |
| WP_000384306.1 | 1.26451 | 0.4358 | *yacL* |
| WP_000734287.1 | 0.9611 | 0.74704 | *speD* |
| WP_000818411.1 | 1.00619 | 0.99511 | *speE* |
| WP_001295568.1 | 0.80198 | 0.28028 | *DXE50_02840* |
| WP_001189647.1 | 1.33372 | 0.2214 | *A1UI_04880* |
| WP_001306211.1 | 0.9638 | 0.82485 | *HMPREF1595_04252* |
| WP_000683335.1 | 0.76557 | 0.05846 | *DXE50_02855* |
| WP_000651599.1 | 0.94031 | 0.85274 | *DXE50_02860* |
| WP_000150637.1 | 0.98231 | 0.86352 | *DXE50_02865* |
| WP_000621515.1 | 1.00898 | 0.97557 | *panD* |
| WP_000905383.1 | 1.03044 | 0.89572 | *panC* |
| WP_000805497.1 | 1.10127 | 0.43012 | *panB* |
| WP_000215139.1 | 0.84768 | 0.41458 | *folK* |
| WP_010723084.1 | 0.76814 | 0.12839 | *pcnB* |
| WP_001155227.1 | 1.14991 | 0.29874 | *dksA* |
| WP_000396036.1 | 0.90082 | 0.59857 | *sfsA* |
| WP_001350480.1 | 0.9749 | 0.87627 | *hrpB* |
| WP_000918162.1 | 0.96317 | 0.98922 | *mrcB* |
| WP_001310529.1 | 0.86023 | 0.45571 | *HMPREF1602_04663* |
| WP_001295564.1 | 1.38027 | 0.07156 | *erpA* |
| WP_000689844.1 | 0.93655 | 0.8259 | *mtnN* |
| WP_000057073.1 | 1.27107 | 0.42547 | *dgt* |
| WP_000272188.1 | 1.12433 | 0.52901 | *yaeH* |
| WP_001186650.1 | 0.92808 | 0.76589 | *dapD* |
| WP_001094586.1 | 0.88223 | 0.36101 | *glnD* |
| WP_001018194.1 | 0.85153 | 0.30086 | *map* |
| pdb\|7OE0\|B | 0.94253 | 0.86861 | - |
| WP_000818114.1 | 0.85373 | 0.31128 | *tsf* |
| WP_000224573.1 | 0.87271 | 0.4098 | *pyrH* |
| WP_000622418.1 | 0.98028 | 0.90812 | *frr* |
| WP_000811923.1 | 0.78308 | 0.2328 | *dxr* |
| WP_001295562.1 | 1.05958 | 0.85999 | *ispU* |
| WP_000922446.1 | 1.46333 | 0.2015 | *cdsA* |

| | | | |
|---|---|---|---|
| WP_001295561.1 | 1.07821 | 0.7534 | *rseP* |
| WP_001240896.1 | 0.96483 | 0.9813 | *bamA* |
| WP_000758956.1 | 1.17448 | 0.24545 | *skp* |
| WP_001139279.1 | 0.91187 | 0.49311 | *lpxD* |
| WP_000210739.1 | 0.9834 | 0.89354 | *fabZ* |
| WP_000565966.1 | 1.09192 | 0.64385 | *lpxA* |
| WP_000139654.1 | 1.46351 | 0.20135 | *lpxB* |
| WP_000569430.1 | 0.94992 | 0.78421 | *rnhB* |
| WP_001294757.1 | 1.13581 | 0.6794 | *dnaE* |
| WP_000055741.1 | 0.96903 | 0.96124 | *accA* |
| WP_001020973.1 | 1.50739 | 0.16806 | *ldcC* |
| WP_000901099.1 | 1.01331 | 0.96099 | *KXD89_17755* |
| WP_000176549.1 | 1.14245 | 0.6649 | *tilS* |
| AIN30718.1 | 1.3214 | 0.23674 | - |
| WP_000417058.1 | 1.33643 | 0.21813 | *yaeP* |
| WP_001185290.1 | 0.95263 | 0.79466 | *DXE50_03195* |
| WP_000239163.1 | 1.49961 | 0.08476 | *nlpE* |
| WP_001260717.1 | 0.90438 | 0.60362 | *proS* |
| WP_000094011.1 | 1.01606 | 0.97338 | *tsaA* |
| WP_001202329.1 | 0.86006 | 0.27613 | *rcsF* |
| WP_000874226.1 | 1.01418 | 0.75491 | *WG3_00475* |
| WP_000594006.1 | 1.05222 | 0.83375 | *metN* |
| WP_001140187.1 | 0.75849 | 0.11119 | *gmhB* |
| WP_000997010.1 | 0.74602 | 0.09145 | *dkgB* |
| WP_000648572.1 | 1.52717 | 0.15476 | *KXD89_17630* |
| WP_001230983.1 | 1.47639 | 0.19101 | *yafD* |
| WP_001052715.1 | 1.0794 | 0.74984 | *gloB* |
| WP_000917883.1 | 0.85327 | 0.43271 | *rnhA* |
| WP_001340895.1 | 0.92192 | 0.67724 | *dnaQ* |
| WP_001118036.1 | 1.31126 | 0.25003 | *yafV* |
| WP_000973083.1 | 1.04626 | 0.82732 | *A1UI_00324* |
| WP_000284050.1 | 1.07233 | 0.52551 | *lpcA* |
| WP_000333380.1 | 0.92693 | 0.69622 | *ECMP0215528_0302* |
| WP_000729703.1 | 1.27459 | 0.30345 | *dinJ* |
| WP_001292994.1 | 0.95562 | 0.96221 | *pepD* |
| WP_000189532.1 | 0.97586 | 0.82797 | *frsA* |
| WP_000174689.1 | 1.13901 | 0.48164 | *crl* |
| WP_001285288.1 | 0.96482 | 0.76734 | *proB* |
| WP_000893278.1 | 1.08697 | 0.47563 | *proA* |
| WP_000081352.1 | 1.38877 | 0.27179 | *A1UI_00392* |
| WP_001136613.1 | 1.0884 | 0.65713 | *yagE* |
| WP_000151261.1 | 0.99478 | 0.95679 | *yagF* |
| WP_000406871.1 | 0.97042 | 0.86329 | *yagH* |
| WP_001121657.1 | 1.0879 | 0.78998 | *kdgR_2* |
| WP_000803998.1 | 0.66951 | 0.0525 | *rpmE2* |
| AIN30811.1 | 1.17912 | 0.58857 | - |
| WP_001046307.1 | 1.27557 | 0.30191 | *KXD89_16970* |
| WP_000370307.1 | 0.77926 | 0.07675 | *KXD89_16990* |

| | | | |
|---|---|---|---|
| WP_000089110.1 | 1.0248 | 0.92265 | *betB* |
| WP_000665120.1 | 0.83231 | 0.36654 | *KZW89_19055* |
| WP_000692754.1 | 1.30743 | 0.07473 | *A1UI_00416* |
| WP_000691956.1 | 1.34921 | 0.09457 | *ECMP0215528_0390* |
| WP_001285927.1 | 1.4093 | 0.05475 | *prpC* |
| WP_000076233.1 | 0.80794 | 0.29627 | *codB* |
| WP_001301240.1 | 0.81402 | 0.1419 | *A1UI_00427* |
| WP_000952503.1 | 0.83302 | 0.36868 | *A1UI_00428* |
| WP_071843335.1 | 0.86787 | 0.41133 | - |
| WP_001301325.1 | 0.91205 | 0.64016 | *KXD89_17195* |
| WP_001013499.1 | 0.84016 | 0.39072 | *mhpE* |
| WP_001096705.1 | 0.99066 | 0.94907 | *DXE50_03945* |
| WP_000419081.1 | 0.8602 | 0.38301 | *fghA* |
| WP_000842106.1 | 0.9761 | 0.82925 | *frmA* |
| WP_001141271.1 | 0.94179 | 0.72368 | *frmR* |
| WP_001295337.1 | 1.15036 | 0.29769 | *hemB* |
| WP_000092067.1 | 0.86177 | 0.46089 | *yaiW* |
| WP_000792970.1 | 1.19231 | 0.33449 | *iraP* |
| WP_001295331.1 | 1.20797 | 0.29849 | *proC* |
| WP_000158159.1 | 1.19008 | 0.46177 | *yaiI* |
| WP_001276420.1 | 0.75786 | 0.17732 | *DXE50_04100* |
| WP_000941942.1 | 1.12122 | 0.63031 | *ppnP* |
| WP_001298537.1 | 0.95101 | 0.69261 | *rdgC* |
| WP_001219309.1 | 1.17446 | 0.49682 | *mak* |
| WP_000698951.1 | 1.15614 | 0.63566 | *sbcC* |
| WP_000113933.1 | 0.88531 | 0.47918 | *phoB* |
| WP_000893623.1 | 1.03253 | 0.92972 | *phoR* |
| WP_000149639.1 | 0.97558 | 0.88318 | *brnQ* |
| WP_001295329.1 | 1.26436 | 0.43603 | *proY* |
| AIN30902.1 | 1.22995 | 0.49346 | - |
| WP_001266503.1 | 1.11651 | 0.55543 | *queA* |
| WP_000667319.1 | 0.81228 | 0.13794 | *tgt* |
| WP_000007629.1 | 1.03277 | 0.67646 | *yajC* |
| WP_000934822.1 | 0.90534 | 0.61001 | *secD* |
| WP_000046637.1 | 0.93235 | 0.59466 | *secF* |
| WP_000974813.1 | 0.91956 | 0.62391 | *DXE50_04205* |
| WP_000543535.1 | 1.02972 | 0.90644 | *nrdR* |
| WP_001150457.1 | 0.97823 | 0.89166 | *ribD* |
| WP_001021161.1 | 0.77525 | 0.06607 | *ribH* |
| WP_000801125.1 | 1.02127 | 0.93626 | *nusB* |
| WP_000742109.1 | 0.91037 | 0.63389 | *thiL* |
| WP_001199800.1 | 1.25965 | 0.20084 | *A1UI_00501* |
| WP_000006797.1 | 0.91719 | 0.51878 | *dxs* |
| WP_000347217.1 | 1.07218 | 0.77167 | *A1UI_00503* |
| WP_001124935.1 | 0.8572 | 0.44564 | *xseB* |
| WP_000668662.1 | 0.91794 | 0.52245 | *thiI* |
| WP_001276305.1 | 0.90732 | 0.57071 | *A1UI_00506* |
| WP_000705865.1 | 0.99262 | 0.95808 | *panE* |

| | | | |
|---|---|---|---|
| WP_001138904.1 | 0.91527 | 0.67691 | *yajQ* |
| WP_000467180.1 | 0.8902 | 0.51282 | *DXE50_04315* |
| WP_001239436.1 | 0.80551 | 0.13099 | *WG3_00715* |
| WP_001295326.1 | 0.82911 | 0.20657 | *DXE50_04330* |
| WP_000973448.1 | 1.04105 | 0.84959 | *bolA* |
| WP_001198386.1 | 0.88509 | 0.48161 | *tig* |
| WP_000122253.1 | 1.02421 | 0.71203 | *clpP* |
| WP_000130305.1 | 1.01767 | 0.73985 | *clpX* |
| WP_001295325.1 | 1.05679 | 0.58207 | *lon* |
| WP_001043542.1 | 0.96782 | 0.96701 | *hupB* |
| WP_000969372.1 | 0.95293 | 0.94295 | *ppiD* |
| WP_001194534.1 | 1.32241 | 0.23545 | *DXE50_04375* |
| WP_000817220.1 | 0.8183 | 0.24674 | *queC* |
| WP_000884589.1 | 0.66999 | 0.05291 | *DXE50_04395* |
| WP_000075876.1 | 0.90277 | 0.45051 | *tesB* |
| WP_000779842.1 | 0.92848 | 0.57496 | *KXD89_16535* |
| WP_000136192.1 | 0.85119 | 0.24581 | *DXE50_04450* |
| WP_000102564.1 | 0.88623 | 0.48285 | *DXE50_04455* |
| WP_001291435.1 | 1.22386 | 0.39226 | *hha* |
| WP_001132469.1 | 0.86583 | 0.37235 | *DXE50_04470* |
| WP_001295324.1 | 0.9309 | 0.78585 | *acrA* |
| WP_000101737.1 | 1.18917 | 0.56878 | *acrR* |
| WP_000177732.1 | 1.04514 | 0.85632 | *mscK* |
| WP_000051153.1 | 1.41717 | 0.13831 | *rsmS* |
| WP_000127356.1 | 1.08228 | 0.49127 | *apt* |
| WP_000122013.1 | 1.18608 | 0.3497 | *dnaX* |
| WP_000467098.1 | 0.86183 | 0.35151 | *ybaB* |
| WP_001195025.1 | 0.73702 | 0.1384 | *recR* |
| WP_000678201.1 | 0.87125 | 0.40167 | *htpG* |
| WP_001220233.1 | 0.9193 | 0.70463 | *adk* |
| WP_001250103.1 | 1.31181 | 0.36571 | *hemH* |
| WP_000671574.1 | 0.90065 | 0.44081 | *gsk* |
| WP_000546237.1 | 0.79217 | 0.17891 | *A1UI_00561* |
| WP_001251608.1 | 1.14813 | 0.65265 | *A1UI_00562* |
| WP_000771748.1 | 1.31695 | 0.06813 | *A13A_00461* |
| WP_000186631.1 | 1.0167 | 0.94963 | *ybaK* |
| WP_000806442.1 | 1.19667 | 0.55433 | *A13A_00465* |
| WP_000083955.1 | 1.39245 | 0.06401 | *copA* |
| WP_001026747.1 | 0.94472 | 0.76423 | *cueR* |
| WP_000904502.1 | 1.66928 | 0.08429 | *DXE50_04660* |
| WP_001300573.1 | 0.8644 | 0.36483 | *cnoX* |
| WP_000148959.1 | 1.26231 | 0.43929 | *ybbO* |
| WP_001295836.1 | 1.08667 | 0.72819 | *ECMP0215528_0556* |
| WP_001110573.1 | 1.10276 | 0.60391 | *DXE50_04690* |
| WP_000141275.1 | 0.99616 | 0.94 | *allR* |
| WP_001333621.1 | 1.26855 | 0.42941 | *glxK* |
| WP_000815571.1 | 0.98379 | 0.91472 | *purK* |
| WP_001295318.1 | 0.97671 | 0.88462 | *purE* |

| | | | |
|---|---|---|---|
| WP_000255997.1 | 0.95387 | 0.94966 | *ppiB* |
| WP_000912385.1 | 0.83895 | 0.24523 | *cysS* |
| WP_000190288.1 | 0.95974 | 0.73966 | *ybcJ* |
| WP_000729160.1 | 0.90849 | 0.47709 | *folD* |
| WP_000662357.1 | 0.85891 | 0.4513 | *nfrA* |
| WP_000770953.1 | 1.08868 | 0.78811 | *cusR* |
| WP_000786319.1 | 1.04275 | 0.90307 | *pheP* |
| WP_001153148.1 | 0.79327 | 0.25778 | *DXE50_05095* |
| WP_000351487.1 | 0.98597 | 0.88163 | *nfsB* |
| WP_001130654.1 | 1.65611 | 0.08926 | *KXD89_15900* |
| WP_001034943.1 | 0.83456 | 0.19473 | *fepA* |
| WP_000026812.1 | 0.89234 | 0.50773 | *entE* |
| WP_001007138.1 | 0.95315 | 0.77574 | *A1UI_00658* |
| WP_000347651.1 | 0.95155 | 0.76835 | *A1UI_00659* |
| WP_001043156.1 | 1.28254 | 0.16694 | *A1UI_00661* |
| WP_000913829.1 | 0.96879 | 0.84797 | *dsbG* |
| WP_000052796.1 | 0.95778 | 0.97761 | *ahpC* |
| WP_000887629.1 | 0.89024 | 0.5131 | *ahpF* |
| WP_000089731.1 | 1.07847 | 0.69542 | *rnk* |
| WP_000643397.1 | 0.85403 | 0.36092 | *A1UI_00676* |
| WP_000062457.1 | 0.6866 | 0.0689 | *citG* |
| WP_000034825.1 | 0.77316 | 0.06274 | *cspE* |
| WP_000042632.1 | 0.8012 | 0.1145 | *lipA* |
| WP_000284027.1 | 0.72996 | 0.06993 | *lipB* |
| WP_000850550.1 | 1.02976 | 0.8987 | *ybeD* |
| WP_001092082.1 | 1.0264 | 0.70283 | *dacA* |
| WP_001231430.1 | 1.11063 | 0.57586 | *rlpA* |
| WP_000131719.1 | 0.91228 | 0.64102 | *mrdB* |
| WP_000776191.1 | 1.14993 | 0.55567 | *mrdA* |
| WP_000776104.1 | 1.04672 | 0.85127 | *rlmH* |
| WP_001161664.1 | 0.89358 | 0.40928 | *rsfS* |
| WP_000620535.1 | 0.92552 | 0.69086 | *EL76_3432* |
| WP_001269673.1 | 0.99825 | 0.98388 | *lptE* |
| WP_001340834.1 | 0.85059 | 0.29644 | *leuS* |
| WP_001044880.1 | 1.33978 | 0.05441 | *DXE50_05445* |
| WP_001207520.1 | 0.95386 | 0.70792 | *rihA* |
| WP_001177086.1 | 1.26484 | 0.19269 | *gltI* |
| WP_000853021.1 | 0.8868 | 0.48517 | *lnt* |
| WP_001278605.1 | 1.15629 | 0.42967 | *corC* |
| WP_000084469.1 | 0.822 | 0.33589 | *ybeY* |
| WP_001018040.1 | 0.89647 | 0.55225 | *ybeZ* |
| WP_000162740.1 | 0.90816 | 0.62878 | *miaB* |
| WP_000184541.1 | 0.76034 | 0.11435 | *A1UI_00729* |
| WP_000337077.1 | 0.78125 | 0.0764 | *WG3_00959* |
| WP_000153129.1 | 1.02039 | 0.94019 | *nagD* |
| WP_000187594.1 | 1.119 | 0.54691 | *nagC* |
| WP_001237072.1 | 0.78689 | 0.08864 | *nagB* |
| WP_038432907.1 | 0.95675 | 0.72348 | - |

| | | | |
|---|---|---|---|
| WP_001287154.1 | 0.91155 | 0.65155 | *glnS* |
| WP_000131702.1 | 1.0407 | 0.64438 | *fur* |
| WP_001018618.1 | 0.92645 | 0.75445 | *fldA* |
| WP_001300829.1 | 0.99677 | 0.96438 | *DXE50_05640* |
| WP_000773288.1 | 0.83029 | 0.18281 | *ybfF* |
| WP_000848387.1 | 0.92375 | 0.55113 | *seqA* |
| WP_001297249.1 | 1.06275 | 0.55995 | *A1UI_00749* |
| WP_001310640.1 | 1.18424 | 0.57844 | *A1UI_00756* |
| WP_000087939.1 | 0.9586 | 0.81767 | *kdpB* |
| WP_000424924.1 | 1.13698 | 0.67682 | *DXE50_05690* |
| WP_001053305.1 | 1.18345 | 0.57999 | *ECMP0215528_0806* |
| WP_000798871.1 | 1.12976 | 0.34885 | *DXE50_05710* |
| WP_001188343.1 | 0.87204 | 0.42712 | *ECMP0215528_0810* |
| WP_000912724.1 | 1.08753 | 0.72565 | *pxpC* |
| WP_000785834.1 | 0.99291 | 0.84971 | *gltA* |
| AIN31197.1 | 0.84691 | 0.23193 | - |
| WP_000775540.1 | 0.98519 | 0.88526 | *sdhA* |
| WP_001235254.1 | 1.03229 | 0.67843 | *A311_01246* |
| WP_001181473.1 | 0.90967 | 0.63888 | *sucA* |
| WP_000099823.1 | 0.9641 | 0.98479 | *WG3_01000* |
| WP_001048602.1 | 1.04163 | 0.6407 | *sucC* |
| WP_000025458.1 | 1.04193 | 0.63949 | *sucD* |
| WP_000509902.1 | 0.81552 | 0.31731 | *mngR* |
| WP_000884361.1 | 1.0365 | 0.66128 | *O3K_17975* |
| WP_000034602.1 | 1.01232 | 0.98342 | *ybgE* |
| WP_001098384.1 | 0.86827 | 0.4829 | *ybgC* |
| WP_000131314.1 | 0.93008 | 0.58309 | *tolQ* |
| WP_000090097.1 | 0.97218 | 0.87006 | *tolR* |
| WP_000030637.1 | 0.9279 | 0.57199 | *tolA* |
| WP_001295307.1 | 1.07866 | 0.50354 | *tolB* |
| WP_001295306.1 | 1.00588 | 0.79132 | *pal* |
| WP_000097571.1 | 0.96876 | 0.78892 | *cpoB* |
| WP_000784351.1 | 1.52436 | 0.15659 | *DXE50_05990* |
| WP_001109196.1 | 0.97545 | 0.8257 | *O3K_17900* |
| WP_001295305.1 | 0.99301 | 0.84925 | *gpmA* |
| WP_000931389.1 | 0.96119 | 0.99871 | *galM* |
| WP_000191497.1 | 0.7629 | 0.05532 | *A1UI_00817* |
| WP_001265438.1 | 0.84068 | 0.25245 | *galE* |
| WP_000096869.1 | 0.90315 | 0.55292 | *G711_00399* |
| WP_001147439.1 | 0.90376 | 0.55549 | *modE* |
| WP_000101984.1 | 1.0234 | 0.92679 | *A1UI_00822* |
| WP_000891692.1 | 1.6354 | 0.09764 | *modC* |
| WP_001300666.1 | 0.94489 | 0.66001 | *A1UI_00825* |
| WP_000815435.1 | 1.10403 | 0.42169 | *pgl* |
| WP_001091569.1 | 0.84252 | 0.21824 | *ECMP0215528_0877* |
| WP_001295303.1 | 1.77385 | 0.05317 | *bioA* |
| WP_000951213.1 | 0.72695 | 0.1218 | *bioB* |
| WP_000044843.1 | 1.00058 | 0.97885 | *bioD* |

| | | | |
|---|---|---|---|
| WP_000042533.1 | 1.05071 | 0.83855 | *uvrB* |
| WP_001295302.1 | 1.05317 | 0.83075 | *ECMP0215528_0925* |
| WP_000084639.1 | 1.25753 | 0.20425 | *moaB* |
| WP_000080885.1 | 1.10881 | 0.66449 | *moaC* |
| WP_000852296.1 | 0.99276 | 0.9587 | *moaE* |
| WP_001393495.1 | 0.94452 | 0.65808 | *hlyD* |
| WP_001296991.1 | 1.48208 | 0.1866 | *cecR* |
| WP_000007101.1 | 0.95515 | 0.78492 | *rhlE* |
| WP_000386551.1 | 1.01187 | 0.97839 | *ECMP0215528_0948* |
| WP_000443530.1 | 1.13464 | 0.49541 | *ECMP0215528_0949* |
| WP_001295299.1 | 1.17578 | 0.4938 | *rlmF* |
| WP_000569083.1 | 0.86737 | 0.30271 | *ECMP0215528_0957* |
| WP_001159065.1 | 0.92548 | 0.65015 | *glnP* |
| WP_000843866.1 | 0.9164 | 0.68465 | *glnH* |
| WP_000091016.1 | 0.84693 | 0.33635 | *DXE50_06600* |
| WP_001056384.1 | 0.91727 | 0.51915 | *ldtB* |
| WP_000961458.1 | 0.88929 | 0.5072 | *ECMP0215528_0968* |
| WP_000168797.1 | 1.15477 | 0.54368 | *ECMP0215528_0969* |
| WP_000114272.1 | 0.87368 | 0.32678 | *ybiV* |
| WP_001336208.1 | 1.23195 | 0.48996 | *fsa* |
| WP_000829217.1 | 1.03727 | 0.91732 | *moeB* |
| WP_000397340.1 | 0.93717 | 0.83032 | *moeA* |
| WP_000513781.1 | 1.02208 | 0.95734 | *iaaA* |
| WP_001301279.1 | 1.16556 | 0.61604 | *gsiA* |
| WP_000090140.1 | 1.17346 | 0.49913 | *gsiB* |
| WP_000049367.1 | 0.80122 | 0.11972 | *rimO* |
| WP_000555031.1 | 1.13945 | 0.67142 | *A1UI_00905* |
| WP_001295292.1 | 1.03919 | 0.8576 | *DXE50_06705* |
| WP_000450121.1 | 0.95999 | 0.80726 | *deoR* |
| WP_000023565.1 | 1.17083 | 0.60526 | *ybjI* |
| WP_001024876.1 | 1.04328 | 0.90171 | *ybjL* |
| WP_001195240.1 | 1.10207 | 0.75621 | *EKN05_012210* |
| WP_000189159.1 | 0.94833 | 0.67828 | *nfsA* |
| WP_000203025.1 | 0.908 | 0.62508 | *DXE50_06785* |
| WP_000126072.1 | 1.72773 | 0.06523 | *O3K_17075* |
| WP_001295339.1 | 1.11397 | 0.72853 | *DXE50_06835* |
| WP_000756569.1 | 1.08604 | 0.47872 | *ECMP0215528_1029* |
| WP_000027205.1 | 1.00076 | 0.99533 | *artP* |
| WP_001270734.1 | 1.05599 | 0.78634 | *O3K_17020* |
| WP_001160737.1 | 1.27037 | 0.31016 | *ybjQ* |
| WP_001252135.1 | 0.94104 | 0.75007 | *amiD* |
| WP_001338420.1 | 0.95642 | 0.80928 | *ECMP0215528_1034* |
| WP_000566376.1 | 0.91958 | 0.53048 | *ltaE* |
| WP_000491142.1 | 0.80194 | 0.28019 | *ECMP0215528_1040* |
| WP_000599802.1 | 0.68714 | 0.06947 | *KXD89_14410* |
| WP_000746442.1 | 0.74895 | 0.09585 | *macA* |
| WP_000410785.1 | 1.10462 | 0.59719 | *cspD* |
| WP_000520781.1 | 0.82263 | 0.33773 | *clpS* |

| | | | |
|---|---|---|---|
| WP_000934041.1 | 1.3153 | 0.06923 | *clpA* |
| WP_001040187.1 | 0.89706 | 0.55604 | *infA* |
| WP_001202177.1 | 1.06197 | 0.80312 | *cydC* |
| WP_001043598.1 | 1.05219 | 0.83386 | *cydD* |
| WP_000537418.1 | 0.96859 | 0.96335 | *trxB* |
| WP_000228473.1 | 1.00353 | 0.80176 | *GA0061070_10177* |
| WP_000076967.1 | 0.95066 | 0.76427 | *ftsK* |
| WP_000067755.1 | 0.94732 | 0.77421 | *DXE50_07020* |
| WP_000886683.1 | 0.91375 | 0.66651 | *serS* |
| WP_000213098.1 | 1.00613 | 0.99987 | *dmsB* |
| WP_000111043.1 | 0.88707 | 0.3812 | *pflA* |
| WP_001292822.1 | 1.04751 | 0.61756 | *A13A_00842* |
| WP_000642546.1 | 1.46191 | 0.10632 | *focA* |
| WP_001295344.1 | 0.88681 | 0.49203 | *DXE50_07080* |
| WP_000642849.1 | 1.44254 | 0.21926 | *ECMP0215528_1073* |
| WP_000057138.1 | 0.93545 | 0.81808 | *serC* |
| WP_000445231.1 | 0.94309 | 0.65049 | *aroA* |
| WP_000125016.1 | 0.86821 | 0.30586 | *cmk* |
| WP_000140327.1 | 0.87299 | 0.41132 | *rpsA* |
| WP_000167336.1 | 1.33868 | 0.10374 | *ihfB* |
| WP_000551270.1 | 1.0076 | 0.98025 | *msbA* |
| WP_000570539.1 | 1.03624 | 0.92 | *lpxK* |
| WP_000350058.1 | 0.82909 | 0.27869 | *ycaR* |
| WP_000011603.1 | 1.09359 | 0.6376 | *kdsB* |
| WP_001298300.1 | 0.89533 | 0.57853 | *cmoM* |
| WP_001288850.1 | 0.90908 | 0.57823 | *mukF* |
| WP_001295347.1 | 0.95492 | 0.78386 | *mukE* |
| WP_000572698.1 | 0.97589 | 0.92874 | *mukB* |
| WP_001109486.1 | 1.28895 | 0.15836 | *A1UI_01038* |
| WP_000462687.1 | 0.94248 | 0.86824 | *DXE50_07195* |
| WP_000117881.1 | 0.90393 | 0.60066 | *asnS* |
| WP_001307697.1 | 0.90363 | 0.45446 | *pncB* |
| WP_000193841.1 | 1.00841 | 0.78011 | *A1UI_01043* |
| WP_001295352.1 | 0.8006 | 0.11817 | *pyrD* |
| WP_000212426.1 | 1.02111 | 0.9349 | *A1UI_01060* |
| WP_001086539.1 | 1.04951 | 0.81351 | *rlmL* |
| WP_000053099.1 | 0.94479 | 0.65948 | *uup* |
| WP_000445533.1 | 1.05773 | 0.81638 | *O3K_16585* |
| WP_000759120.1 | 1.35644 | 0.19532 | *pqiC* |
| WP_000227927.1 | 0.91701 | 0.68885 | *fabA* |
| WP_000156518.1 | 0.97561 | 0.87955 | *A1UI_01068* |
| WP_000877161.1 | 0.87743 | 0.5146 | *matP* |
| WP_000750416.1 | 1.20989 | 0.1824 | *ompA* |
| WP_001261235.1 | 0.83298 | 0.29073 | *DXE50_07360* |
| WP_001295354.1 | 1.11555 | 0.38783 | *A1UI_01075* |
| WP_000424181.1 | 0.83089 | 0.21327 | *mgsA* |
| WP_001301418.1 | 1.16784 | 0.39724 | *KXD89_13960* |
| WP_000116297.1 | 0.98925 | 0.90188 | *rlmI* |

| | | | |
|---|---|---|---|
| WP_000904442.1 | 0.86553 | 0.4026 | *tusE* |
| WP_000375136.1 | 0.92143 | 0.67537 | *yccA* |
| WP_001300464.1 | 0.82791 | 0.3533 | *G711_00113* |
| WP_001044279.1 | 1.15063 | 0.44623 | *agp* |
| WP_001143120.1 | 1.41568 | 0.05157 | *DXE50_07910* |
| WP_000191701.1 | 1.13272 | 0.68622 | *rutR* |
| WP_001326840.1 | 0.97377 | 0.93876 | *putA* |
| WP_001018479.1 | 0.84104 | 0.21374 | *putP* |
| WP_000154398.1 | 0.72529 | 0.06444 | *EKN05_011205* |
| WP_000533522.1 | 1.32347 | 0.34993 | *phoH* |
| WP_000351317.1 | 1.08398 | 0.48554 | *ghrA* |
| WP_000283667.1 | 1.17907 | 0.36745 | *ycdX* |
| WP_001001921.1 | 1.05295 | 0.79902 | *A13A_00953* |
| WP_001189321.1 | 1.44676 | 0.11631 | *csgG* |
| WP_000857405.1 | 1.26374 | 0.32094 | *ymdB* |
| WP_001343212.1 | 0.94304 | 0.87224 | *mdoG* |
| WP_001295445.1 | 0.93603 | 0.6975 | *mdoH* |
| WP_000180056.1 | 1.30993 | 0.13287 | *ECMP0215528_1226* |
| WP_000183364.1 | 0.93462 | 0.69116 | *lpxL* |
| WP_000749261.1 | 1.04732 | 0.84935 | *yceI* |
| WP_000872833.1 | 1.03792 | 0.86313 | *solA* |
| WP_001217754.1 | 0.97368 | 0.87584 | *dinI_4* |
| WP_000126534.1 | 0.85564 | 0.32051 | *pyrC* |
| WP_001295443.1 | 0.79767 | 0.19206 | *DXE50_08675* |
| WP_000780912.1 | 1.1165 | 0.38512 | *O3K_15240* |
| WP_000468186.1 | 1.13349 | 0.59765 | *rimJ* |
| WP_000877107.1 | 1.16762 | 0.39784 | *yceH* |
| WP_001050683.1 | 0.84721 | 0.41308 | *murJ* |
| WP_000827360.1 | 0.9269 | 0.75762 | *rne* |
| WP_000846343.1 | 1.18057 | 0.3636 | *rluC* |
| WP_001125202.1 | 1.10862 | 0.58293 | *EKN05_010785* |
| WP_001174481.1 | 0.92355 | 0.64154 | *yceD* |
| WP_000197597.1 | 1.23249 | 0.48902 | *plsX* |
| WP_000288132.1 | 1.03686 | 0.65979 | *fabH* |
| WP_000191372.1 | 1.06602 | 0.54804 | *fabD* |
| WP_001008535.1 | 1.05165 | 0.60155 | *fabG* |
| WP_000103754.1 | 0.91807 | 0.69613 | *acpP* |
| WP_000044679.1 | 0.93509 | 0.8155 | *fabF* |
| WP_000478681.1 | 1.12964 | 0.69305 | *pabC* |
| WP_000756827.1 | 0.91903 | 0.66631 | *yceG* |
| WP_001257000.1 | 0.92772 | 0.66012 | *tmk* |
| WP_001267956.1 | 1.17002 | 0.60691 | *holB* |
| WP_000480245.1 | 1.07769 | 0.75497 | *ECMP0215528_1276* |
| WP_000475719.1 | 0.99306 | 0.84903 | *ptsG* |
| WP_000807125.1 | 1.01056 | 0.98428 | *hinT* |
| WP_000164439.1 | 1.08518 | 0.66943 | *lpoB* |
| WP_000529320.1 | 0.99875 | 0.98616 | *nagZ* |
| WP_000587933.1 | 1.23034 | 0.25239 | *ycfP* |

| | | | |
|---|---|---|---|
| WP_000211045.1 | 0.94078 | 0.85607 | *DXE50_08910* |
| WP_001043459.1 | 0.8499 | 0.34653 | *DXE50_08915* |
| WP_001336528.1 | 1.18642 | 0.57416 | *comR* |
| WP_001115094.1 | 1.01209 | 0.76402 | *mfd* |
| WP_000284714.1 | 1.06666 | 0.8422 | *lolC* |
| WP_001033694.1 | 1.10629 | 0.67156 | *lolD* |
| WP_001251348.1 | 0.96166 | 0.82949 | *HMPREF1620_01675* |
| WP_000291270.1 | 1.17005 | 0.60685 | *nagK* |
| AIN31583.1 | 1.02465 | 0.92317 | - |
| WP_000759317.1 | 0.98933 | 0.90234 | *potD* |
| WP_000580316.1 | 1.02238 | 0.93069 | *potC* |
| AIN31588.1 | 1.00257 | 0.98638 | - |
| WP_000531594.1 | 1.00948 | 0.98917 | *potA* |
| WP_000359434.1 | 1.30417 | 0.13949 | *pepT* |
| WP_000456506.1 | 1.02436 | 0.92411 | *AB05_1317* |
| WP_000423742.1 | 0.96317 | 0.98921 | *purB* |
| WP_001297479.1 | 1.06422 | 0.79613 | *hflD* |
| WP_001297484.1 | 0.93794 | 0.62356 | *trmU* |
| WP_000476093.1 | 1.32076 | 0.35355 | *nudJ* |
| WP_000444484.1 | 0.94233 | 0.86717 | *icd* |
| WP_000848748.1 | 0.82654 | 0.34922 | *A311_03318* |
| WP_000888772.1 | 0.67962 | 0.0618 | *ariR* |
| WP_001185665.1 | 0.91918 | 0.52851 | *minE* |
| WP_000101055.1 | 1.10434 | 0.42077 | *minD* |
| WP_001301105.1 | 1.08877 | 0.72199 | *minC* |
| WP_001056840.1 | 0.73219 | 0.07267 | *HMPREF1602_05104* |
| WP_000695215.1 | 1.04002 | 0.87281 | *KXD89_12850* |
| WP_001295992.1 | 0.94612 | 0.66652 | *ycgL* |
| WP_000943459.1 | 0.96616 | 0.84684 | *dsbB* |
| WP_000406391.1 | 0.76015 | 0.11403 | *nhaB* |
| WP_000234823.1 | 0.8679 | 0.41144 | *fadR* |
| WP_001266908.1 | 0.93618 | 0.82324 | *dadA* |
| WP_000197881.1 | 1.13371 | 0.49838 | *dadX* |
| WP_000051560.1 | 1.04819 | 0.84658 | *A1UI_01327* |
| WP_001301104.1 | 0.85682 | 0.37084 | *emtA* |
| WP_000841714.1 | 1.34023 | 0.10234 | *treA* |
| WP_001301101.1 | 0.97338 | 0.81432 | *A1UI_01333* |
| WP_000059411.1 | 0.98689 | 0.88886 | *EL76_2821* |
| WP_000733715.1 | 1.13044 | 0.34707 | *A1UI_01335* |
| WP_000505866.1 | 0.93565 | 0.81949 | *ychF* |
| WP_000152933.1 | 0.87289 | 0.32372 | *pth* |
| WP_000823885.1 | 1.11267 | 0.65374 | *ychH* |
| WP_001033352.1 | 0.94965 | 0.75962 | *dauA* |
| WP_001298109.1 | 0.96093 | 1 | *prs* |
| WP_001260332.1 | 0.93604 | 0.69755 | *ispE* |
| WP_001130692.1 | 1.06157 | 0.76327 | *lolB* |
| WP_001299679.1 | 1.10918 | 0.73959 | *hemA* |
| WP_000804726.1 | 1.06009 | 0.76936 | *prfA* |

| | | | |
|---|---|---|---|
| WP_000456467.1 | 0.95669 | 0.8103 | *prmC* |
| WP_000811065.1 | 1.06646 | 0.54647 | *kdsA* |
| WP_000063607.1 | 1.61491 | 0.10666 | *chaA* |
| WP_001146444.1 | 1.06637 | 0.84293 | *chaB* |
| WP_001169669.1 | 0.99515 | 0.93443 | *ychN* |
| WP_000070491.1 | 0.98183 | 0.86089 | *narL* |
| WP_000918073.1 | 1.20054 | 0.43932 | *narX* |
| WP_000019827.1 | 1.30333 | 0.26087 | *narK* |
| WP_000032939.1 | 1.21689 | 0.17169 | *KXD89_12605* |
| WP_000571681.1 | 1.33354 | 0.22161 | *narJ* |
| WP_001160108.1 | 1.41301 | 0.14168 | *narI* |
| WP_000555857.1 | 0.92352 | 0.54998 | *purU* |
| WP_001307143.1 | 0.97483 | 0.8803 | *ychJ* |
| WP_000193447.1 | 1.126 | 0.7012 | *rssB* |
| WP_001287378.1 | 1.08268 | 0.4899 | *DXE50_00785* |
| WP_000068077.1 | 0.97464 | 0.87507 | *tdk* |
| WP_000301651.1 | 1.0954 | 0.44841 | *adhE* |
| WP_001393463.1 | 1.08807 | 0.47202 | *oppA* |
| WP_000911112.1 | 1.03634 | 0.86997 | *oppB* |
| WP_000979661.1 | 1.12952 | 0.60809 | *O3K_14420* |
| WP_000110948.1 | 1.16084 | 0.41669 | *oppD* |
| WP_000994906.1 | 1.08554 | 0.48035 | *oppF* |
| WP_000214516.1 | 1.16786 | 0.61132 | *cls* |
| WP_001309467.1 | 0.8516 | 0.42725 | *DXE50_00840* |
| WP_001295624.1 | 1.31132 | 0.36638 | *A1UI_01400* |
| WP_000967595.1 | 0.8329 | 0.29051 | *DXE50_00850* |
| WP_000108160.1 | 0.88021 | 0.45887 | *yciA* |
| WP_000028545.1 | 0.78077 | 0.22732 | *yciC* |
| WP_000737226.1 | 1.09286 | 0.45651 | *ompW* |
| WP_001079505.1 | 1.48223 | 0.09415 | *A1UI_01407* |
| WP_000443067.1 | 1.15984 | 0.4195 | *trpA* |
| WP_000209520.1 | 1.12136 | 0.53893 | *trpB* |
| WP_000763511.1 | 0.91213 | 0.64046 | *trpD* |
| WP_001194582.1 | 1.49261 | 0.08843 | *A1UI_01414* |
| WP_001295575.1 | 1.37205 | 0.07711 | *DXE50_17760* |
| WP_001291217.1 | 0.99475 | 0.93222 | *WG3_01862* |
| WP_001278906.1 | 0.84014 | 0.31371 | *HMPREF1591_00308* |
| WP_000559286.1 | 0.96076 | 0.74524 | *KXD89_12335* |
| WP_000422045.1 | 0.99473 | 0.96776 | *sohB* |
| WP_001031530.1 | 1.02748 | 0.90874 | *DXE50_17730* |
| WP_001297122.1 | 1.04479 | 0.6282 | *topA* |
| WP_000776253.1 | 0.92983 | 0.66959 | *cysB* |
| WP_000099534.1 | 1.33864 | 0.05503 | *acnA* |
| WP_001176295.1 | 0.87123 | 0.31731 | *ribA* |
| WP_000876286.1 | 0.76527 | 0.19267 | *lapA* |
| WP_000891353.1 | 0.93791 | 0.73809 | *lapB* |
| WP_000176270.1 | 0.87706 | 0.43438 | *pyrF* |
| WP_001295580.1 | 0.96668 | 0.8382 | *ECMP0215528_1563* |

| | | | |
|---|---|---|---|
| WP_001088620.1 | 0.99243 | 0.95719 | *KXD89_12255* |
| WP_000484984.1 | 0.94914 | 0.91586 | *rnb* |
| WP_000506490.1 | 0.92278 | 0.72876 | *fabI* |
| WP_001146163.1 | 0.97332 | 0.87445 | *sapC* |
| WP_001250216.1 | 0.9719 | 0.86901 | *O3K_13860* |
| WP_001296746.1 | 0.86463 | 0.47053 | *puuA* |
| WP_001300506.1 | 1.30699 | 0.25583 | *A1UI_01451* |
| WP_001009090.1 | 0.87511 | 0.50652 | *A1UI_01453* |
| WP_000069229.1 | 0.92368 | 0.6839 | *EL76_2650* |
| WP_001301108.1 | 0.94807 | 0.77708 | *pspF* |
| WP_000511025.1 | 1.173 | 0.24843 | *pspA* |
| WP_000907387.1 | 0.96914 | 0.85833 | *pspC* |
| WP_000473109.1 | 0.93582 | 0.61253 | *pspE* |
| WP_000075375.1 | 0.91183 | 0.63934 | *A1UI_01474* |
| WP_000825775.1 | 0.89645 | 0.58261 | *A1UI_01475* |
| WP_000138728.1 | 1.46718 | 0.19836 | *ycjF* |
| WP_001300658.1 | 1.14548 | 0.4617 | *tyrR* |
| WP_000084387.1 | 0.91692 | 0.68823 | *tpx* |
| WP_001261211.1 | 1.10333 | 0.60182 | *A1UI_01479* |
| WP_000219122.1 | 0.95439 | 0.80142 | *mpaA* |
| WP_001300523.1 | 0.98617 | 0.92387 | *A1UI_01482* |
| WP_000683020.1 | 1.01819 | 0.95002 | *mppA* |
| WP_000559900.1 | 1.3276 | 0.34448 | *ynaI* |
| WP_001262123.1 | 1.27742 | 0.09941 | *uspE* |
| WP_000611911.1 | 0.77993 | 0.07775 | *fnr* |
| WP_000945011.1 | 0.8847 | 0.54027 | *DXE50_17365* |
| WP_000444929.1 | 0.71002 | 0.09704 | *abgA* |
| WP_001046829.1 | 1.2152 | 0.51978 | *smrA* |
| WP_000387388.1 | 1.07318 | 0.82597 | *zntB* |
| WP_000123737.1 | 0.98579 | 0.92241 | *dbpA* |
| WP_001157406.1 | 0.8635 | 0.28847 | *ttcA* |
| WP_000948459.1 | 0.95835 | 0.81673 | *C4A13_02150* |
| WP_001300461.1 | 1.20878 | 0.2967 | *A1UI_01500* |
| WP_000628244.1 | 0.93072 | 0.58634 | *nifJ* |
| WP_000762236.1 | 1.12099 | 0.37256 | *HMPREF1591_00398* |
| WP_000698145.1 | 0.9474 | 0.77451 | *EKN05_007005* |
| WP_000039887.1 | 1.07984 | 0.80959 | *A1UI_01524* |
| WP_001123464.1 | 1.38635 | 0.27439 | *EL76_2579* |
| WP_000097801.1 | 1.24619 | 0.35093 | *O3K_13465* |
| WP_000048950.1 | 0.77532 | 0.14232 | *azoR* |
| WP_000139543.1 | 1.02189 | 0.93352 | *hrpA* |
| WP_001027942.1 | 1.0885 | 0.65673 | *HGD79_04235* |
| WP_000115943.1 | 1.11616 | 0.38609 | *aldA* |
| WP_000428998.1 | 1.27057 | 0.42625 | *cybB* |
| WP_000414567.1 | 0.76413 | 0.19026 | *KXD89_11535* |
| WP_000013789.1 | 1.29571 | 0.27164 | *KXD89_11530* |
| WP_000375961.1 | 1.02853 | 0.91035 | *opgD* |
| WP_001296778.1 | 1.04301 | 0.84119 | *DXE50_16735* |

| | | | |
|---|---|---|---|
| WP_000140873.1 | 1.10872 | 0.74066 | *rimL* |
| WP_000586732.1 | 1.12798 | 0.61217 | *O3K_13350* |
| WP_001261013.1 | 1.19961 | 0.31731 | *DXE50_16710* |
| WP_001303492.1 | 0.91172 | 0.58967 | *CSW52_29105* |
| WP_000047424.1 | 1.29599 | 0.14938 | *A1UI_01562* |
| WP_000531462.1 | 1.35975 | 0.08611 | *CSW52_29020* |
| WP_000689363.1 | 0.7665 | 0.12535 | *KXD89_11385* |
| WP_000550695.1 | 0.79397 | 0.10084 | *KXD89_11380* |
| WP_001120143.1 | 0.801 | 0.27769 | *pptA* |
| WP_010723100.1 | 1.23982 | 0.23466 | *fdnG* |
| WP_001240582.1 | 1.42264 | 0.23755 | *fdxH* |
| WP_000045648.1 | 1.55727 | 0.05944 | *fdnI* |
| WP_000152305.1 | 1.34045 | 0.10214 | *osmC* |
| WP_000830550.1 | 0.84815 | 0.34052 | *EL76_2497* |
| WP_000350395.1 | 0.87273 | 0.49825 | *EKN05_007410* |
| WP_001244968.1 | 0.99212 | 0.95575 | *KXD89_11170* |
| WP_000832437.1 | 1.07902 | 0.81158 | *KXD89_11165* |
| WP_001125439.1 | 0.92103 | 0.67387 | *A1UI_01641* |
| WP_000154342.1 | 1.58929 | 0.11903 | *lsrR* |
| WP_000172465.1 | 1.3343 | 0.2207 | *lsrB* |
| WP_000774165.1 | 1.21046 | 0.29304 | *lsrF* |
| WP_000558527.1 | 1.34641 | 0.32055 | *lsrG* |
| WP_001286597.1 | 1.41424 | 0.14068 | *tam* |
| WP_000156615.1 | 1.0954 | 0.772 | *sad* |
| WP_000210799.1 | 0.84195 | 0.39635 | *ydeA* |
| WP_000671737.1 | 1.29327 | 0.27517 | *A311_02188* |
| WP_000210373.1 | 1.0102 | 0.77227 | *dcp* |
| WP_000636571.1 | 0.97181 | 0.94803 | *ydfG* |
| WP_000215549.1 | 1.11046 | 0.73662 | *DXE50_16075* |
| WP_001019606.1 | 0.85056 | 0.24374 | *WG3_02248* |
| WP_000534858.1 | 0.81405 | 0.31317 | *relB* |
| WP_000448564.1 | 1.28009 | 0.41159 | *WG3_02272* |
| WP_001295394.1 | 1.49196 | 0.17915 | *rspA* |
| WP_001340362.1 | 1.0814 | 0.80576 | *KXD89_10665* |
| WP_000148710.1 | 1.0834 | 0.73789 | *dmsD* |
| WP_000919231.1 | 1.45354 | 0.20969 | *bioD* |
| WP_000225262.1 | 0.99102 | 0.94244 | *mlc* |
| WP_000014036.1 | 1.0156 | 0.96158 | *pntB* |
| WP_001300486.1 | 1.02805 | 0.90621 | *A1UI_01794* |
| WP_000769303.1 | 1.04693 | 0.6198 | *KXD89_10570* |
| WP_000520804.1 | 1.05582 | 0.86954 | *folM* |
| WP_000135181.1 | 1.34011 | 0.32841 | *tus* |
| WP_001099085.1 | 1.04277 | 0.63616 | *fumC* |
| WP_000066639.1 | 0.93972 | 0.84848 | *DXE50_15495* |
| WP_001170664.1 | 0.87534 | 0.42458 | *A1UI_01804* |
| WP_000945878.1 | 1.01378 | 0.95941 | |
| WP_000969092.1 | 1.67658 | 0.08165 | *DXE50_15465* |
| WP_000125583.1 | 0.83075 | 0.3618 | *malX* |

| | | | |
|---|---|---|---|
| WP_000567490.1 | 0.85329 | 0.30918 | *add* |
| WP_001282550.1 | 0.94237 | 0.72632 | *KXD89_10470* |
| WP_000217950.1 | 0.83541 | 0.29843 | *ydgT* |
| WP_000991805.1 | 0.82192 | 0.33565 | *rsxB* |
| WP_000915778.1 | 1.09829 | 0.76515 | *rsxC* |
| WP_000920784.1 | 1.18974 | 0.56768 | *rsxG* |
| WP_000100932.1 | 0.78608 | 0.16504 | *dtpA* |
| WP_000765749.1 | 1.21877 | 0.1689 | *gstA* |
| WP_001307229.1 | 1.02102 | 0.93735 | *pdxY* |
| WP_001295400.1 | 1.07831 | 0.50475 | *tyrS* |
| WP_001282319.1 | 1.13502 | 0.49421 | *pdxH* |
| WP_000835039.1 | 1.27406 | 0.30428 | *anmK* |
| WP_001296937.1 | 1.25513 | 0.33538 | *sodC* |
| WP_000250656.1 | 1.07163 | 0.72251 | *DXE50_15315* |
| WP_000093589.1 | 0.94541 | 0.66275 | *nemA* |
| WP_001237796.1 | 0.90862 | 0.47768 | *gloA* |
| WP_001282283.1 | 0.98859 | 0.93953 | *rnt* |
| WP_000108172.1 | 0.77429 | 0.06452 | *DXE50_15280* |
| WP_000007283.1 | 1.05264 | 0.59779 | *sodB* |
| WP_000190982.1 | 1.06712 | 0.74062 | *purR* |
| WP_000098896.1 | 1.27481 | 0.17781 | *DXE50_15240* |
| WP_000493947.1 | 1.05324 | 0.79781 | *ribE* |
| WP_001174940.1 | 1.27293 | 0.42258 | *mdtK* |
| WP_000534271.1 | 0.98002 | 0.85091 | *KXD89_10260* |
| WP_000212657.1 | 1.13182 | 0.60202 | *EKN05_007655* |
| WP_000716950.1 | 1.35804 | 0.30647 | *KXD89_10240* |
| WP_000528342.1 | 1.15237 | 0.64361 | *fumD* |
| WP_001295403.1 | 0.99749 | 0.82888 | *DXE50_15165* |
| WP_000648420.1 | 0.83847 | 0.24326 | *lpp* |
| WP_000817105.1 | 1.55418 | 0.06059 | *ldtE* |
| WP_001196530.1 | 1.04496 | 0.89736 | *sufE* |
| WP_000907979.1 | 1.35152 | 0.3143 | *sufD* |
| WP_000948863.1 | 1.51312 | 0.16411 | *sufC* |
| WP_000089364.1 | 1.01095 | 0.98711 | *sufB* |
| WP_000367160.1 | 1.41625 | 0.13904 | *sufA* |
| WP_001296104.1 | 1.13743 | 0.67583 | *DXE50_15120* |
| WP_000637982.1 | 0.90944 | 0.63043 | *ydiI* |
| WP_000613008.1 | 1.0556 | 0.82305 | *KXD89_10145* |
| WP_000248636.1 | 0.97269 | 0.87203 | *ydiK* |
| WP_000860201.1 | 0.98371 | 0.91698 | *aroD* |
| WP_000368046.1 | 0.90964 | 0.4825 | *ppsR* |
| WP_001082229.1 | 1.02791 | 0.90684 | *aroH* |
| WP_001229265.1 | 1.01487 | 0.75194 | *ihfA* |
| WP_000672380.1 | 0.99358 | 0.84666 | *pheT* |
| WP_000018596.1 | 0.96551 | 0.97804 | *pheS* |
| pdb\|6H58\|QQ | 0.86159 | 0.35025 | - |
| pdb\|6H58\|33 | 1.31814 | 0.06735 | - |
| WP_001700733.1 | 1.12747 | 0.35494 | *infC* |

| | | | |
|---|---|---|---|
| WP_001144202.1 | 0.94079 | 0.85613 | *thrS* |
| WP_000251727.1 | 1.35172 | 0.09249 | *EL76_2143* |
| WP_000146159.1 | 1.54989 | 0.06223 | *ghoS* |
| WP_000106833.1 | 0.8861 | 0.37712 | *A1UI_01927* |
| WP_001010696.1 | 1.02013 | 0.93817 | *A1UI_01929* |
| WP_000412169.1 | 1.11454 | 0.72723 | *chbB* |
| WP_001039044.1 | 1.15999 | 0.27588 | *osmE* |
| WP_000175026.1 | 1.04662 | 0.62104 | *nadE* |
| WP_001228984.1 | 0.89819 | 0.42969 | *A1UI_01943* |
| WP_000368506.1 | 0.96843 | 0.85562 | *astE* |
| WP_000177206.1 | 1.38023 | 0.17091 | *astD* |
| WP_000673918.1 | 0.91575 | 0.51177 | *xth* |
| WP_000939212.1 | 1.41773 | 0.13786 | *A1UI_01951* |
| WP_001215339.1 | 1.44566 | 0.21651 | *A1UI_01954* |
| WP_001350515.1 | 1.24275 | 0.35707 | *ynjE* |
| WP_000373021.1 | 1.00708 | 1 | *gdhA* |
| WP_001235800.1 | 1.05452 | 0.87283 | *topB* |
| WP_001295485.1 | 1.02626 | 0.91412 | *selD* |
| WP_000339283.1 | 1.11925 | 0.54607 | *O3K_11135* |
| WP_001259810.1 | 1.05252 | 0.80085 | *sppA* |
| WP_001170162.1 | 0.97494 | 0.87642 | *DXE50_14700* |
| WP_001135066.1 | 1.07654 | 0.81768 | *KXD89_09735* |
| WP_000719096.1 | 0.8888 | 0.49328 | *HMPREF1591_01708* |
| WP_000645221.1 | 0.78431 | 0.23574 | *KXD89_09695* |
| WP_001300415.1 | 1.03745 | 0.88117 | *KXD89_09690* |
| WP_001284618.1 | 1.11407 | 0.56386 | *msrB* |
| WP_000153502.1 | 0.87617 | 0.42929 | *gapA* |
| WP_001335909.1 | 0.83791 | 0.20439 | *A1UI_01981* |
| WP_000163771.1 | 0.97313 | 0.94177 | *DXE50_14620* |
| WP_000219687.1 | 1.4224 | 0.23778 | *yeaH* |
| WP_000766132.1 | 0.96344 | 0.83635 | *O3K_11030* |
| WP_000138043.1 | 1.02932 | 0.90776 | *A311_02408* |
| WP_001326526.1 | 1.49173 | 0.0889 | *KXD89_09615* |
| WP_000691930.1 | 0.86522 | 0.4725 | *ECMP0215528_2082* |
| WP_000939317.1 | 0.96462 | 0.8409 | *yeaR* |
| WP_000758422.1 | 1.11204 | 0.5709 | *fadD* |
| WP_000290576.1 | 1.13824 | 0.58532 | *yeaY* |
| WP_001220966.1 | 0.79285 | 0.18051 | *A1UI_02007* |
| WP_001295493.1 | 1.03046 | 0.89562 | *DXE50_14475* |
| WP_001111995.1 | 1.22272 | 0.50624 | *AC28_1962* |
| WP_000457334.1 | 1.34905 | 0.31731 | *yoaH* |
| WP_000854958.1 | 0.9631 | 0.75795 | *pabB* |
| WP_000456715.1 | 1.23719 | 0.48091 | *nudL* |
| WP_000624298.1 | 1.13593 | 0.33287 | *DXE50_14450* |
| WP_000150543.1 | 0.82754 | 0.2008 | *manX* |
| WP_000406926.1 | 0.94042 | 0.71742 | *DXE50_14430* |
| WP_000228655.1 | 0.76618 | 0.0592 | *DXE50_14425* |
| WP_000156255.1 | 0.92785 | 0.69971 | *yobD* |

| | | | |
|---|---|---|---|
| WP_001062678.1 | 1.0652 | 0.55102 | *HMPREF0880_03203* |
| WP_001006866.1 | 0.98197 | 0.90776 | *ECMP0215528_2119* |
| WP_001211011.1 | 1.18649 | 0.57402 | *DXE50_14385* |
| WP_001262203.1 | 0.92063 | 0.53568 | *kdgR* |
| WP_000984520.1 | 0.8267 | 0.17318 | *htpX* |
| WP_001055791.1 | 1.06503 | 0.55165 | *DXE50_14365* |
| WP_000431381.1 | 0.86862 | 0.3873 | *proQ* |
| WP_001043877.1 | 0.92829 | 0.7674 | *msrC* |
| WP_001326728.1 | 1.16706 | 0.51409 | *yebT* |
| WP_001352260.1 | 1.14973 | 0.55614 | *rsmF* |
| WP_001295499.1 | 1.60824 | 0.10976 | *DXE50_14335* |
| WP_000976476.1 | 0.89048 | 0.50013 | *EKN05_006800* |
| WP_000168747.1 | 1.04002 | 0.87281 | *yobA* |
| WP_000944256.1 | 0.75501 | 0.17164 | *exoX* |
| WP_000024757.1 | 1.49599 | 0.08664 | *KXD89_09330* |
| WP_000257738.1 | 0.72033 | 0.11166 | *yebG* |
| WP_000173484.1 | 1.06473 | 0.79457 | *purT* |
| WP_000800512.1 | 0.90473 | 0.60594 | *DXE50_14265* |
| WP_001069467.1 | 0.87316 | 0.32476 | *edd* |
| WP_000301727.1 | 0.91147 | 0.65106 | *zwf* |
| WP_001056706.1 | 0.93949 | 0.7132 | *O3K_10665* |
| WP_000091148.1 | 1.14216 | 0.31731 | *DXE50_14245* |
| WP_000448381.1 | 0.95637 | 0.79053 | *msbB* |
| WP_001300644.1 | 0.7926 | 0.09839 | *znuA* |
| WP_000202996.1 | 1.02201 | 0.95752 | *znuC* |
| WP_000568519.1 | 1.19071 | 0.46038 | *ruvB* |
| WP_000580323.1 | 1.01784 | 0.96863 | *ruvA* |
| WP_000907248.1 | 1.02207 | 0.93269 | *yebC* |
| WP_001258678.1 | 1.03993 | 0.64748 | *aspS* |
| WP_000639274.1 | 1.40903 | 0.25083 | *yecE* |
| WP_000019590.1 | 0.88012 | 0.35241 | *cmoA* |
| WP_000564725.1 | 0.81439 | 0.14276 | *cmoB* |
| WP_001185741.1 | 1.00617 | 1 | *cutC* |
| WP_001326725.1 | 1.57063 | 0.12887 | *GKR75_01495* |
| WP_001025322.1 | 0.91809 | 0.69628 | *argS* |
| WP_000066951.1 | 0.78534 | 0.08612 | *flhA* |
| WP_001295647.1 | 0.70971 | 0.09661 | *flhD* |
| WP_000122416.1 | 0.8652 | 0.47245 | *uspC* |
| WP_001295646.1 | 1.44509 | 0.11747 | *otsA* |
| WP_001295645.1 | 1.05484 | 0.82547 | *A1UI_02107* |
| WP_000100205.1 | 1.00875 | 0.99247 | *EKN05_006250* |
| WP_001187819.1 | 0.97484 | 0.93371 | *DXE50_13950* |
| WP_000548675.1 | 0.86835 | 0.38582 | *DXE50_13945* |
| WP_000082127.1 | 0.91158 | 0.58906 | *DXE50_13935* |
| WP_000917208.1 | 1.33276 | 0.05833 | *ftnA* |
| WP_000797560.1 | 0.80035 | 0.276 | *tyrP* |
| WP_000847902.1 | 1.09312 | 0.70928 | *yecA* |
| WP_001160187.1 | 1.1594 | 0.62881 | *pgsA* |

| | | | |
|---|---|---|---|
| WP_001283421.1 | 1.04552 | 0.85511 | *uvrC* |
| WP_000611335.1 | 0.89422 | 0.51549 | *uvrY* |
| WP_000106474.1 | 1.1275 | 0.61345 | *EKN05_006160* |
| WP_001272991.1 | 1.06012 | 0.85863 | *yecC* |
| WP_001158220.1 | 1.21293 | 0.52392 | *tcyL* |
| WP_001128215.1 | 1.06113 | 0.76508 | *dcyD* |
| WP_001296168.1 | 1.12844 | 0.35235 | *HMPREF1604_01381* |
| WP_000079739.1 | 0.92585 | 0.6921 | *fliC* |
| WP_001350519.1 | 0.98874 | 0.89906 | *KXD89_08915* |
| WP_000118897.1 | 0.96698 | 0.83961 | *HMPREF1595_04608* |
| WP_000790504.1 | 0.7604 | 0.11446 | *yedF* |
| WP_000922686.1 | 1.34887 | 0.31753 | *KXD89_08780* |
| WP_001157239.1 | 0.83217 | 0.2882 | *G711_02468* |
| WP_000218212.1 | 1.19218 | 0.21201 | *hchA* |
| WP_000920120.1 | 1.01511 | 0.97592 | *HMPREF1611_00004* |
| WP_001007805.1 | 0.69713 | 0.0807 | *HMPREF1589_00386* |
| WP_001302302.1 | 0.90087 | 0.54327 | *mtfA* |
| WP_001060244.1 | 1.05235 | 0.80156 | *amn* |
| WP_000532923.1 | 0.81574 | 0.14591 | *yeeN* |
| WP_001011008.1 | 0.83732 | 0.38191 | *HMPREF1602_03546* |
| AIN32411.1 | 1.55139 | 0.13979 | - |
| WP_001166160.1 | 1.14662 | 0.65589 | *cobT* |
| WP_001326708.1 | 0.92177 | 0.67667 | *cobS* |
| WP_000450409.1 | 0.99509 | 0.83975 | *yeeX* |
| WP_001105415.1 | 1.26595 | 0.31731 | *sbmC* |
| WP_000980589.1 | 0.87081 | 0.42242 | *sbcB* |
| WP_000234896.1 | 0.87337 | 0.43224 | *FSF13_012590* |
| WP_000019197.1 | 0.83458 | 0.37345 | *plaP* |
| WP_000754737.1 | 1.07517 | 0.70841 | *yeeZ* |
| WP_000131782.1 | 0.91784 | 0.52196 | *hisG* |
| WP_000009594.1 | 0.90289 | 0.45104 | *hisD* |
| WP_000108941.1 | 0.94242 | 0.64697 | *hisC* |
| WP_000080105.1 | 0.93205 | 0.59312 | *hisB* |
| WP_001103560.1 | 1.01621 | 0.95128 | *hisH* |
| WP_000586462.1 | 0.89689 | 0.5266 | *hisA* |
| WP_000880182.1 | 1.07384 | 0.71371 | *hisF* |
| WP_000954911.1 | 0.92288 | 0.6386 | *hisI* |
| WP_001393575.1 | 0.87121 | 0.40144 | *G736_02383* |
| WP_000043484.1 | 0.89891 | 0.56794 | *G736_02385* |
| WP_000520320.1 | 0.99207 | 0.91746 | *KXD89_08400* |
| WP_000601187.1 | 0.96935 | 0.85054 | *G736_02388* |
| WP_001407542.1 | 0.90456 | 0.45873 | *G736_02389* |
| WP_000272486.1 | 0.80093 | 0.11397 | *G736_02391* |
| WP_001100981.1 | 0.79734 | 0.19127 | *G736_02393* |
| WP_000783975.1 | 0.99324 | 0.9239 | *G736_02394* |
| WP_001023610.1 | 0.90633 | 0.56643 | *G736_02395* |
| WP_000699460.1 | 0.85513 | 0.31803 | *rfbB* |
| WP_000183060.1 | 0.85945 | 0.33937 | *galF* |

| | | | |
|---|---|---|---|
| WP_000454701.1 | 0.97529 | 0.88206 | *EKN05_005265* |
| WP_001252331.1 | 1.09019 | 0.71784 | *ECMP0215528_2416* |
| WP_001234768.1 | 1.14193 | 0.47256 | *dcd* |
| WP_001295424.1 | 0.96962 | 0.79361 | *udk* |
| WP_000288420.1 | 1.20201 | 0.43624 | *alkA* |
| WP_000469708.1 | 0.92492 | 0.55701 | *yegD* |
| WP_000678989.1 | 0.99669 | 0.96408 | *mdtA* |
| WP_000137877.1 | 1.1626 | 0.52469 | *baeR* |
| WP_000476011.1 | 0.94256 | 0.6477 | *ECMP0215528_2437* |
| WP_000844219.1 | 0.76809 | 0.06154 | *A1UI_02300* |
| WP_000823270.1 | 1.09506 | 0.44951 | *A1UI_02302* |
| WP_000853883.1 | 0.86227 | 0.35376 | *gatZ* |
| WP_001307281.1 | 0.98345 | 0.89334 | *gatY* |
| WP_000129551.1 | 1.1765 | 0.24143 | *fbaB* |
| WP_000846217.1 | 1.40244 | 0.15058 | *ECMP0215528_2449* |
| WP_000434038.1 | 0.99119 | 0.94309 | *DXE50_12360* |
| WP_001005448.1 | 0.93436 | 0.81031 | *apbC* |
| WP_001350533.1 | 0.89745 | 0.55854 | *metG* |
| WP_000950409.1 | 1.01306 | 0.96185 | *DXE50_12250* |
| WP_000598641.1 | 1.11994 | 0.6338 | *yehT* |
| WP_001295431.1 | 1.36197 | 0.30183 | *btsS* |
| WP_000569315.1 | 1.05652 | 0.86775 | *yehX* |
| WP_001130308.1 | 1.13327 | 0.49981 | *osmF* |
| WP_000871504.1 | 1.03223 | 0.88789 | *bglX* |
| WP_000097403.1 | 0.98824 | 0.87114 | *dld* |
| WP_000079538.1 | 1.66843 | 0.0846 | *DXE50_11940* |
| WP_001264861.1 | 1.22307 | 0.5056 | *dusC* |
| WP_000553555.1 | 0.8985 | 0.59008 | *cdd* |
| WP_001136389.1 | 1.22945 | 0.49433 | *A1UI_02361* |
| WP_001275118.1 | 0.72315 | 0.06204 | *mglC* |
| WP_000255039.1 | 0.93751 | 0.83277 | *A1UI_02364* |
| WP_001036964.1 | 0.9365 | 0.82552 | *mglB* |
| WP_000628648.1 | 0.89936 | 0.59322 | *galS* |
| WP_001139613.1 | 0.96036 | 0.99599 | *folE* |
| WP_000425438.1 | 1.14694 | 0.56315 | *O3K_08675* |
| WP_000489247.1 | 0.76181 | 0.05408 | *O3K_08670* |
| WP_000253273.1 | 1.14713 | 0.6548 | *lysP* |
| WP_000548294.1 | 0.80366 | 0.20707 | *DXE50_11840* |
| WP_000182053.1 | 1.00928 | 0.99159 | *DXE50_11835* |
| WP_000873894.1 | 1.05498 | 0.79053 | *nfo* |
| WP_001292460.1 | 1.21944 | 0.5121 | *psuG* |
| WP_000854447.1 | 1.12092 | 0.54042 | *fruA* |
| WP_000091263.1 | 0.9087 | 0.47807 | *fruK* |
| WP_000487246.1 | 0.87733 | 0.34117 | *HMPREF1589_01474* |
| WP_001136827.1 | 0.81447 | 0.15691 | *yeiP* |
| WP_001091940.1 | 0.87168 | 0.49462 | *KXD89_07695* |
| WP_000198828.1 | 1.06674 | 0.78835 | *yeiR* |
| WP_000470560.1 | 1.10437 | 0.75081 | *A1UI_02392* |

| | | | |
|---|---|---|---|
| WP_000637046.1 | 0.99503 | 0.95775 | *KXD89_07670* |
| WP_000194927.1 | 1.14992 | 0.64883 | *A1UI_02396* |
| WP_001234850.1 | 1.08174 | 0.6827 | *rsuA* |
| WP_000494183.1 | 0.9618 | 0.99581 | *rplY* |
| WP_000050793.1 | 0.96183 | 0.99564 | *yejK* |
| WP_001135667.1 | 1.16975 | 0.25507 | *yejL* |
| WP_000256203.1 | 0.9214 | 0.67526 | *yejM* |
| WP_001113639.1 | 0.9767 | 0.88458 | *narP* |
| WP_001026418.1 | 0.983 | 0.91169 | *ccmE* |
| WP_000778061.1 | 1.33821 | 0.21601 | *napA* |
| WP_000849209.1 | 0.87495 | 0.33174 | *eco* |
| WP_000758077.1 | 0.75345 | 0.16857 | *mqo* |
| WP_000865568.1 | 0.9691 | 0.96088 | *ompC* |
| WP_001249081.1 | 1.02776 | 0.90751 | *rcsD* |
| WP_001061917.1 | 1.08843 | 0.47086 | *rcsB* |
| WP_000876011.1 | 1.31518 | 0.36109 | *rcsC* |
| WP_000125282.1 | 1.6712 | 0.08359 | *A1UI_02435* |
| WP_000012305.1 | 0.94103 | 0.75002 | *ECMP0215528_2584* |
| WP_001281242.1 | 0.94827 | 0.90967 | *gyrA* |
| WP_000990765.1 | 0.92177 | 0.54128 | *ubiG* |
| WP_001075164.1 | 0.92497 | 0.74408 | *ECMP0215528_2589* |
| WP_000332037.1 | 0.96996 | 0.9568 | *DXE50_11490* |
| WP_000779105.1 | 0.91747 | 0.69199 | *glpQ* |
| WP_000948731.1 | 0.82277 | 0.18394 | *glpT* |
| WP_000857257.1 | 1.01912 | 0.94584 | *glpA* |
| WP_001209927.1 | 0.95475 | 0.71268 | *glpB* |
| WP_001000379.1 | 0.91998 | 0.62576 | *A1UI_02458* |
| WP_000921621.1 | 1.07118 | 0.77473 | *A1UI_02465* |
| WP_001300392.1 | 1.26006 | 0.32704 | *A1UI_02466* |
| WP_000577625.1 | 1.51303 | 0.16417 | *menE* |
| WP_001255628.1 | 1.0908 | 0.71603 | *menC* |
| WP_000639996.1 | 1.1298 | 0.34876 | *menB* |
| WP_001295284.1 | 0.96612 | 0.84668 | *menD* |
| WP_001191419.1 | 0.74144 | 0.08488 | *menF* |
| WP_000070621.1 | 1.03499 | 0.66738 | *elaB* |
| WP_000574091.1 | 1.39616 | 0.1561 | *EKN05_004155* |
| WP_001300687.1 | 1.11197 | 0.65568 | *rbn* |
| WP_000156701.1 | 0.87505 | 0.43869 | *nuoN* |
| WP_001056643.1 | 0.96229 | 0.8179 | *nuoL* |
| WP_000612644.1 | 1.01975 | 0.96354 | *nuoK* |
| WP_000172749.1 | 0.93321 | 0.59906 | *nuoI* |
| WP_000118507.1 | 0.83169 | 0.2867 | *nuoH* |
| WP_000190939.1 | 0.901 | 0.58148 | *A1UI_02496* |
| WP_000789500.1 | 0.94706 | 0.90098 | *WG3_01307* |
| WP_000545042.1 | 0.85706 | 0.26563 | *nuoE* |
| AIN32696.1 | 0.95838 | 0.9819 | - |
| WP_000386733.1 | 0.95503 | 0.71418 | *nuoB* |
| WP_000062997.1 | 0.86386 | 0.39638 | *nuoA* |

| | | | |
|---|---|---|---|
| WP_000622280.1 | 0.75267 | 0.16705 | *lrhA* |
| WP_000074527.1 | 0.92248 | 0.54482 | *DXE50_11220* |
| WP_000813859.1 | 1.05261 | 0.83252 | *yfbR* |
| WP_001203392.1 | 1.18082 | 0.36296 | *A1UI_02507* |
| WP_000426124.1 | 0.95567 | 0.96257 | *yfbU* |
| WP_000095707.1 | 1.07318 | 0.52252 | *ackA* |
| WP_000086722.1 | 1.04171 | 0.64038 | *EKN05_004025* |
| WP_000437935.1 | 0.98594 | 0.92729 | *yfcD* |
| WP_000772452.1 | 1.27448 | 0.30362 | *DXE50_11170* |
| WP_000042442.1 | 1.23341 | 0.37414 | *O3K_07990* |
| WP_000068457.1 | 0.79027 | 0.09432 | *folX* |
| WP_001028341.1 | 1.31272 | 0.24809 | *A1UI_02518* |
| WP_000737621.1 | 0.87205 | 0.32046 | *hisJ* |
| WP_000334220.1 | 1.13007 | 0.60664 | *purF* |
| WP_000146992.1 | 0.8793 | 0.52115 | *dedD* |
| WP_000584546.1 | 0.89839 | 0.43061 | *A1UI_02530* |
| WP_000118404.1 | 0.93667 | 0.82679 | *accD* |
| WP_001283586.1 | 0.82385 | 0.34131 | *truA* |
| WP_001289167.1 | 0.89327 | 0.4079 | *A1UI_02534* |
| WP_000699148.1 | 1.01659 | 0.74451 | *pdxB* |
| WP_000615834.1 | 1.28516 | 0.40396 | *A1UI_02536* |
| WP_000817178.1 | 0.83789 | 0.24084 | *fabB* |
| WP_000683799.1 | 1.03896 | 0.85861 | *mnmC* |
| WP_000559764.1 | 0.87073 | 0.42213 | *DXE50_11045* |
| WP_001043825.1 | 1.00321 | 0.9888 | *mepA* |
| WP_001333535.1 | 1.0499 | 0.81187 | *aroC* |
| WP_001300582.1 | 1.06643 | 0.74345 | *prmB* |
| WP_000730806.1 | 1.12641 | 0.70027 | *smrB* |
| WP_001195819.1 | 0.74198 | 0.14711 | *sixA* |
| WP_000426176.1 | 1.09485 | 0.70426 | *fadJ* |
| WP_000531952.1 | 1.13177 | 0.60217 | *fadI* |
| WP_001296261.1 | 1.26369 | 0.19448 | *WG3_01240* |
| WP_001295701.1 | 1.11543 | 0.38817 | *AC28_2640* |
| WP_203277081.1 | 1.03011 | 0.89716 | - |
| WP_000958671.1 | 0.93925 | 0.74321 | *G925_02382* |
| WP_000703651.1 | 1.03725 | 0.8818 | *KY274_06780* |
| WP_001030215.1 | 0.99925 | 0.98848 | *KXD89_06785* |
| WP_000556048.1 | 0.84898 | 0.41876 | |
| WP_000785931.1 | 1.03691 | 0.88291 | *DXE50_10435* |
| WP_001295458.1 | 1.0018 | 0.98347 | *DXE50_10425* |
| WP_000955906.1 | 0.74474 | 0.15209 | *A1UI_02584* |
| WP_000170346.1 | 1.26782 | 0.10869 | *glk* |
| WP_000376337.1 | 0.95579 | 0.7183 | *nupC* |
| AIN32800.1 | 0.82082 | 0.33247 | - |
| WP_000695655.1 | 0.94497 | 0.88604 | *gltX* |
| WP_000443661.1 | 1.12008 | 0.54326 | *ligA* |
| WP_001300494.1 | 0.9852 | 0.87953 | *zipA* |
| WP_000254839.1 | 0.87679 | 0.51237 | *cysZ* |

| | | | |
|---|---|---|---|
| WP_000034402.1 | 1.18953 | 0.21676 | *cysK* |
| WP_000487600.1 | 0.93828 | 0.83824 | *HMPREF0454_04299* |
| WP_000623140.1 | 0.88673 | 0.49155 | *ptsI* |
| WP_000522247.1 | 1.092 | 0.45926 | *DXE50_10250* |
| WP_000096674.1 | 0.78696 | 0.16702 | *pdxK* |
| WP_001336044.1 | 1.05944 | 0.811 | *cysM* |
| WP_000290230.1 | 0.92427 | 0.55375 | *G711_02922* |
| WP_000517431.1 | 0.90302 | 0.45162 | *ucpA* |
| WP_000966470.1 | 0.83993 | 0.39001 | *murR* |
| WP_001159160.1 | 1.02985 | 0.90599 | *murQ* |
| WP_001327042.1 | 1.14866 | 0.65151 | *pbp4b* |
| WP_001350538.1 | 0.98689 | 0.88883 | *yfeX* |
| WP_000838944.1 | 0.9407 | 0.71873 | *EKN05_003410* |
| WP_000405996.1 | 1.12742 | 0.69801 | *ypeA* |
| WP_000102886.1 | 1.42065 | 0.23945 | *amiA* |
| WP_000801365.1 | 1.06614 | 0.84349 | *hemF* |
| WP_001111023.1 | 1.16649 | 0.61414 | *A1UI_02631* |
| WP_000372316.1 | 1.0459 | 0.89493 | *eutC* |
| WP_000769961.1 | 1.05207 | 0.87908 | *eutB* |
| WP_010723125.1 | 1.19146 | 0.45876 | *yffS* |
| WP_000387713.1 | 0.79896 | 0.27235 | *eutM* |
| WP_000342644.1 | 1.04063 | 0.64467 | *maeB* |
| WP_000258236.1 | 1.55813 | 0.05912 | *KXD89_06205* |
| WP_001277801.1 | 1.42778 | 0.13003 | *dapE* |
| WP_000679799.1 | 1.17167 | 0.60356 | *ypfH* |
| WP_001267498.1 | 0.98767 | 0.93526 | *DXE50_09995* |
| WP_001295467.1 | 0.95145 | 0.9324 | *purC* |
| WP_001297320.1 | 1.00867 | 0.77897 | *bamC* |
| WP_001311023.1 | 0.9517 | 0.93415 | *dapA* |
| WP_000176187.1 | 0.9853 | 0.88006 | *EKN05_003185* |
| WP_001068682.1 | 0.81109 | 0.14676 | *EKN05_003180* |
| WP_000489667.1 | 1.28526 | 0.28702 | *bepA* |
| WP_000166446.1 | 1.00122 | 0.96785 | *arsC* |
| WP_001307333.1 | 0.90707 | 0.62162 | *hda* |
| WP_000198328.1 | 0.7067 | 0.09262 | *uraA* |
| WP_001295473.1 | 0.77533 | 0.06619 | *upp* |
| WP_001336050.1 | 1.00042 | 0.96347 | *purM* |
| WP_001028612.1 | 0.85519 | 0.439 | *purN* |
| WP_000529576.1 | 1.12344 | 0.53198 | *ppk1* |
| WP_001121363.1 | 0.97549 | 0.82592 | *O3K_06910* |
| WP_000076001.1 | 1.23994 | 0.47621 | *O3K_06895* |
| WP_000138270.1 | 0.81645 | 0.16311 | *guaA* |
| WP_001299507.1 | 0.88395 | 0.47478 | *guaB* |
| WP_000937912.1 | 0.90682 | 0.56856 | *xseA* |
| WP_001301158.1 | 1.02227 | 0.95683 | *KXD89_06000* |
| WP_000249410.1 | 1.06411 | 0.75287 | *engA* |
| WP_001177052.1 | 1.09114 | 0.46203 | *bamB* |
| WP_000409205.1 | 1.0171 | 0.95489 | *DXE50_09780* |

| | | | |
|---|---|---|---|
| WP_001107167.1 | 0.95132 | 0.93145 | *hisS* |
| WP_000551807.1 | 0.88355 | 0.47235 | *ispG* |
| WP_001090866.1 | 1.13607 | 0.49087 | *rodZ* |
| WP_000003317.1 | 0.81154 | 0.13628 | *trmG/rlmN* |
| WP_000963837.1 | 1.10469 | 0.41972 | *ndk* |
| WP_000736312.1 | 0.98936 | 0.90248 | *yfhM* |
| WP_000108626.1 | 1.23555 | 0.24251 | *sseA* |
| WP_001295479.1 | 0.99071 | 0.9493 | *sseB* |
| WP_000523616.1 | 1.13995 | 0.58089 | *iscX* |
| WP_001124469.1 | 1.43 | 0.12835 | *fdx* |
| WP_001196613.1 | 1.35691 | 0.08833 | *hscA* |
| WP_000384413.1 | 0.78642 | 0.16581 | *hscB* |
| WP_000028953.1 | 0.94454 | 0.7362 | *iscA* |
| WP_000331707.1 | 0.79767 | 0.11094 | *iscU* |
| WP_001295373.1 | 0.94137 | 0.86032 | *iscS* |
| WP_001241357.1 | 1.22067 | 0.27156 | *iscR* |
| WP_000940019.1 | 0.97095 | 0.80094 | *trmJ* |
| WP_000553451.1 | 1.07898 | 0.50247 | *suhB* |
| WP_000982994.1 | 0.83756 | 0.38265 | *csiE* |
| WP_000919159.1 | 0.97189 | 0.94767 | *glyA* |
| WP_000717694.1 | 0.83192 | 0.18729 | *glnB* |
| WP_001295369.1 | 1.1088 | 0.66452 | *A1UI_02746* |
| WP_001215861.1 | 1.43173 | 0.22904 | *qseG* |
| WP_000970102.1 | 1.03612 | 0.87093 | *purL* |
| WP_001013779.1 | 0.95278 | 0.79524 | *DXE50_09565* |
| WP_001297412.1 | 1.0606 | 0.76724 | *pdxJ* |
| WP_000020749.1 | 0.85215 | 0.24899 | *era* |
| WP_001068343.1 | 0.98012 | 0.85145 | *rnc* |
| WP_000002541.1 | 1.03195 | 0.89912 | *lepB* |
| WP_000790168.1 | 1.00091 | 0.8135 | *lepA* |
| WP_000812053.1 | 1.05425 | 0.82733 | *rseB* |
| WP_001168459.1 | 0.93867 | 0.70947 | *rseA* |
| WP_001295364.1 | 1.52798 | 0.15424 | *rpoE* |
| WP_001094491.1 | 1.04256 | 0.86461 | *nadB* |
| WP_001295363.1 | 0.77599 | 0.21627 | *trmN* |
| WP_000219203.1 | 1.04573 | 0.82956 | *srmB* |
| WP_000627807.1 | 1.12902 | 0.35081 | *grcA* |
| WP_001262716.1 | 1.12128 | 0.63017 | *ung* |
| WP_000997403.1 | 0.90984 | 0.64005 | *DXE50_09455* |
| WP_001098726.1 | 0.9575 | 0.79576 | *trxC* |
| WP_001300438.1 | 0.70805 | 0.0944 | *A1UI_02775* |
| WP_000083005.1 | 1.14827 | 0.5598 | *pat* |
| WP_000949265.1 | 1.08856 | 0.65651 | *pssA* |
| WP_000841103.1 | 1.16769 | 0.5126 | *kgtP* |
| WP_001235102.1 | 1.03119 | 0.68293 | *clpB* |
| WP_000040169.1 | 1.19316 | 0.45507 | *pgeF* |
| WP_000079100.1 | 1.1375 | 0.48635 | *rluD* |
| WP_000197686.1 | 0.96121 | 0.99864 | *bamD* |

| | | | |
|---|---|---|---|
| WP_000200120.1 | 0.96535 | 0.77023 | *A1UI_02786* |
| WP_000225229.1 | 1.01243 | 0.96396 | *tyrA* |
| WP_001168037.1 | 0.92162 | 0.6761 | *aroF* |
| pdb\|6H58\|PP | 0.82254 | 0.18316 | - |
| WP_000264777.1 | 1.1373 | 0.58773 | *trmD* |
| WP_000043335.1 | 0.88733 | 0.3823 | *rimM* |
| WP_000256450.1 | 0.98102 | 0.90466 | *rpsP* |
| WP_000460035.1 | 0.89165 | 0.52183 | *ffh* |
| WP_001393454.1 | 1.04405 | 0.6311 | *grpE* |
| WP_001059169.1 | 1.07177 | 0.82946 | *nadK* |
| WP_001117838.1 | 0.89785 | 0.5877 | *DXE50_09280* |
| WP_001300750.1 | 1.06657 | 0.84242 | *ratA* |
| WP_000162574.1 | 1.06712 | 0.78717 | *smpB* |
| WP_000101723.1 | 0.93909 | 0.7426 | *KXD89_05435* |
| WP_000155570.1 | 0.97919 | 0.89708 | *rnlA* |
| WP_000993126.1 | 1.16669 | 0.40038 | *csiD* |
| WP_000772831.1 | 1.30591 | 0.07583 | *gabD* |
| WP_001087611.1 | 1.30406 | 0.07719 | *gabT* |
| WP_001229442.1 | 1.13745 | 0.67578 | *A1UI_02878* |
| WP_000115383.1 | 1.03327 | 0.88333 | *stpA* |
| WP_001295174.1 | 1.14049 | 0.57952 | *DXE50_18090* |
| WP_000985494.1 | 0.97507 | 0.93261 | *proV* |
| WP_000774988.1 | 0.99532 | 0.93538 | *proW* |
| WP_001216525.1 | 0.90089 | 0.58076 | *DXE50_18140* |
| WP_000378442.1 | 0.9925 | 0.91983 | *emrR* |
| WP_001326681.1 | 0.94004 | 0.71571 | *emrA* |
| WP_001130211.1 | 1.32891 | 0.06059 | *luxS* |
| WP_000611804.1 | 1.03727 | 0.86594 | *gshA* |
| WP_000273290.1 | 1.08776 | 0.79032 | *A1UI_02899* |
| WP_000906486.1 | 1.13704 | 0.48782 | *csrA* |
| WP_000047184.1 | 0.95644 | 0.96805 | *alaS* |
| WP_000963143.1 | 1.2193 | 0.16814 | *recA* |
| WP_000132231.1 | 0.99248 | 0.9197 | *pncC* |
| WP_001295177.1 | 1.11995 | 0.63377 | *mltB* |
| WP_000148878.1 | 0.75692 | 0.10856 | *srlE* |
| WP_000216194.1 | 0.76981 | 0.06372 | *A1UI_02912* |
| WP_001077358.1 | 0.79681 | 0.10605 | *srlD* |
| WP_000804550.1 | 0.88323 | 0.47084 | *srlR* |
| WP_001287420.1 | 1.09011 | 0.65065 | *WG3_03337* |
| WP_001394680.1 | 1.34031 | 0.32815 | *ascG* |
| WP_000337665.1 | 1.17646 | 0.59389 | *hypB* |
| WP_001212985.1 | 1.44872 | 0.21384 | *hypD* |
| WP_001272928.1 | 0.96503 | 0.76848 | *mutS* |
| WP_001272592.1 | 1.39434 | 0.06291 | *nlpD* |
| WP_000254708.1 | 1.2972 | 0.3863 | *pcm* |
| WP_001295182.1 | 0.95063 | 0.76412 | *surE* |
| WP_000568943.1 | 0.92631 | 0.65384 | *truD* |
| WP_001219242.1 | 0.91311 | 0.49903 | *ispF* |

| | | | |
|---|---|---|---|
| WP_000246138.1 | 0.92314 | 0.68185 | *ispD* |
| WP_000517476.1 | 0.9756 | 0.88324 | *ftsB* |
| WP_001173673.1 | 1.19632 | 0.555 | *cysC* |
| WP_001090361.1 | 1.04254 | 0.8432 | *cysN* |
| WP_000372108.1 | 0.92547 | 0.65009 | *cysD* |
| WP_000039850.1 | 1.04551 | 0.85515 | *cysH* |
| WP_001290679.1 | 1.0656 | 0.7468 | *cysI* |
| WP_000211913.1 | 1.07105 | 0.7248 | *cysJ* |
| WP_000108294.1 | 0.95664 | 0.7918 | *queD* |
| WP_000036723.1 | 0.85492 | 0.31701 | *eno* |
| WP_000210878.1 | 0.89381 | 0.53541 | *pyrG* |
| WP_001071648.1 | 1.34612 | 0.32091 | *mazG* |
| WP_000254738.1 | 0.76739 | 0.1972 | *A1UI_02996* |
| WP_000226815.1 | 1.02108 | 0.93501 | *DXE50_18695* |
| WP_000046812.1 | 1.05836 | 0.86307 | *rumA* |
| WP_000186450.1 | 1.11055 | 0.73642 | *barA* |
| WP_000098255.1 | 1.28276 | 0.29079 | *gudD* |
| WP_000807753.1 | 1.07399 | 0.7131 | *EKN05_002460* |
| WP_000206990.1 | 1.00779 | 0.9796 | *ECMP0215528_3251* |
| WP_000342431.1 | 1.3304 | 0.34082 | *syd* |
| WP_000100421.1 | 0.90839 | 0.5753 | *queF* |
| WP_000627995.1 | 1.12159 | 0.37088 | *ppnN* |
| WP_000450476.1 | 0.88077 | 0.35507 | *DXE50_18755* |
| WP_000013588.1 | 1.18063 | 0.48275 | *fucO* |
| WP_000724153.1 | 1.05581 | 0.82242 | *fucI* |
| WP_000920840.1 | 1.24973 | 0.34471 | *fucU* |
| WP_000642344.1 | 0.95771 | 0.81426 | *fucR* |
| WP_001045520.1 | 0.98232 | 0.91059 | *rlmM* |
| WP_000044401.1 | 0.99659 | 0.9637 | *gcvA* |
| WP_000750398.1 | 0.83743 | 0.30489 | *DXE50_18820* |
| WP_001300698.1 | 0.97257 | 0.87157 | *O3K_05445* |
| WP_000184261.1 | 0.96752 | 0.84209 | *A1UI_03026* |
| WP_000117728.1 | 0.89311 | 0.5109 | *O3K_05435* |
| WP_000678646.1 | 0.95395 | 0.77938 | *mltA* |
| WP_001138201.1 | 0.94143 | 0.72205 | *ptrA* |
| WP_000946938.1 | 0.94018 | 0.7468 | *recC* |
| WP_000816232.1 | 1.26197 | 0.32386 | *thyA* |
| WP_000204658.1 | 0.96918 | 0.84976 | *lgt* |
| WP_000957910.1 | 0.92602 | 0.56252 | *EL76_1009* |
| WP_000564489.1 | 0.95422 | 0.8008 | *rppH* |
| WP_001199295.1 | 1.15793 | 0.42496 | *EKN05_001155* |
| WP_000899054.1 | 0.72085 | 0.05953 | *aas* |
| WP_000201063.1 | 0.99256 | 0.94834 | *A311_03510* |
| WP_001120711.1 | 0.71058 | 0.09779 | *lysA* |
| WP_000848659.1 | 1.03134 | 0.89179 | *ygeA* |
| WP_000256438.1 | 1.08056 | 0.49708 | *DXE50_18990* |
| WP_000603502.1 | 1.12956 | 0.608 | *HMPREF1620_04171* |
| WP_000383237.1 | 1.2067 | 0.53542 | *kduI* |

| | | | |
|---|---|---|---|
| WP_000656030.1 | 0.79923 | 0.1959 | *KXD89_04355* |
| WP_000065950.1 | 0.8269 | 0.272 | *DXE50_19010* |
| WP_001301085.1 | 0.69628 | 0.0797 | *ECMP0215528_3346* |
| WP_000417791.1 | 0.84713 | 0.41282 | *A1UI_03071* |
| WP_000502395.1 | 1.15547 | 0.63706 | *ygfK* |
| WP_001192820.1 | 1.30868 | 0.37003 | *idi* |
| WP_000003071.1 | 0.96519 | 0.97954 | *lysS* |
| WP_010723217.1 | 0.9896 | 0.86485 | *prfB* |
| WP_000813200.1 | 0.96019 | 0.80819 | *recJ* |
| WP_000715214.1 | 0.84558 | 0.22774 | *dsbC* |
| WP_000806638.1 | 1.11667 | 0.72233 | *xerD* |
| WP_001055874.1 | 0.83566 | 0.19787 | *fldB* |
| WP_000354046.1 | 1.33009 | 0.22583 | *sdhE* |
| WP_000886062.1 | 1.07418 | 0.51902 | *ygfZ* |
| QRG61511.1 | 1.18797 | 0.34505 | - |
| WP_000195062.1 | 0.79551 | 0.10584 | *gcvP* |
| WP_000068701.1 | 0.77888 | 0.07218 | *gcvT* |
| WP_001192229.1 | 1.114 | 0.65007 | *ubiI* |
| WP_000111195.1 | 1.60437 | 0.11159 | *ubiH* |
| WP_001290136.1 | 1.18123 | 0.3619 | *pepP* |
| WP_001295378.1 | 0.86812 | 0.41225 | *ygfB* |
| WP_001276008.1 | 1.02403 | 0.92523 | *zapA* |
| WP_001151604.1 | 0.88417 | 0.47606 | *DXE50_19325* |
| WP_000189743.1 | 0.94454 | 0.88297 | *rpiA* |
| WP_000828351.1 | 0.99858 | 0.9854 | *argP* |
| WP_000669834.1 | 1.07622 | 0.51194 | *DXE50_19370* |
| WP_000389818.1 | 1.0984 | 0.43901 | *mscS* |
| WP_000034372.1 | 0.92259 | 0.72742 | *fbaA* |
| WP_000111269.1 | 0.91938 | 0.70521 | *pgk* |
| WP_000218480.1 | 1.03837 | 0.86116 | *epd* |
| WP_000098614.1 | 0.9174 | 0.69154 | *ECMP0215528_3427* |
| WP_000105566.1 | 0.92297 | 0.73014 | *speB* |
| WP_001300904.1 | 0.98091 | 0.90517 | *speA* |
| WP_001062128.1 | 1.02225 | 0.72031 | *metK* |
| WP_001222509.1 | 0.97499 | 0.82316 | *rsmE* |
| WP_000593273.1 | 1.0052 | 0.79434 | *gshB* |
| WP_001053178.1 | 0.98693 | 0.88908 | *yqgE* |
| WP_000017106.1 | 1.02042 | 0.96174 | *yqgF* |
| WP_000997795.1 | 0.85003 | 0.24199 | *yggS* |
| WP_001094831.1 | 0.9601 | 0.74165 | *yggT* |
| WP_001174777.1 | 0.91436 | 0.50507 | *A1UI_03160* |
| WP_000239943.1 | 1.10151 | 0.75753 | *A1UI_03161* |
| WP_000984796.1 | 1.06997 | 0.72915 | *DXE50_19655* |
| WP_001107564.1 | 0.88735 | 0.49533 | *DXE50_19660* |
| WP_000786911.1 | 1.07689 | 0.50962 | *trmB* |
| WP_000091700.1 | 0.94561 | 0.89062 | *yggX* |
| WP_000760323.1 | 1.29862 | 0.38426 | *mltC* |
| WP_001049791.1 | 1.0832 | 0.80137 | *nupG* |

| | | | |
|---|---|---|---|
| WP_001326492.1 | 0.85629 | 0.36895 | *A1UI_03171* |
| WP_000084131.1 | 1.00433 | 0.79821 | *glcB* |
| WP_000853256.1 | 1.01126 | 0.9811 | *DXE50_19835* |
| WP_001297764.1 | 0.87408 | 0.50292 | *ECMP0215528_3474* |
| WP_001297309.1 | 1.14967 | 0.4491 | *EKN05_000500* |
| WP_001295515.1 | 1.40249 | 0.05834 | *ECMP0215528_3491* |
| WP_001221939.1 | 1.54416 | 0.14411 | *A1UI_03277* |
| WP_000083065.1 | 1.29023 | 0.27962 | *hybC* |
| WP_000262172.1 | 1.14796 | 0.45421 | *AB05_3609* |
| WP_000439331.1 | 1.14098 | 0.57826 | *yqhA* |
| WP_000018760.1 | 1.19011 | 0.3398 | *ECMP0215528_3505* |
| WP_001240712.1 | 0.75711 | 0.10887 | *exbD* |
| WP_000527844.1 | 0.76883 | 0.06248 | *DXE50_20010* |
| WP_001301079.1 | 1.54333 | 0.06482 | *G894_03028* |
| WP_000268419.1 | 1.4819 | 0.18673 | *DXE50_20020* |
| WP_001058802.1 | 0.95878 | 0.9847 | *yqhD* |
| WP_000095187.1 | 0.92179 | 0.63376 | *ygiQ* |
| WP_000059388.1 | 0.95324 | 0.77615 | *ftsP* |
| WP_000965722.1 | 0.92072 | 0.62902 | *A1UI_03298* |
| WP_001281881.1 | 0.96355 | 0.7604 | *parC* |
| WP_001295629.1 | 1.23444 | 0.24459 | *A1UI_03300* |
| WP_001221495.1 | 1.25505 | 0.33553 | *qseB* |
| WP_000673402.1 | 1.11814 | 0.71898 | *A1UI_03306* |
| WP_000065430.1 | 0.95276 | 0.70197 | *mdaB* |
| WP_000958598.1 | 1.1047 | 0.59692 | *DXE50_20115* |
| WP_000195296.1 | 0.94568 | 0.66419 | *parE* |
| WP_000444747.1 | 1.00549 | 0.98738 | *cpdA* |
| WP_000917117.1 | 0.93369 | 0.68693 | *DXE50_20140* |
| WP_000831543.1 | 1.00905 | 0.97536 | *ygiB* |
| WP_000442860.1 | 1.10978 | 0.57884 | *DXE50_20155* |
| WP_001076997.1 | 1.25156 | 0.21411 | *ribB* |
| WP_001295626.1 | 0.92386 | 0.55172 | *ubiK* |
| WP_000350095.1 | 1.15819 | 0.63134 | *glgS* |
| WP_000869178.1 | 1.03179 | 0.68047 | *hldE* |
| WP_001301081.1 | 1.0432 | 0.84037 | *glnE* |
| WP_000708487.1 | 1.0962 | 0.70034 | *cca* |
| WP_001295541.1 | 0.93275 | 0.68268 | *folB* |
| WP_001264352.1 | 0.88736 | 0.38245 | *tsaD* |
| pdb\|6H58\|uu | 0.79382 | 0.10198 | - |
| WP_000918827.1 | 0.82238 | 0.33699 | *dnaG* |
| WP_000437376.1 | 1.02152 | 0.72343 | *rpoD* |
| WP_000228937.1 | 1.13654 | 0.58971 | *mug* |
| WP_000018003.1 | 0.98247 | 0.91128 | *EKN05_000130* |
| WP_000094721.1 | 1.60313 | 0.11219 | *HMPREF1599_02928* |
| WP_001301395.1 | 1.18725 | 0.57253 | *ygjG* |
| WP_000450594.1 | 0.81404 | 0.31316 | *HMPREF1599_02930* |
| WP_000212475.1 | 1.17797 | 0.48879 | *HMPREF1599_02931* |
| WP_000695487.1 | 0.96216 | 0.83142 | *ygjK* |

| | | | |
|---|---|---|---|
| WP_000018695.1 | 0.97376 | 0.87098 | *rlmG* |
| WP_000617698.1 | 0.98839 | 0.9386 | *KXD89_03155* |
| WP_001098806.1 | 0.67892 | 0.06112 | *ECMP0215528_3592* |
| WP_000211655.1 | 1.04169 | 0.90583 | *sstT* |
| WP_001199390.1 | 1.60044 | 0.11349 | *HMPREF1599_02950* |
| WP_000187442.1 | 1.07203 | 0.7209 | *uxaC* |
| WP_000406488.1 | 0.91969 | 0.6245 | *exuR* |
| WP_001295543.1 | 1.06115 | 0.85604 | *DXE50_20445* |
| WP_000096091.1 | 1.54683 | 0.1425 | *HMPREF1599_02960* |
| WP_000531213.1 | 1.37888 | 0.28255 | *HMPREF1599_02962* |
| WP_001041009.1 | 1.14799 | 0.5605 | *DXE50_20485* |
| WP_000622126.1 | 0.79908 | 0.27267 | *A1UI_03392* |
| WP_000719990.1 | 0.96482 | 0.76734 | *DXE50_20515* |
| WP_000861734.1 | 0.76558 | 0.12365 | *O3K_03370* |
| WP_000107723.1 | 0.91052 | 0.63446 | *tdcC* |
| WP_001307405.1 | 1.33437 | 0.3357 | *O3K_03300* |
| WP_000072187.1 | 0.9107 | 0.63514 | *DXE50_20600* |
| WP_001336162.1 | 1.31749 | 0.35795 | *agaV* |
| WP_000809262.1 | 0.86695 | 0.30113 | *rsmI* |
| WP_000249104.1 | 0.92075 | 0.53625 | *lpoA* |
| WP_001158034.1 | 0.78199 | 0.15614 | *diaA* |
| WP_000646033.1 | 1.08734 | 0.66115 | *ECMP0215528_3660* |
| WP_000147574.1 | 1.36264 | 0.30105 | *A1UI_03435* |
| WP_001300423.1 | 1.14433 | 0.66082 | *A1UI_03436* |
| WP_000908554.1 | 1.0288 | 0.90293 | *DXE50_20720* |
| WP_001295552.1 | 0.84861 | 0.41756 | *ubiT* |
| WP_000130380.1 | 1.01555 | 0.95347 | *O3K_03135* |
| WP_000224351.1 | 0.96026 | 0.82407 | *DXE50_20745* |
| WP_001295553.1 | 0.9968 | 0.83199 | *deaD* |
| WP_000802080.1 | 0.9786 | 0.89338 | *nlpI* |
| WP_001295554.1 | 0.95436 | 0.95321 | *pnp* |
| WP_000089698.1 | 0.86335 | 0.39453 | *truB* |
| WP_001040205.1 | 1.00792 | 0.7823 | *rbfA* |
| WP_000133044.1 | 1.03721 | 0.65839 | *infB* |
| WP_001031057.1 | 0.95611 | 0.96571 | *nusA* |
| WP_001300397.1 | 0.90407 | 0.4565 | *rimP* |
| WP_000207680.1 | 1.33546 | 0.05679 | *argG* |
| WP_001295556.1 | 0.94819 | 0.67753 | *secG* |
| WP_000071134.1 | 0.92215 | 0.72436 | *glmM* |
| WP_000764731.1 | 1.12615 | 0.52297 | *folP* |
| WP_001107467.1 | 0.99334 | 0.84775 | *ftsH* |
| WP_000145975.1 | 1.01285 | 0.96256 | *rlmE* |
| WP_001054420.1 | 0.78554 | 0.16385 | *yhbY* |
| WP_001148001.1 | 0.98007 | 0.90911 | *greA* |
| WP_001212619.1 | 0.96331 | 0.8226 | *WG3_03920* |
| WP_000673575.1 | 0.90028 | 0.43914 | *cgtA* |
| pdb\|6H58\|WW | 0.77099 | 0.05942 | - |
| WP_000271401.1 | 0.9449 | 0.88549 | *rplU* |

| | | | |
|---|---|---|---|
| WP_001047336.1 | 0.80322 | 0.11855 | *ispB* |
| WP_000357259.1 | 0.9125 | 0.65801 | *murA* |
| WP_000429656.1 | 0.73331 | 0.07408 | *ibaG* |
| WP_000004488.1 | 1.21355 | 0.41257 | *mlaB* |
| WP_000476487.1 | 1.16128 | 0.27304 | *mlaC* |
| WP_001296448.1 | 1.23207 | 0.24908 | *mlaD* |
| WP_000925795.1 | 1.20725 | 0.42536 | *mlaE* |
| WP_000438245.1 | 1.53797 | 0.067 | *mlaF* |
| WP_001295557.1 | 0.98132 | 0.85808 | *kdsD* |
| WP_000030005.1 | 1.09739 | 0.62351 | *kdsC* |
| WP_000030537.1 | 1.02787 | 0.91252 | *lptC* |
| WP_000669785.1 | 0.99437 | 0.93014 | *lptA* |
| WP_000224099.1 | 1.10111 | 0.60989 | *lptB* |
| WP_000809057.1 | 1.13619 | 0.5906 | *rpoN* |
| WP_001176599.1 | 1.39991 | 0.15278 | *DXE50_20960* |
| WP_000183676.1 | 0.8531 | 0.25214 | *ptsN* |
| WP_000243741.1 | 1.11752 | 0.55199 | *rapZ* |
| WP_001300411.1 | 1.08893 | 0.46923 | *elbB* |
| WP_000809774.1 | 0.96455 | 0.76589 | *arcB* |
| WP_000054239.1 | 0.84213 | 0.21704 | *nanE* |
| WP_000108454.1 | 0.73214 | 0.0726 | *nanT* |
| WP_000523845.1 | 0.88501 | 0.47796 | *nanR* |
| WP_000366129.1 | 1.04911 | 0.84363 | *sspB* |
| WP_000257293.1 | 1.01306 | 0.75977 | *sspA* |
| WP_000829818.1 | 0.87681 | 0.43293 | *rpsI* |
| WP_000847559.1 | 0.89324 | 0.53181 | *rplM* |
| WP_001192332.1 | 0.88128 | 0.4631 | *zapE* |
| WP_001295270.1 | 0.94112 | 0.85852 | *DXE50_21085* |
| WP_001295271.1 | 1.01302 | 0.9732 | *degQ* |
| WP_000497723.1 | 1.18585 | 0.57526 | *DXE50_21095* |
| WP_001295272.1 | 1.07112 | 0.52979 | *mdh* |
| WP_001257846.1 | 0.89371 | 0.40985 | *argR* |
| WP_000055909.1 | 0.97701 | 0.83431 | *tldD* |
| WP_001253618.1 | 0.84257 | 0.39829 | *yhdP* |
| WP_000123197.1 | 1.09228 | 0.64248 | *DXE50_21150* |
| WP_000203105.1 | 1.13119 | 0.6037 | *A1UI_03526* |
| WP_000802511.1 | 0.95799 | 0.79803 | *WG3_03983* |
| WP_000913396.1 | 0.93526 | 0.8167 | *mreB* |
| WP_001241469.1 | 1.20309 | 0.54217 | *A1UI_03530* |
| WP_001148481.1 | 1.14902 | 0.30083 | *EL76_0511* |
| WP_000884639.1 | 0.95213 | 0.93722 | *accC* |
| WP_001175728.1 | 1.12438 | 0.70484 | *panF* |
| WP_001145827.1 | 1.09616 | 0.62805 | *prmA* |
| WP_000462905.1 | 1.05662 | 0.7837 | *fis* |
| WP_001286216.1 | 1.18425 | 0.2265 | *EKN05_022560* |
| WP_000451243.1 | 1.01544 | 0.95383 | *aroE* |
| WP_001301412.1 | 0.84471 | 0.32886 | *tsaC* |
| WP_001129722.1 | 0.9702 | 0.86245 | *A1UI_03556* |

| | | | |
|---|---|---|---|
| WP_000114984.1 | 0.97423 | 0.81895 | *def* |
| WP_000004473.1 | 1.01505 | 0.96406 | *fmt* |
| WP_000744778.1 | 0.93813 | 0.62453 | *rsmB* |
| WP_000691382.1 | 1.1033 | 0.68001 | *trkA* |
| WP_000022442.1 | 1.16824 | 0.39613 | *mscL* |
| WP_000285607.1 | 0.80975 | 0.30124 | *zntR* |
| WP_001216368.1 | 1.02364 | 0.71443 | *rplQ* |
| WP_001162094.1 | 0.99897 | 0.82219 | *rpoA* |
| pdb\|7OI0\|D | 0.97051 | 0.95421 | - |
| pdb\|6H58\|kk | 0.83955 | 0.24772 | - |
| pdb\|6H58\|mm | 0.92864 | 0.76983 | - |
| WP_001118861.1 | 0.90465 | 0.45915 | *secY* |
| pdb\|6H58\|LL | 0.85285 | 0.30711 | - |
| pdb\|6H58\|ZZ | 0.78675 | 0.08691 | - |
| pdb\|7OE0\|E | 0.97401 | 0.9376 | - |
| pdb\|6H58\|OO | 0.92819 | 0.7667 | - |
| pdb\|6H58\|GG | 0.93849 | 0.83974 | - |
| WP_000062611.1 | 0.84431 | 0.26806 | *rpsH* |
| pdb\|7OE0\|N | 0.88676 | 0.49171 | - |
| WP_001096200.1 | 0.87431 | 0.41871 | *rplE* |
| WP_000729185.1 | 1.00655 | 0.78835 | *rplX* |
| WP_000613955.1 | 0.87788 | 0.43909 | *rplN* |
| pdb\|6H58\|qq | 0.89272 | 0.52854 | - |
| WP_000644741.1 | 0.89748 | 0.55876 | *rpmC* |
| WP_000941212.1 | 0.90928 | 0.63629 | *rplP* |
| pdb\|7OE0\|C | 1.00222 | 0.80759 | - |
| WP_000447529.1 | 1.05715 | 0.58071 | *rplV* |
| pdb\|6H58\|ss | 0.92011 | 0.71023 | - |
| pdb\|6H58\|CC | 0.89626 | 0.55092 | - |
| WP_000617544.1 | 0.80855 | 0.13941 | *rplW* |
| WP_000424395.1 | 0.92512 | 0.74513 | *rplD* |
| WP_000579833.1 | 0.92944 | 0.77552 | *rplC* |
| WP_001181004.1 | 0.96326 | 0.98878 | *rpsJ* |
| WP_000675504.1 | 1.22793 | 0.15588 | *bfr* |
| WP_000031784.1 | 0.93475 | 0.81313 | *tuf* |
| WP_000124700.1 | 0.90292 | 0.59404 | *fusA* |
| pdb\|7OE0\|G | 0.88097 | 0.45708 | - |
| pdb\|7OI0\|L | 0.87828 | 0.44138 | - |
| WP_000903373.1 | 1.42639 | 0.234 | *tusB* |
| WP_000820714.1 | 0.7291 | 0.12524 | *tusC* |
| WP_000091466.1 | 1.02393 | 0.92445 | *DXE50_21575* |
| WP_000838250.1 | 1.08962 | 0.46696 | *A1UI_03620* |
| WP_001153615.1 | 0.87565 | 0.44104 | *slyX* |
| WP_000861334.1 | 0.94707 | 0.90107 | *DXE50_21590* |
| WP_001007730.1 | 1.38017 | 0.17097 | *DXE50_21595* |
| WP_000634798.1 | 0.90114 | 0.44306 | *DXE50_21610* |
| WP_000057356.1 | 0.84479 | 0.40534 | *A1UI_03627* |
| WP_001274680.1 | 1.01082 | 0.98311 | *DXE50_21625* |

| | | | |
|---|---|---|---|
| WP_001148908.1 | 1.08364 | 0.67535 | *EKN05_022235* |
| WP_000242755.1 | 0.9654 | 0.97856 | *GA0061070_104725* |
| WP_000963792.1 | 1.00139 | 0.99822 | *argD* |
| WP_001280641.1 | 1.42287 | 0.23733 | *DXE50_21655* |
| WP_000477225.1 | 1.24037 | 0.13957 | *ppiA* |
| WP_000049208.1 | 1.55275 | 0.06114 | *nirB* |
| WP_000084764.1 | 1.29082 | 0.39558 | *nirD* |
| WP_000349855.1 | 1.12398 | 0.53018 | *cysG* |
| WP_001276847.1 | 1.23564 | 0.48357 | *phoB* |
| WP_001254790.1 | 1.02616 | 0.94652 | *KXD89_01675* |
| WP_000165552.1 | 0.90396 | 0.60089 | *trpS* |
| WP_001031729.1 | 1.04492 | 0.85704 | *gph* |
| WP_000816280.1 | 1.02682 | 0.70108 | *rpe* |
| WP_000742143.1 | 0.80532 | 0.28921 | *O3K_02135* |
| WP_000343215.1 | 0.92126 | 0.5388 | *damX* |
| WP_000439848.1 | 0.93359 | 0.60105 | *aroB* |
| WP_000818618.1 | 0.79295 | 0.10003 | *aroK* |
| WP_001336003.1 | 0.89181 | 0.40155 | *mrcA* |
| WP_000045744.1 | 1.03131 | 0.9012 | *nudE* |
| WP_000104533.1 | 0.94842 | 0.75398 | *igaA* |
| WP_001295168.1 | 0.88688 | 0.48548 | *DXE50_21820* |
| WP_000660483.1 | 0.75215 | 0.16604 | *O3K_02070* |
| WP_001135574.1 | 0.95535 | 0.96026 | *hslO* |
| WP_001265681.1 | 0.92218 | 0.72458 | *pckA* |
| WP_000980727.1 | 1.20771 | 0.18585 | *KXD89_01555* |
| WP_001200455.1 | 1.11135 | 0.73457 | *feoA* |
| WP_000737039.1 | 1.00266 | 0.997 | *feoB* |
| WP_001060070.1 | 0.91055 | 0.63455 | *bioH* |
| WP_000619389.1 | 0.88969 | 0.50969 | *nfuA* |
| WP_001131758.1 | 0.78856 | 0.17061 | *gntT* |
| WP_000815099.1 | 1.05789 | 0.81588 | *DXE50_21955* |
| WP_000371928.1 | 0.7822 | 0.23069 | *glpE* |
| WP_000448136.1 | 0.78828 | 0.09002 | *glpD* |
| WP_000993449.1 | 1.24295 | 0.22903 | *DXE50_21995* |
| WP_001197646.1 | 1.34817 | 0.09544 | *glgA* |
| WP_000253975.1 | 1.35167 | 0.09253 | *glgC* |
| WP_000192523.1 | 1.33964 | 0.21433 | *glgX* |
| WP_000799956.1 | 1.06041 | 0.56859 | *asd* |
| WP_000210111.1 | 0.86368 | 0.46731 | *gntU* |
| WP_000730252.1 | 1.10354 | 0.67933 | *A13A_03767* |
| WP_000639811.1 | 1.30925 | 0.25275 | *DXE50_22045* |
| WP_000236293.1 | 1.02518 | 0.70797 | *DXE50_22050* |
| WP_001295206.1 | 1.35388 | 0.19812 | *G711_04008* |
| WP_000595082.1 | 1.45722 | 0.20658 | *ggt* |
| WP_000825996.1 | 1.58391 | 0.0503 | *EKN05_021790* |
| WP_000073623.1 | 0.97313 | 0.86804 | *ugpQ* |
| WP_000907792.1 | 1.66345 | 0.08645 | *EL76_0318* |
| WP_000803190.1 | 1.37752 | 0.07338 | *A1UI_03729* |

| | | | |
|---|---|---|---|
| WP_000778768.1 | 1.31378 | 0.12859 | *panM* |
| WP_001042003.1 | 0.96195 | 0.81631 | *ftsX* |
| WP_000617723.1 | 0.9832 | 0.89451 | *ftsE* |
| WP_001040654.1 | 0.88946 | 0.39141 | *ftsY* |
| WP_000743193.1 | 1.02243 | 0.93051 | *ECMP0215528_3971* |
| WP_001311191.1 | 1.58098 | 0.12332 | *KXD89_01265* |
| WP_000042895.1 | 0.83379 | 0.37104 | *KXD89_01260* |
| WP_000106551.1 | 0.98919 | 0.90157 | *zntA* |
| WP_000130621.1 | 0.97513 | 0.87731 | *tusA* |
| WP_001245295.1 | 1.34542 | 0.05142 | *dcrB* |
| WP_001190062.1 | 0.84818 | 0.3406 | *nikR* |
| WP_000149156.1 | 0.95467 | 0.80251 | *G711_04052* |
| WP_000361477.1 | 1.22999 | 0.25307 | *KXD89_01155* |
| WP_000439170.1 | 1.07128 | 0.77441 | *KXD89_01130* |
| WP_000902780.1 | 1.04204 | 0.84536 | *DXE50_22455* |
| WP_000323571.1 | 1.1982 | 0.20152 | *uspA* |
| WP_000686620.1 | 0.76146 | 0.11631 | *rsmJ* |
| WP_001298719.1 | 0.91504 | 0.67531 | *HMPREF1595_03282* |
| WP_001300574.1 | 0.88909 | 0.49444 | *rlmJ* |
| WP_000160816.1 | 0.9435 | 0.87554 | *gorA* |
| WP_001296814.1 | 1.0796 | 0.81017 | *DXE50_22570* |
| WP_001298717.1 | 1.33107 | 0.11085 | *hdeB* |
| WP_000756550.1 | 1.09764 | 0.62258 | *hdeA* |
| WP_001081984.1 | 1.4869 | 0.18293 | *mdtE* |
| WP_000784827.1 | 1.06903 | 0.83628 | *ccp* |
| WP_000934216.1 | 1.29964 | 0.38279 | *treF* |
| WP_000357790.1 | 0.90655 | 0.6197 | *KXD89_00965* |
| WP_000191237.1 | 1.34949 | 0.31677 | *A1UI_03828* |
| WP_000037562.1 | 0.81858 | 0.15268 | *WG3_04286* |
| WP_001163141.1 | 0.99616 | 0.97431 | *A1UI_03833* |
| WP_000858214.1 | 0.83714 | 0.20214 | *dctA* |
| WP_001266293.1 | 0.98381 | 0.9148 | *hmsP* |
| WP_000107012.1 | 1.27309 | 0.30583 | *A1UI_03843* |
| WP_001196486.1 | 1.1982 | 0.32058 | *dppD* |
| WP_000084677.1 | 1.34697 | 0.31986 | *dppC* |
| WP_000938855.1 | 1.10226 | 0.68298 | *ECMP0215528_4093* |
| WP_001222883.1 | 1.2582 | 0.11876 | *dppA* |
| WP_001269197.1 | 0.7458 | 0.15404 | *HMPREF1620_03849* |
| WP_000438957.1 | 1.28784 | 0.39997 | *tag* |
| WP_000013950.1 | 1.05959 | 0.81056 | *bisC* |
| WP_000747625.1 | 1.05379 | 0.79552 | *DXE50_22830* |
| WP_000805038.1 | 0.91905 | 0.70289 | *ghrB* |
| WP_000190516.1 | 1.16751 | 0.39813 | *DXE50_22840* |
| WP_000014594.1 | 0.93057 | 0.78347 | *cspA* |
| WP_001291772.1 | 0.93466 | 0.81249 | *glyS* |
| WP_001168560.1 | 0.96849 | 0.96381 | *glyQ* |
| WP_000275334.1 | 1.20383 | 0.54079 | *xylB* |
| WP_001149591.1 | 0.79146 | 0.09675 | *xylA* |

| | | | |
|---|---|---|---|
| WP_000694881.1 | 0.98429 | 0.87449 | *DXE50_22900* |
| WP_000494484.1 | 0.76096 | 0.18365 | *DXE50_22915* |
| WP_000144363.1 | 0.9578 | 0.72916 | *avtA* |
| WP_000514240.1 | 1.14599 | 0.65724 | *A1UI_03880* |
| WP_000164036.1 | 0.77289 | 0.13749 | *O3K_00925* |
| WP_000183980.1 | 1.27853 | 0.17251 | *aldB* |
| WP_000582468.1 | 1.07355 | 0.76751 | *selB* |
| WP_000206275.1 | 0.93824 | 0.70753 | *selA* |
| WP_000779792.1 | 1.07828 | 0.75319 | *DXE50_23010* |
| WP_000072850.1 | 0.91789 | 0.66202 | *O3K_00870* |
| WP_000093247.1 | 1.08089 | 0.68597 | *O3K_00845* |
| WP_000645439.1 | 1.19184 | 0.21262 | *mtlD* |
| WP_000517100.1 | 1.49101 | 0.08929 | *DXE50_23040* |
| WP_000665680.1 | 0.96878 | 0.84792 | *ECMP0215528_4158* |
| WP_000586962.1 | 0.88241 | 0.4656 | *lldD* |
| WP_000932347.1 | 1.14459 | 0.56907 | *trmL* |
| WP_001277561.1 | 1.04224 | 0.86566 | *cysE* |
| WP_001076194.1 | 1.16025 | 0.41834 | *gpsA* |
| WP_000003377.1 | 1.05716 | 0.58069 | *secB* |
| WP_000024392.1 | 0.89496 | 0.41534 | *grxC* |
| WP_001156181.1 | 1.08939 | 0.65339 | *DXE50_23100* |
| WP_001350558.1 | 0.89747 | 0.5587 | *gpmM* |
| AIN33940.1 | 0.96312 | 0.83513 | - |
| WP_000646007.1 | 0.97811 | 0.91828 | *tdh* |
| WP_001213834.1 | 1.07911 | 0.50202 | *kbl* |
| WP_000842820.1 | 1.00178 | 1 | *yibB* |
| WP_000587764.1 | 0.96109 | 0.99921 | *rfaD* |
| WP_000699219.1 | 0.81571 | 0.31785 | *O3K_00725* |
| WP_001264584.1 | 1.02882 | 0.93949 | *rfaC* |
| WP_001395405.1 | 1.083 | 0.80187 | *waaL* |
| WP_001236433.1 | 0.79039 | 0.25056 | *KXD89_00410* |
| WP_000790279.1 | 0.93428 | 0.72421 | *waaZ* |
| WP_000615254.1 | 0.85928 | 0.37966 | *rfaY* |
| WP_000376841.1 | 0.95025 | 0.78548 | *rfaJ* |
| WP_001188013.1 | 0.95948 | 0.82107 | *waaO* |
| WP_000158225.1 | 0.68365 | 0.06584 | *rfaS* |
| WP_000229840.1 | 0.93874 | 0.74126 | *rfaP* |
| WP_000634283.1 | 1.15362 | 0.64098 | *rfaG* |
| WP_000891564.1 | 0.96468 | 0.76658 | *A13A_03970* |
| WP_001171866.1 | 0.94226 | 0.7258 | *coaD* |
| WP_001114543.1 | 1.02474 | 0.92287 | *mutM* |
| pdb\|6H58\|XX | 1.03974 | 0.64823 | - |
| WP_000050139.1 | 0.94117 | 0.64042 | *coaBC* |
| AIN33967.1 | 1.14247 | 0.31657 | - |
| WP_000818601.1 | 0.99555 | 0.93667 | *slmA* |
| WP_000806177.1 | 0.87678 | 0.44543 | *pyrE* |
| WP_000621340.1 | 0.99772 | 0.94863 | *yicC* |
| WP_001295237.1 | 1.0699 | 0.72943 | *gmk* |

| | | | |
|---|---|---|---|
| WP_000135058.1 | 0.85399 | 0.31253 | *rpoZ* |
| WP_000280488.1 | 1.00951 | 0.97379 | *spoT* |
| WP_001070177.1 | 1.03301 | 0.89565 | *trmH* |
| WP_000678419.1 | 1.02854 | 0.94023 | *recG* |
| WP_000468833.1 | 1.19349 | 0.56042 | *gltS* |
| WP_001300954.1 | 1.15594 | 0.54082 | *KXD89_00250* |
| WP_000779426.1 | 1.32521 | 0.34762 | *O3K_00230* |
| WP_001065718.1 | 1.42127 | 0.23886 | *adeD* |
| WP_000879194.1 | 0.92785 | 0.6997 | *uhpT* |
| WP_000633668.1 | 1.089 | 0.72131 | *uhpA* |
| WP_001181706.1 | 0.98572 | 0.92628 | *ilvN* |
| WP_000168475.1 | 0.9053 | 0.46216 | *A1UI_03990* |
| WP_000620888.1 | 0.68015 | 0.06232 | *EKN05_019820* |
| WP_001336371.1 | 1.05761 | 0.86497 | *KXD89_00070* |
| WP_000174305.1 | 1.11733 | 0.72083 | *ECMP0215528_4259* |
| WP_000985549.1 | 0.86907 | 0.3091 | *DXE50_23915* |
| WP_000522208.1 | 1.21669 | 0.4063 | *yidB* |
| WP_000072067.1 | 0.95156 | 0.93319 | *gyrB* |
| WP_000060112.1 | 0.96089 | 0.82651 | *recF* |
| WP_000673464.1 | 0.93826 | 0.62523 | *dnaN* |
| WP_000059111.1 | 1.06317 | 0.75669 | *dnaA* |
| WP_000831330.1 | 0.97128 | 0.80272 | *rpmH* |
| WP_000239730.1 | 1.21397 | 0.41173 | *rnpA* |
| WP_000378250.1 | 0.84944 | 0.29114 | *yidC* |
| WP_001282346.1 | 1.11341 | 0.56615 | *mnmE* |
| WP_001295247.1 | 0.87304 | 0.41161 | *tnaA* |
| WP_000131925.1 | 0.96836 | 0.85535 | *A1UI_04026* |
| WP_000377786.1 | 0.97832 | 0.89208 | *phoU* |
| WP_000063125.1 | 0.9348 | 0.69197 | *pstB* |
| WP_000867146.1 | 0.94294 | 0.64974 | *pstS* |
| WP_000334099.1 | 1.17091 | 0.25267 | *glmS* |
| WP_000933736.1 | 0.89659 | 0.42254 | *glmU* |
| WP_001251965.1 | 0.8857 | 0.48526 | *atpC* |
| WP_000190506.1 | 0.96223 | 0.99373 | *atpD* |
| WP_000896498.1 | 0.92101 | 0.71648 | *atpG* |
| WP_001176745.1 | 0.91515 | 0.67611 | *atpA* |
| WP_001288587.1 | 0.88103 | 0.45741 | *atpH* |
| WP_001052219.1 | 0.96717 | 0.97011 | *atpF* |
| WP_000429386.1 | 1.07013 | 0.53329 | *atpE* |
| WP_000135625.1 | 0.97677 | 0.83296 | *atpB* |
| WP_000932839.1 | 0.92209 | 0.63509 | *rsmG* |
| WP_000499788.1 | 0.90139 | 0.4442 | *mnmG* |
| WP_000432970.1 | 1.33741 | 0.21697 | *asnC* |
| WP_001299914.1 | 1.19735 | 0.44609 | *ravA* |
| WP_000102319.1 | 1.0914 | 0.71429 | *kup* |
| WP_001301979.1 | 0.93772 | 0.62241 | *rbsD* |
| WP_000387770.1 | 0.93046 | 0.58498 | *rbsA* |
| WP_000211858.1 | 0.91696 | 0.61246 | *rbsC* |

| | | | |
|---|---|---|---|
| WP_001056271.1 | 0.94961 | 0.9192 | *rbsB* |
| WP_001300603.1 | 0.90868 | 0.47797 | *rbsK* |
| WP_001131164.1 | 0.96594 | 0.83478 | *DXE50_24215* |
| WP_000379245.1 | 0.88047 | 0.5253 | *hdfR* |
| WP_000841001.1 | 0.85498 | 0.31731 | *maoP* |
| WP_000208520.1 | 1.01053 | 0.98441 | *ilvE* |
| WP_001127399.1 | 1.03617 | 0.88531 | *ilvD* |
| WP_000785596.1 | 0.92619 | 0.69341 | *ilvA* |
| WP_000024939.1 | 0.96088 | 0.74588 | *ilvC* |
| WP_001140251.1 | 0.88481 | 0.37172 | *DXE50_24310* |
| WP_001238899.1 | 0.9461 | 0.76952 | *rep* |
| WP_001295254.1 | 1.03367 | 0.89347 | *gppA* |
| WP_000047499.1 | 0.85214 | 0.30371 | *rhlB* |
| WP_001280776.1 | 0.91794 | 0.69526 | *trxA* |
| WP_001054527.1 | 0.90567 | 0.61216 | *rho* |
| WP_001050960.1 | 1.20489 | 0.5388 | *wecA* |
| WP_001295256.1 | 0.78556 | 0.08647 | *wzzE* |
| WP_001340422.1 | 0.94198 | 0.72453 | *wecB* |
| WP_000006621.1 | 1.00232 | 0.97391 | *wecC* |
| WP_001226601.1 | 0.80121 | 0.20085 | *rffG* |
| WP_000676056.1 | 0.98933 | 0.93598 | *rfbA* |
| WP_000612043.1 | 0.94308 | 0.72954 | *wecE* |
| WP_000217234.1 | 1.0585 | 0.86274 | *rffT* |
| WP_001064040.1 | 0.82746 | 0.27372 | *rffM* |
| WP_000774731.1 | 0.92299 | 0.68127 | *DXE50_24410* |
| WP_000921791.1 | 1.04963 | 0.60932 | *hemY* |
| WP_000138997.1 | 1.04073 | 0.85098 | *hemX* |
| WP_000026046.1 | 1.23467 | 0.48525 | *EL76_4618* |
| WP_001338644.1 | 0.91974 | 0.53129 | *hemC* |
| WP_000999947.1 | 1.11311 | 0.5672 | *cyaY* |
| WP_000799889.1 | 1.0844 | 0.73489 | *DXE50_24485* |
| WP_001160654.1 | 0.72335 | 0.06225 | *dapF* |
| WP_000812796.1 | 1.03929 | 0.8752 | *DXE50_24495* |
| WP_000130691.1 | 0.76496 | 0.19201 | *xerC* |
| WP_001213584.1 | 0.84488 | 0.32944 | *yigB* |
| WP_000383406.1 | 0.98325 | 0.91486 | *uvrD* |
| WP_000947159.1 | 0.976 | 0.88134 | *corA* |
| WP_001277142.1 | 0.85459 | 0.43701 | *DXE50_24560* |
| WP_001259700.1 | 0.97066 | 0.79933 | *pldA* |
| WP_000487654.1 | 0.84225 | 0.39728 | *pldB* |
| WP_000285362.1 | 0.97832 | 0.89208 | *yigL* |
| WP_000153907.1 | 0.96952 | 0.95893 | *metE* |
| WP_001336442.1 | 1.07591 | 0.70551 | *EL76_4589* |
| WP_000045177.1 | 1.21171 | 0.17956 | *O3K_24700* |
| WP_000347714.1 | 0.99057 | 0.94072 | *rmuC* |
| WP_000227958.1 | 0.97855 | 0.84276 | *ubiE* |
| WP_001295259.1 | 0.81431 | 0.31392 | *ubiJ* |
| WP_000187530.1 | 1.74178 | 0.0613 | *ubiB* |

| | | | |
|---|---|---|---|
| WP_001295260.1 | 1.14703 | 0.45699 | *tatA* |
| WP_000459594.1 | 1.04407 | 0.85976 | *tatB* |
| WP_001192396.1 | 1.17325 | 0.60036 | *rfaH* |
| WP_000339804.1 | 1.06208 | 0.76118 | *ubiD* |
| WP_000209826.1 | 0.83143 | 0.18594 | *DXE50_24695* |
| WP_000438725.1 | 1.34502 | 0.09812 | *fadA* |
| WP_000965936.1 | 1.33806 | 0.1043 | *fadB* |
| WP_000444561.1 | 0.97388 | 0.93822 | *pepQ* |
| WP_000853959.1 | 1.17761 | 0.4896 | *hemG* |
| WP_001295263.1 | 1.26195 | 0.19719 | *DXE50_24765* |
| WP_001065497.1 | 1.10559 | 0.67355 | *srkA* |
| WP_000725337.1 | 0.94447 | 0.88245 | *dsbA* |
| WP_000250006.1 | 0.92013 | 0.71038 | *polA* |
| WP_000183349.1 | 1.04763 | 0.82146 | *engB* |
| WP_001295266.1 | 0.93087 | 0.58707 | *yihI* |
| WP_000116090.1 | 1.17975 | 0.36569 | *hemN* |
| WP_001188777.1 | 0.84124 | 0.31731 | *ntrC* |
| WP_000190577.1 | 0.85284 | 0.30705 | *DXE50_24825* |
| WP_001271717.1 | 0.76894 | 0.0564 | *DXE50_24830* |
| WP_000570668.1 | 0.97764 | 0.92051 | *typA* |
| WP_000059678.1 | 0.82726 | 0.27309 | *A13A_04267* |
| WP_001295269.1 | 1.25771 | 0.20396 | *yihX* |
| WP_000560983.1 | 0.993 | 0.95981 | *dtd* |
| WP_001297068.1 | 1.23809 | 0.3655 | *WG3_04779* |
| WP_000027703.1 | 1.00344 | 0.98006 | *fdhE* |
| WP_000331377.1 | 0.97244 | 0.86488 | *fdxH* |
| WP_010723259.1 | 0.97146 | 0.94968 | *fdnG* |
| WP_000753617.1 | 0.96541 | 0.84395 | *fdhD* |
| WP_000122641.1 | 0.8421 | 0.25849 | *sodA* |
| WP_001270260.1 | 0.94483 | 0.73754 | *O3K_24310* |
| WP_000580417.1 | 1.04733 | 0.89125 | *cpxA* |
| WP_001033722.1 | 1.2504 | 0.1275 | *cpxR* |
| WP_001223800.1 | 1.68428 | 0.07895 | *cpxP* |
| WP_000591795.1 | 0.97586 | 0.92886 | *pfkA* |
| WP_001326656.1 | 0.84512 | 0.33023 | *cdh* |
| WP_001216325.1 | 1.12072 | 0.37331 | *tpiA* |
| WP_000802216.1 | 0.92057 | 0.67212 | *KXD89_21165* |
| WP_000655989.1 | 1.21711 | 0.40547 | *EL76_4498* |
| WP_000323556.1 | 1.14243 | 0.47103 | *ECMP0215528_4537* |
| WP_000796332.1 | 1.08063 | 0.74616 | *EL76_4496* |
| WP_001250644.1 | 1.09513 | 0.63186 | *glpX* |
| WP_000136788.1 | 0.96104 | 0.99946 | *glpK* |
| WP_000084268.1 | 1.06594 | 0.7908 | *glpF* |
| WP_001296623.1 | 1.10785 | 0.41024 | *zapB* |
| WP_000872908.1 | 1.18725 | 0.34681 | *rraA* |
| WP_000139496.1 | 0.79352 | 0.1821 | *menA* |
| WP_001293341.1 | 0.99644 | 0.83362 | *hslU* |
| WP_000208242.1 | 0.94161 | 0.86197 | *hslV* |

| | | | |
|---|---|---|---|
| WP_000068828.1 | 0.93849 | 0.62641 | *ftsN* |
| WP_000644904.1 | 1.04259 | 0.86454 | *cytR* |
| WP_000710769.1 | 0.85827 | 0.33349 | *rpmE* |
| WP_000852812.1 | 0.84708 | 0.23247 | *metJ* |
| WP_001295694.1 | 0.97075 | 0.85702 | *A1UI_04255* |
| WP_000110772.1 | 0.91584 | 0.51223 | *metL* |
| WP_001295695.1 | 1.25087 | 0.12696 | *katG* |
| WP_000374004.1 | 1.13642 | 0.48978 | *gldA* |
| WP_000424840.1 | 0.94486 | 0.73769 | *fsa* |
| WP_001174077.1 | 0.9909 | 0.94198 | *ptsP* |
| WP_000556306.1 | 1.00127 | 0.96817 | *KXD89_20995* |
| WP_001005586.1 | 0.98823 | 0.87116 | *ppc* |
| WP_001298964.1 | 0.96363 | 0.76084 | *argE* |
| WP_000935370.1 | 1.07998 | 0.80925 | *argC* |
| WP_001302318.1 | 1.26599 | 0.31724 | *argB* |
| WP_001230087.1 | 1.21473 | 0.28391 | *argH* |
| WP_001025939.1 | 1.01603 | 0.95967 | *oxyR* |
| WP_001120810.1 | 0.90856 | 0.63145 | *sthA* |
| WP_000187022.1 | 0.82445 | 0.16732 | *trmA* |
| WP_000201820.1 | 0.9481 | 0.75251 | *murI* |
| WP_001016699.1 | 1.04247 | 0.86491 | *murB* |
| WP_000654630.1 | 0.89824 | 0.53221 | *birA* |
| WP_000023081.1 | 0.98391 | 0.87239 | *coaA* |
| WP_001275702.1 | 0.96292 | 0.82079 | *secE* |
| WP_001287516.1 | 0.88433 | 0.47706 | *nusG* |
| WP_001085926.1 | 0.87086 | 0.39951 | *rplK* |
| WP_001096684.1 | 0.92014 | 0.71044 | *rplA* |
| WP_001207201.1 | 0.89316 | 0.53131 | *rplJ* |
| WP_000028878.1 | 0.87236 | 0.40784 | *rplL* |
| WP_000263098.1 | 0.9396 | 0.84766 | *rpoB* |
| WP_000653944.1 | 0.9235 | 0.73383 | *rpoC* |
| WP_000934302.1 | 0.85382 | 0.3602 | *rsd* |
| WP_000373940.1 | 1.19193 | 0.45774 | *nudC* |
| WP_000137657.1 | 0.95953 | 0.73854 | *hemE* |
| WP_000362388.1 | 1.27143 | 0.42491 | *nfi* |
| WP_001044513.1 | 1.02907 | 0.69173 | *hupA* |
| WP_000866800.1 | 0.97177 | 0.86178 | *purD* |
| WP_001187559.1 | 0.93359 | 0.60104 | *purH* |
| WP_000138905.1 | 1.23893 | 0.14138 | *aceB* |
| WP_000857856.1 | 1.21685 | 0.17175 | *aceA* |
| WP_001137220.1 | 1.75684 | 0.05735 | *aceK* |
| WP_000632913.1 | 0.9388 | 0.74149 | *espL4* |
| WP_000096011.1 | 0.86761 | 0.3036 | *metH* |
| WP_000956830.1 | 1.53326 | 0.15087 | *DXE50_25790* |
| WP_000421763.1 | 1.31647 | 0.12569 | *pepE* |
| WP_000936377.1 | 1.12497 | 0.7035 | *A1UI_04335* |
| WP_001207634.1 | 0.81575 | 0.31796 | *A1UI_04336* |
| WP_001290310.1 | 0.78624 | 0.1654 | *A1UI_04337* |

| | | | |
|---|---|---|---|
| WP_000789986.1 | 0.98235 | 0.89847 | *pgi* |
| WP_001326644.1 | 0.94505 | 0.76547 | *ubiC* |
| WP_000455227.1 | 0.96002 | 0.82318 | *ubiA* |
| WP_000017354.1 | 0.9455 | 0.8898 | *plsB* |
| WP_000646078.1 | 0.90332 | 0.45301 | *lexA* |
| WP_001030593.1 | 1.18209 | 0.35971 | *DXE50_25970* |
| WP_001295691.1 | 0.89907 | 0.59217 | *zur* |
| WP_001298868.1 | 1.01929 | 0.94508 | *dusA* |
| WP_000235508.1 | 1.24244 | 0.137 | *KXD89_20510* |
| WP_000918363.1 | 0.90162 | 0.54644 | *DXE50_26005* |
| WP_001147328.1 | 0.96129 | 0.81327 | *alr* |
| WP_000486985.1 | 0.8222 | 0.16159 | *tyrB* |
| WP_001226928.1 | 0.89197 | 0.40222 | *DXE50_26025* |
| WP_000270375.1 | 1.53183 | 0.15177 | *DXE50_26030* |
| WP_000155657.1 | 1.06859 | 0.73471 | *DXE50_26035* |
| WP_000357740.1 | 1.08656 | 0.66415 | *uvrA* |
| WP_000168305.1 | 0.93996 | 0.85024 | *DXE50_26045* |
| WP_000106882.1 | 0.91278 | 0.64289 | *DXE50_26070* |
| WP_000402210.1 | 0.78851 | 0.24592 | *KY274_20395* |
| WP_000832573.1 | 1.30142 | 0.26353 | *actP* |
| WP_000078239.1 | 1.31105 | 0.07216 | *acs* |
| WP_000719886.1 | 0.87946 | 0.52174 | *DXE50_26140* |
| WP_001046187.1 | 0.87273 | 0.42977 | *alsB* |
| WP_000083216.1 | 0.88225 | 0.36111 | *rpiR* |
| WP_000716794.1 | 0.99788 | 0.98217 | *rpiB* |
| WP_000971886.1 | 0.97679 | 0.88784 | *phnN* |
| WP_001300891.1 | 1.10621 | 0.59152 | *DXE50_26305* |
| WP_000697926.1 | 1.10811 | 0.74209 | *basR* |
| WP_000986601.1 | 1.19155 | 0.56417 | *A1UI_04504* |
| WP_000611288.1 | 0.93337 | 0.72071 | *dcuR* |
| WP_001295383.1 | 1.0355 | 0.8736 | *DXE50_26450* |
| WP_001188520.1 | 1.26996 | 0.18492 | *DXE50_26655* |
| WP_000068922.1 | 1.34184 | 0.32623 | *dsbD* |
| WP_000883400.1 | 0.82981 | 0.359 | *cutA* |
| WP_000961959.1 | 0.809 | 0.22104 | *DXE50_26670* |
| WP_000069437.1 | 1.11226 | 0.39729 | *aspA* |
| WP_001267448.1 | 1.12092 | 0.71265 | *ECMP0215528_4825* |
| WP_001026276.1 | 0.88378 | 0.47372 | *groES* |
| WP_000729117.1 | 0.89587 | 0.54847 | *groL* |
| WP_000558209.1 | 1.17186 | 0.38638 | *DXE50_26700* |
| WP_000940549.1 | 0.85358 | 0.35933 | *epmB* |
| WP_000257278.1 | 0.77503 | 0.06571 | *efp* |
| WP_000239596.1 | 1.16026 | 0.41832 | *ecnB* |
| WP_001238378.1 | 1.34048 | 0.21335 | *O3K_22885* |
| WP_001336292.1 | 1.03272 | 0.8966 | *KXD89_19985* |
| WP_000208757.1 | 1.06028 | 0.85821 | *frdC* |
| WP_000829498.1 | 1.13406 | 0.59616 | *DXE50_26755* |
| WP_001192973.1 | 1.21488 | 0.28358 | *frdA* |

| | | | |
|---|---|---|---|
| WP_000004771.1 | 0.91149 | 0.58866 | *epmA* |
| WP_001236847.1 | 1.05718 | 0.81811 | *ECMP0215528_4846* |
| WP_000934920.1 | 1.02532 | 0.91827 | *psd* |
| WP_000041964.1 | 0.99428 | 0.92963 | *rsgA* |
| WP_001295188.1 | 1.00127 | 0.96815 | *orn* |
| WP_001295189.1 | 1.16889 | 0.50979 | *nnrD* |
| WP_000981977.1 | 0.8986 | 0.53373 | *tsaE* |
| WP_001280345.1 | 1.14091 | 0.57844 | *miaA* |
| WP_001051883.1 | 1.08883 | 0.46955 | *hfq* |
| WP_000460362.1 | 1.02138 | 0.93575 | *hflX* |
| WP_000312488.1 | 0.9286 | 0.7696 | *hflK* |
| WP_001232412.1 | 1.02873 | 0.69312 | *hflC* |
| WP_000527955.1 | 1.03679 | 0.66011 | *purA* |
| WP_001177639.1 | 0.87691 | 0.5128 | *nsrR* |
| WP_000076332.1 | 1.16589 | 0.40259 | *rnr* |
| WP_001293282.1 | 0.94848 | 0.75425 | *rlmB* |
| WP_000569707.1 | 1.23755 | 0.4803 | *yjfP* |
| WP_000133631.1 | 1.11498 | 0.72621 | *ulaR* |
| WP_001216676.1 | 0.8051 | 0.12987 | *rpsF* |
| WP_001315977.1 | 1.12181 | 0.71064 | *priB* |
| pdb\|7OI0\|R | 1.11373 | 0.39307 | - |
| WP_001196062.1 | 0.99246 | 0.85177 | *rplI* |
| WP_001119478.1 | 1.16893 | 0.50969 | *ytfB* |
| WP_000211225.1 | 0.8875 | 0.49622 | *DXE50_27040* |
| WP_000331456.1 | 0.72817 | 0.06779 | *ytfE* |
| WP_000560561.1 | 1.00078 | 0.99546 | *A13A_04864* |
| WP_000589431.1 | 1.0111 | 0.76833 | *cpdB* |
| WP_000886919.1 | 1.05202 | 0.80295 | *cysQ* |
| WP_000175279.1 | 1.01885 | 0.94706 | *KXD89_19650* |
| WP_000935036.1 | 0.86156 | 0.46018 | *WG3_00121* |
| WP_001295196.1 | 1.21032 | 0.29335 | *msrA* |
| WP_001269327.1 | 1.17298 | 0.50024 | *tamA* |
| WP_000060911.1 | 1.2019 | 0.31206 | *tamB* |
| WP_001219160.1 | 1.03037 | 0.90429 | *DXE50_27115* |
| WP_000055075.1 | 0.77357 | 0.06338 | *ppa* |
| WP_000265913.1 | 1.40195 | 0.05863 | *ytfQ* |
| WP_001219813.1 | 0.8735 | 0.32606 | *mpl* |
| WP_000166270.1 | 1.32256 | 0.23526 | *yjgA* |
| WP_001162171.1 | 0.99867 | 0.82356 | *pmbA* |
| WP_000187778.1 | 1.08167 | 0.74304 | *nrdD* |
| WP_000047539.1 | 0.813 | 0.15244 | *DXE50_27225* |
| WP_000230281.1 | 0.83127 | 0.3634 | *ECMP0215528_4971* |
| WP_000256658.1 | 0.79873 | 0.27177 | *KXD89_19490* |
| WP_000036434.1 | 0.95965 | 0.82173 | *KY274_19440* |
| WP_000002953.1 | 1.04861 | 0.81733 | *rraB* |
| WP_001059402.1 | 0.91775 | 0.61596 | *A311_00387* |
| WP_000416392.1 | 0.85985 | 0.34141 | *valS* |
| WP_000786400.1 | 0.92355 | 0.68339 | *holC* |

| | | | |
|---|---|---|---|
| WP_000397144.1 | 0.93057 | 0.7835 | *pepA* |
| WP_000584114.1 | 0.94931 | 0.75809 | *lptF* |
| WP_001295681.1 | 0.96392 | 0.82543 | *lptG* |
| WP_001294573.1 | 1.31522 | 0.12702 | *HMPREF1599_05895* |
| WP_000998695.1 | 1.07846 | 0.81296 | *O3K_22310* |
| WP_000175457.1 | 0.90998 | 0.58215 | *DXE50_19750* |
| WP_000373366.1 | 0.98522 | 0.92398 | *A13A_04969* |
| WP_000695564.1 | 1.57182 | 0.12821 | *HMPREF1589_00471* |
| WP_000438562.1 | 0.9742 | 0.81881 | *uxuA* |
| WP_000208205.1 | 0.94874 | 0.68047 | *HMPREF1589_00461* |
| WP_000833679.1 | 0.85451 | 0.36263 | *A1UI_04679* |
| WP_000568432.1 | 0.97406 | 0.81802 | *iadA* |
| WP_001300012.1 | 0.78565 | 0.23895 | *HMPREF1589_00446* |
| WP_000443951.1 | 0.68574 | 0.068 | *mcrB* |
| WP_001272447.1 | 0.78879 | 0.17112 | *HMPREF1589_00433* |
| WP_001063204.1 | 0.7933 | 0.1008 | *HMPREF1589_00432* |
| WP_000217931.1 | 0.93468 | 0.69142 | *HMPREF1589_00430* |
| WP_001312515.1 | 0.83956 | 0.20929 | *O3K_21780* |
| WP_001299714.1 | 1.10656 | 0.59025 | *G925_04708* |
| WP_001308243.1 | 0.90539 | 0.61541 | *DXE50_27710* |
| WP_000799911.1 | 1.19885 | 0.55019 | *dnaC* |
| WP_000098818.1 | 0.92599 | 0.6524 | *dnaT* |
| WP_001272330.1 | 1.06131 | 0.76434 | *rsmC* |
| WP_000204012.1 | 1.08804 | 0.78965 | *G925_04726* |
| WP_001092461.1 | 1.03978 | 0.91079 | *rimI* |
| WP_000175940.1 | 0.9036 | 0.59852 | *prfC* |
| WP_001295748.1 | 0.99616 | 0.8349 | *osmY* |
| WP_001143230.1 | 1.39714 | 0.15523 | - |
| WP_001298497.1 | 1.05928 | 0.77269 | *deoC* |
| WP_000477807.1 | 1.08353 | 0.67577 | *deoA* |
| WP_000816471.1 | 1.11983 | 0.37577 | *deoB* |
| WP_000224877.1 | 1.05546 | 0.58708 | *deoD* |
| WP_000105884.1 | 0.95279 | 0.77406 | *lplA* |
| WP_001132955.1 | 0.94751 | 0.74982 | *DXE50_28030* |
| WP_001029687.1 | 0.94496 | 0.66038 | *radA* |
| WP_000093814.1 | 0.92832 | 0.6628 | *nadR* |
| WP_000046749.1 | 1.00318 | 0.80331 | *ettA* |
| WP_000409451.1 | 1.28192 | 0.29208 | *slt* |
| WP_000068679.1 | 1.08961 | 0.71953 | *trpR* |
| WP_000942344.1 | 0.81034 | 0.22464 | *gpmB* |
| WP_000371666.1 | 1.26923 | 0.18601 | *robA* |
| WP_000875487.1 | 1.02185 | 0.93369 | *creA* |
| WP_001194358.1 | 1.17348 | 0.24746 | *arcA* |
| WP_001223132.1 | 1.0654 | 0.79249 | *trmJ* |
| QBM92672.2 | 0.97589 | 0.88437 | - |
| WP_000737282.1 | 0.81067 | 0.30378 | - |
| WP_000875061.1 | 1.11914 | 0.71669 | - |
| WP_001306825.1 | 1.16561 | 0.61595 | *ycgN* |

| | | | |
|---|---|---|---|
| WP_001350503.1 | 0.79271 | 0.09857 | - |
| WP_001307844.1 | 1.02113 | 0.93485 | *rnd* |
| QWY90515.1 | 0.71325 | 0.10146 | - |
| QWY90557.1 | 0.96193 | 0.83051 | - |

The up-regulated proteins are highlighted in orange, while blue for down-regulated proteins. Target genes for analysis are in red.

| Table S5. The differentially expressed proteins between *E. coli* BW25113 carrying WT MCR-1 and M6 | | | | | | | | | | |
|---|---|---|---|---|---|---|---|---|---|---|
| The significantly enriched GO terms of differentially expressed proteins between *E. coli* BW25113 carrying WT MCR-1 and M6 | | | | | | | | | | |
| GO_ID | Term | Test | Ref | TestAll | RefAll | Test_per | Ref_per | P value | FDR | Rich Factor |
| GO:0030313 | cell envelope | 17 | 172 | 97 | 2350 | 0.175257732 | 0.073191489 | 0.0004768 | 0.16655752 | 0.098837209 |
| GO:0031975 | envelope | 17 | 175 | 97 | 2350 | 0.175257732 | 0.074468085 | 0.000585571 | 0.16655752 | 0.097142857 |
| GO:0030288 | outer membrane-bounded periplasmic space | 11 | 103 | 97 | 2350 | 0.113402062 | 0.043829787 | 0.00282408 | 0.344537731 | 0.106796117 |
| GO:0042597 | periplasmic space | 12 | 127 | 97 | 2350 | 0.12371134 | 0.054042553 | 0.005088546 | 0.349201477 | 0.094488189 |
| GO:0071944 | cell periphery | 32 | 559 | 97 | 2350 | 0.329896907 | 0.23787234 | 0.02272628 | 0.387364975 | 0.057245081 |
| GO:0019867 | outer membrane | 7 | 70 | 97 | 2350 | 0.072164948 | 0.029787234 | 0.023382679 | 0.387364975 | 0.1 |
| GO:0009279 | cell outer membrane | 6 | 63 | 97 | 2350 | 0.06185567 | 0.026808511 | 0.04304829 | 0.397201869 | 0.095238095 |
| GO:0044462 | external encapsulating structure part | 6 | 63 | 97 | 2350 | 0.06185567 | 0.026808511 | 0.04304829 | 0.397201869 | 0.095238095 |
| GO:0009338 | exodeoxyribonuclease V complex | 1 | 2 | 97 | 2350 | 0.010309278 | 0.000851064 | 0.080866281 | 0.446419352 | 0.5 |
| GO:0009289 | pilus | 1 | 2 | 97 | 2350 | 0.010309278 | 0.000851064 | 0.080866281 | 0.446419352 | 0.5 |
| GO:0030312 | external encapsulating structure | 6 | 75 | 97 | 2350 | 0.06185567 | 0.031914894 | 0.086215692 | 0.459912075 | 0.08 |
| GO:0005886 | plasma membrane | 26 | 502 | 97 | 2350 | 0.268041237 | 0.213617021 | 0.114912024 | 0.499936259 | 0.051792829 |
| GO:0016021 | integral component of membrane | 24 | 472 | 97 | 2350 | 0.24742268 | 0.200851064 | 0.149410422 | 0.53946267 | 0.050847458 |
| GO:0031224 | intrinsic component of membrane | 24 | 487 | 97 | 2350 | 0.24742268 | 0.207234043 | 0.190702895 | 0.572108684 | 0.049281314 |
| GO:0016020 | membrane | 33 | 696 | 97 | 2350 | 0.340206186 | 0.296170213 | 0.194793149 | 0.581203473 | 0.047413793 |
| GO:0044425 | membrane part | 25 | 524 | 97 | 2350 | 0.257731959 | 0.222978723 | 0.234081901 | 0.630392568 | 0.047709924 |
| GO:0046930 | pore complex | 1 | 7 | 97 | 2350 | 0.010309278 | 0.002978723 | 0.255806449 | 0.642735654 | 0.142857143 |
| GO:0045203 | integral component of cell outer membrane | 1 | 8 | 97 | 2350 | 0.010309278 | 0.003404255 | 0.286615998 | 0.673885152 | 0.125 |
| GO:0042995 | cell projection | 1 | 8 | 97 | 2350 | 0.010309278 | 0.003404255 | 0.286615998 | 0.673885152 | 0.125 |
| GO:0043231 | intracellular membrane-bounded organelle | 1 | 10 | 97 | 2350 | 0.010309278 | 0.004255319 | 0.344497641 | 0.703082545 | 0.1 |

**Table S6. The differentially expressed proteins between *E. coli* BW25113 carrying WT MCR-1 and M6**

| The KEGG pathways map of differentially expressed proteins between *E. coli* BW25113 carrying WT MCR-1 and M6 | | | | | | | | | | |
|---|---|---|---|---|---|---|---|---|---|---|
| **Map_ID** | **Pathways** | **Test** | **TestAll** | **Ref** | **RefAll** | **Test_per** | **Ref_per** | **P value** | **FDR** | **Rich Factor** |
| ko02026 | Biofilm formation - Escherichia coli | 3 | 97 | 28 | 2350 | 0.030928 | 0.011915 | 0.106128204 | 0.742897429 | 0.107142857 |
| ko04216 | Ferroptosis | 1 | 97 | 2 | 2350 | 0.010309 | 0.000851 | 0.080866281 | 0.742897429 | 0.5 |
| ko00040 | Pentose and glucuronate interconversions | 2 | 97 | 19 | 2350 | 0.020619 | 0.008085 | 0.183464562 | 0.791519469 | 0.105263158 |
| ko00550 | Peptidoglycan biosynthesis | 2 | 97 | 21 | 2350 | 0.020619 | 0.008936 | 0.214007878 | 0.791519469 | 0.095238095 |
| ko00330 | Arginine and proline metabolism | 2 | 97 | 22 | 2350 | 0.020619 | 0.009362 | 0.229463379 | 0.791519469 | 0.090909091 |
| ko00790 | Folate biosynthesis | 2 | 97 | 23 | 2350 | 0.020619 | 0.009787 | 0.244994121 | 0.791519469 | 0.086956522 |
| ko00633 | Nitrotoluene degradation | 1 | 97 | 6 | 2350 | 0.010309 | 0.002553 | 0.223680603 | 0.791519469 | 0.166666667 |
| ko00051 | Fructose and mannose metabolism | 2 | 97 | 27 | 2350 | 0.020619 | 0.011489 | 0.307262512 | 0.807041971 | 0.074074074 |
| ko00670 | One carbon pool by folate | 1 | 97 | 11 | 2350 | 0.010309 | 0.004681 | 0.371670174 | 0.807041971 | 0.090909091 |
| ko00410 | beta-Alanine metabolism | 1 | 97 | 11 | 2350 | 0.010309 | 0.004681 | 0.371670174 | 0.807041971 | 0.090909091 |
| ko00220 | Arginine biosynthesis | 1 | 97 | 12 | 2350 | 0.010309 | 0.005106 | 0.397727461 | 0.807041971 | 0.083333333 |
| ko03010 | Ribosome | 1 | 97 | 56 | 2350 | 0.010309 | 0.02383 | 0.908299313 | 0.908299313 | 0.017857143 |
| ko00270 | Cysteine and methionine metabolism | 1 | 97 | 34 | 2350 | 0.010309 | 0.014468 | 0.763916442 | 0.843534434 | 0.029411765 |
| ko00630 | Glyoxylate and dicarboxylate metabolism | 1 | 97 | 29 | 2350 | 0.010309 | 0.01234 | 0.707686248 | 0.825633956 | 0.034482759 |
| ko00052 | Galactose metabolism | 1 | 97 | 22 | 2350 | 0.010309 | 0.009362 | 0.606083889 | 0.807041971 | 0.045454545 |
| ko00920 | Sulfur metabolism | 1 | 97 | 18 | 2350 | 0.010309 | 0.00766 | 0.533065071 | 0.807041971 | 0.055555556 |
| ko00561 | Glycerolipid metabolism | 1 | 97 | 14 | 2350 | 0.010309 | 0.005957 | 0.446675733 | 0.807041971 | 0.071428571 |
| ko00450 | Selenocompound metabolism | 1 | 97 | 13 | 2350 | 0.010309 | 0.005532 | 0.422714816 | 0.807041971 | 0.076923077 |
| ko00260 | Glycine, serine and threonine metabolism | 2 | 97 | 35 | 2350 | 0.020619 | 0.014894 | 0.427731824 | 0.807041971 | 0.057142857 |
| ko00511 | Other glycan degradation | 1 | 97 | 2 | 2350 | 0.010309 | 0.000851 | 0.080866281 | 0.742897429 | 0.5 |
| ko00910 | Nitrogen metabolism | 2 | 97 | 16 | 2350 | 0.020619 | 0.006809 | 0.139178119 | 0.751493081 | 0.125 |

The up-regulated pathways are highlighted in red, while blue for down-regulated pathways.
