## Supplementary material for "Identification of a novel MCR-1 variant displaying low level of co-resistance to β-lactam antibiotics that uncovers a potential novel antimicrobial peptide": Table S4-10: Table S7-10 strains, plasmid and primers .pdf

**Table S7. Bacterial strains used in this study**

| Bacterial strains | Description | Reference/source |
|---|---|---|
| *E.coli* DH5α | Type strain (F-, φ80d lacZ ΔM15, Δ(lacZYA argF) U169, deoR , recA1 , endA1 , hsdR17 (rK-, mK+), phoA, supE44 , λ-, thi-1, gyrA96 , relA1) | Our lab |
| *E.coli* BW25113 | Type strain (F- DE(araD-araB)567 lacZ4787(del)::rrnB-3 LAM-rph-1 DE(rhaD-rhaB)568 hsdR514) | Our lab |
| *E.coli* ATCC25922 | Type strain | Our lab |
| *P. aeruginosa* ATCC27853 | Type strain | Our lab |
| *S. Typhimurium* SL1344 | Type strain | Our lab |
| *K. pneumoniae* ATCC13883 | Type strain | Our lab |
| *A. baumannii* ATCC19606 | Type strain | Our lab |
| *S. aureus* ATCC25913 | Type strain | Our lab |
| *A. baumannii* clinical isolate 1 | Clinical isolate | Our lab |
| *A. baumannii* clinical isolate 2 | Clinical isolate | Our lab |
| *E. coli* CRE clinical isolate 1 | Clinical isolate | Our lab |
| *K. pneumoniae* CRKP clinical isolate 1 | Clinical isolate | Our lab |
| *S. aureus* MRSA clinical isolate 1 | Clinical isolate | Our lab |
| *S. aureus* MRSA clinical isolate 2 | Clinical isolate | Our lab |
| *E. coli* $mcr\text{-}1^{+}$ clinical isolate 1 | Clinical isolate | Our lab |
| *E. coli* $mcr\text{-}1^{+}$ clinical isolate 2 | Clinical isolate | Our lab |
| *E.coli* BW25113 Δ *mrcB* | *E. coli* BW25113 derivative with the deletion of *mrcB* | This study |
| *E.coli* BW25113 Δ *ycbB* | *E. coli* BW25113 derivative with the deletion of *ycbB* | This study |
| *E.coli* BW25113 Δ *ycfM* | *E. coli* BW25113 derivative with the deletion of *ycfM* | This study |

**Table S8. Plasmids used in this study**

| Plasmids | Description | Reference/source |
|---|---|---|
| pACYCDuet-1 | Vector carrying the P15A replicon, *lacI* gene and chloramphenicol resistance gene | Addgene #71147 |
| pBAD24 | A tightly controlled expression vectors regulated by the arabinose operon, carrying pBR322 replicon and ampicillin gene. | Our lab |
| pACYC-Para-*mcr-1* | pACYCDuet-1 derivative replacing T7 promoter with arabinose promoter and carrying wide type mcr-1 gene. | Our lab |
| pACYC-Para-M6 | pACYCDuet-1 derivative replacing T7 promoter with arabinose promoter and carrying mcr-1 gene with mutations P188A+P195S. | This study |
| pACYC-Para-*mcr-1* P188A | pACYCDuet-1 derivative replacing T7 promoter with arabinose promoter and carrying mcr-1 gene with mutation P188A. | This study |
| pACYC-Para-*mcr-1* P195S | pACYCDuet-1 derivative replacing T7 promoter with arabinose promoter and carrying mcr-1 gene with mutation P195S. | This study |
| pACYC-NP-*mcr-1* | pACYCDuet-1 derivative replacing T7 promoter with native promoter of *mcr-1* and carrying wide type mcr-1 gene. | Our lab |
| pACYC-NP-M6 | pACYCDuet-1 derivative replacing T7 promoter with native promoter of *mcr-1* and carrying mcr-1 gene with mutations P188A+P195S. | This study |
| pACYC-Para-*mcr-1* K187A | pACYCDuet-1 derivative replacing T7 promoter with arabinose promoter and carrying mcr-1 gene with mutation K187A. | This study |
| pACYC-Para-*mcr-1* L189A | pACYCDuet-1 derivative replacing T7 promoter with arabinose promoter and carrying mcr-1 gene with mutation L189A. | This study |
| pACYC-Para-*mcr-1* R190A | pACYCDuet-1 derivative replacing T7 promoter with arabinose promoter and carrying mcr-1 gene with mutation R190A. | This study |
| pACYC-Para-*mcr-1* S191A | pACYCDuet-1 derivative replacing T7 promoter with arabinose promoter and carrying mcr-1 gene with mutation S191A. | This study |
| pACYC-Para-*mcr-1* Y192A | pACYCDuet-1 derivative replacing T7 promoter with arabinose promoter and carrying mcr-1 gene with mutation Y192A. | This study |
| pACYC-Para-*mcr-1* V193A | pACYCDuet-1 derivative replacing T7 promoter with arabinose promoter and carrying mcr-1 gene with mutation V193A. | This study |
| pACYC-Para-*mcr-1* N194A | pACYCDuet-1 derivative replacing T7 promoter with arabinose promoter and carrying mcr-1 gene with mutation N194A. | This study |
| pACYC-Para-*mcr-1* P195A | pACYCDuet-1 derivative replacing T7 promoter with arabinose promoter and carrying mcr-1 gene with mutation P195A. | This study |
| pACYC-Para-*mcr-1* I196A | pACYCDuet-1 derivative replacing T7 promoter with arabinose promoter and carrying mcr-1 gene with mutation I196A. | This study |
| pACYC-Para-*mcr-1* M197A | pACYCDuet-1 derivative replacing T7 promoter with arabinose promoter and carrying mcr-1 gene with mutation M197A. | This study |
| pACYC-Para-*mcr-1* P198A | pACYCDuet-1 derivative replacing T7 promoter with arabinose promoter and carrying mcr-1 gene with mutation P198A. | This study |
| pACYC-Para-*mcr-1* I199A | pACYCDuet-1 derivative replacing T7 promoter with arabinose promoter and carrying mcr-1 gene with mutation I199A. | This study |
| pACYC-Para-*mcr-1* Y200A | pACYCDuet-1 derivative replacing T7 promoter with arabinose promoter and carrying mcr-1 gene with mutation Y200A. | This study |
| pACYC-Para-*mcr-1* S201A | pACYCDuet-1 derivative replacing T7 promoter with arabinose promoter and carrying mcr-1 gene with mutation S201A. | This study |
| pACYC-Para-*mcr-1* V202A | pACYCDuet-1 derivative replacing T7 promoter with arabinose promoter and carrying mcr-1 gene with mutation V202A. | This study |
| pACYC-Para-*mcr-1* G203A | pACYCDuet-1 derivative replacing T7 promoter with arabinose promoter and carrying mcr-1 gene with mutation G203A. | This study |
| pACYC-Para-*mcr-1* K204A | pACYCDuet-1 derivative replacing T7 promoter with arabinose promoter and carrying mcr-1 gene with mutation K204A. | This study |
| pACYC-Para-*mcr-1* L205A | pACYCDuet-1 derivative replacing T7 promoter with arabinose promoter and carrying mcr-1 gene with mutation L205A. | This study |
| pACYC-Para-*mcr-1* M197E | pACYCDuet-1 derivative replacing T7 promoter with arabinose promoter and carrying mcr-1 gene with mutation M197E. | This study |
| pACYC-Para-*mcr-1* L64F+I65Y | pACYCDuet-1 derivative replacing T7 promoter with arabinose promoter and carrying mcr-1 gene with mutation L64F+I65Y. | This study |
| pACYC-Para-*mcr-1* L165A+I168A | pACYCDuet-1 derivative replacing T7 promoter with arabinose promoter and carrying mcr-1 gene with mutation L165A+I168A. | This study |
| pACYC-Para-*mcr-1* M197A+Y200A | pACYCDuet-1 derivative replacing T7 promoter with arabinose promoter and carrying mcr-1 gene with mutation M197A+Y200A. | This study |
| pACYC-Para-*mcr-1* K204E+K211A | pACYCDuet-1 derivative replacing T7 promoter with arabinose promoter and carrying mcr-1 gene with mutation K204E+K211A. | This study |
| pACYC-Para-*mcr-1* M197E+R184A+K187A | pACYCDuet-1 derivative replacing T7 promoter with arabinose promoter and carrying mcr-1 gene with mutation M197E+R184A+K187A. | This study |
| pACYC-Para-*mcr-1* L64A+I65A+L68A+L69A | pACYCDuet-1 derivative replacing T7 promoter with arabinose promoter and carrying mcr-1 gene with mutation L64A+I65A+L68A+L69A. | This study |
| pACYC-Para-*mcr-1* ΔP188-P195 | pACYCDuet-1 derivative replacing T7 promoter with arabinose promoter and carrying mcr-1 gene with the deletion of residues P188 to P195. | This study |

**Table S9. Primers used in this study**

| Primers | Description | Reference/source |
|---|---|---|
| lpoB-sgRNA1-5F | tagcCGGGTGCCTAAACTGTCCTG | This study |
| lpoB-sgRNA1-3R | aaacCAGGACAGTTTAGGCACCCG | This study |
| lpoB-sgRNA2-5F | tagcGATGGTCAGTAAGATGCTTG | This study |
| lpoB-sgRNA2-3R | aaacCAAGCATCTTACTGACCATC | This study |
| ldtD-sgRNA1-5F | tagcTCGCTGGTCTACTATCAGAA | This study |
| ldtD-sgRNA1-3R | aaacTTCTGATAGTAGACCAGCGA | This study |
| ldtD-sgRNA2-5F | tagcATGTTGCTTAATATGATGTG | This study |
| ldtD-sgRNA2-3R | aaacCACATCATATTAAGCAACAT | This study |
| pbp1b-sgRNA1-5F | tagcGAGGTGTATCTCGGTCAGAG | This study |
| pbp1b-sgRNA1-3R | aaacCTCTGACCGAGATACACCTC | This study |
| pbp1b-sgRNA2-5F | tagcAAATCCGGGAAACCACTGCG | This study |
| pbp1b-sgRNA2-3R | aaacCGCAGTGGTTTCCCGGATTT | This study |
| lpoB-LR-5F | CTGATCATGAATAACCAACCGCC | This study |
| lpoB-LR-3R | TGCGAAACGGCACAAGATTCACCCCTTACAAAATATAG | This study |
| lpoB-RR-5F | TGCCGTTTCGCAGCAATAATCCCATCAC | This study |
| lpoB-RR-3R | CTGACGCGCTATGCACTAAA | This study |
| ldtD-LR-5F | AGCAGCGCGACTGGATGCTC | This study |
| ldtD-LR-3R | ATTTCCCCGAACTACTTCATCCCTTGCCCCCTGTTTTTAT | This study |
| ldtD-RR-5F | ATAAAAACAGGGGGCAAGGGATGAAGTAGTTCGGGGAAAT | This study |
| ldtD-RR-3R | AGTTGCACCGGTTTGCGCGT | This study |
| pbp1b-LR-5F | AAGCCGGATCGCCATTCGTTAT | This study |
| pbp1b-LR-3R | CGGTATTTCACGCTTAGATGGCTTTTTCTCCGCAATATTC | This study |
| pbp1b-RR-5F | TGCGGAGAAAAAGCCATCTAAGCGTGAAATACCG | This study |
| pbp1b-RR-3R | ATCTCTTCGGCGGTCACAACAGAA | This study |
| *mcr-1* NP fuse-F | agataaaatatttctagaAGGAAAAAAGCGAAGGCT | This study |
| *mcr-1* TAA fuse-R | attgagatctgccatatgTCAGCGGATGAATGCGGT | This study |
| pACYC-araC-F | GTCGACGGTATCGATAAGCTTTTATGACAACTTGACGGCTACATCA | This study |
| ParaBAD-mcr1-R | AAACGGGTATGGAGAAACAGTAGAGA | This study |
| ParaBAD-mcr1-F | CTGTTTCTCCATACCCGTTTTTTTGAGTAGTTTCTCATGATGCAG | This study |
| terminator-mcr1-R | CGTCGTTTTACTCAGCGGATGAATGCGGT | This study |
| mcr1-terminator-F | ATCCGCTGAGTAAAACGACGGCCAGTcaaa | This study |
| terminator-pACYC-R | CGCTCTAGAACTAGTGGATCCCAGctgcagTAACACTCGTcga | This study |
| MCR-1-linker delet-F | GTTTCTTTCGCGTGCATAAGATCATGCCAATCTACTCGGTGGG | This study |
| MCR-1-linker delet-R | ACCGAGTAGATTGGCATGATCTTATGCACGCGAAAGAAACTGG | This study |
| MCR-1 L189A-F | gccCGTAGCTATGTCAATCCGATCATGCCAAT | This study |
| MCR-1 L189A-R | TTGACATAGCTACGggcCGGCTTATGCACGCGAAA | This study |
| MCR-1 R190A-F | CTGgccAGCTATGTCAATCCGATCATGCCAAT | This study |
| MCR-1 R190A-R | TTGACATAGCTggcCAGCGGCTTATGCACGCG | This study |
| MCR-1 S191A-F | CTGCGTgccTATGTCAATCCGATCATGCCAAT | This study |
| MCR-1 S191A-R | TTGACATAggcACGCAGCGGCTTATGCACGCG | This study |
| MCR-1 Y192A-F | gccGTCAATCCGATCATGCCAATCTACTCGGT | This study |
| MCR-1 Y192A-R | ATGATCGGATTGACggcGCTACGCAGCGGCTTATGC | This study |
| MCR-1 V193A-F | TAGCTATgccAATCCGATCATGCCAATCTACTC | This study |
| MCR-1 V193A-R | TCGGATTggcATAGCTACGCAGCGGCTTATGC | This study |
| MCR-1 N194A-F | TATGTCgccCCGATCATGCCAATCTACTCGGT | This study |
| MCR-1 N194A-R | ATGATCGGggcGACATAGCTACGCAGCGGCTT | This study |
| MCR-1 M197A-F | AATCCGATCgccCCAATCTACTCGGTGGGTAAGC | This study |
| MCR-1 M197A-R | ATTGGggcGATCGGATTGACATAGCTACGCAG | This study |
| MCR-1 P198A-F | GATCATGgccATCTACTCGGTGGGTAAGCTTGC | This study |
| MCR-1 P198A-R | AGTAGATggcCATGATCGGATTGACATAGCTACG | This study |
| MCR-1 I199A-F | ATCATGCCAgccTACTCGGTGGGTAAGCTTGCC | This study |
| MCR-1 I199A-R | GAGTAggcTGGCATGATCGGATTGACATAGCT | This study |
| MCR-1 Y200A-F | ATCATGCCAATCgccTCGGTGGGTAAGCTTGCCA | This study |
| MCR-1 Y200A-R | GAggcGATTGGCATGATCGGATTGACATAGCT | This study |
| MCR-1 S201A-F | AATCTACgccGTGGGTAAGCTTGCCAGTATTGA | This study |
| MCR-1 S201A-R | TACCCACggcGTAGATTGGCATGATCGGATTGA | This study |
| MCR-1 V202A-F | ATCTACTCGgccGGTAAGCTTGCCAGTATTGAGTATAAA | This study |
| MCR-1 V202A-R | TTACCggcCGAGTAGATTGGCATGATCGGATT | This study |
| MCR-1 G203A-F | ATCTACTCGGTGgccAAGCTTGCCAGTATTGAGTATAAAAAA | This study |
| MCR-1 G203A-R | TTggcCACCGAGTAGATTGGCATGATCGGATT | This study |
| MCR-1 K204A-F | GTgccCTTGCCAGTATTGAGTATAAAAAAGCC | This study |
| MCR-1 K204A-R | AATACTGGCAAGggcACCCACCGAGTAGATTGGCA | This study |
| MCR-1 L205A-F | GTAAGgccGCCAGTATTGAGTATAAAAAAGCCAG | This study |
| MCR-1 L205A-R | AATACTGGCggcCTTACCCACCGAGTAGATTGGC | This study |

| | | |
|---|---|---|
| MCR-1 K187A-F | TCTTTCGCGTGCATgccCCGCTGCGTAGCTATGTCAA | This study |
| MCR-1 K187A-R | ggcATGCACGCGAAAGAAACTGGCATAATGAC | This study |
| MCR-1 I196A-F | TATGTCAATCCGgccTGCCAATCTACTCGGTGGGT | This study |
| MCR-1 I196A-R | CAggcCGGATTGACATAGCTACGCAGCGGCTT | This study |
| Ec-LpxC-F | TTCAGGCGACGGGTGTCGGTTTACA | This study |
| Ec-LpxC-R | CCGGCGCGTTAACTTCGATAACAAT | This study |
| Ec-LdtD-F | TGTCGGCAATCAGTTTGTGCCTGGC | This study |
| Ec-LdtD-R | CGGCATACAACAGTTGAAGCTGGTT | This study |
| Ec-PBP1B-F | CTATGAGGATGAAGAACCGATGCCG | This study |
| Ec-PBP1B-R | GCTTCACCATCTCGTTCTTGCTGAT | This study |
| Ec-LpoB-F | TTGATTACCGCGCTGGCGATGTTTC | This study |
| Ec-LpoB-R | TCTTACTGACCATCGGCTGCATTGC | This study |
| Ec-16S-F | TCAACGATCAGTTCGTGATCGACAG | This study |
| Ec-AmpH-F | GTCTGCTTTTTTCTGCCGTGCT | This study |
| Ec-AmpH-R | TTCACGGTCCCCTGGTCGAGCAATT | This study |
| Ec-16S-R | CGGCCAGTTCACCAAAGTTAAAGGT | This study |
| rseP-ATG-5F | ATGCTGAGTTTTCTCTGG | This study |
| rseP-180bp-3R | aaCATATTCGGTGCCGAG | This study |
| cpxP-5F | ATGCGCATAGTTACCGCT | This study |
| cpxP-180bp-3R | TCGCATCTGCTGACGCTG | This study |
| M197A-Y200A-5F | ATCgcgccaatcgccTCGGTGGGTAAGCTTGCCA | This study |
| M197A-Y200A-3R | CGAggcgattggcgcGATCGGATTGACATAGCTACGCA | This study |
| M197E-5F | TCCGATCgagCCAATCTACTCGGTGGGTAAGC | This study |
| M197E-3R | AGATTGGctcGATCGGATTGACATAGCTACGCA | This study |
| R184A-K187A-M197E-5F | gctgcgtagctatgtcaatccgatcgagCCAATCTACTCGGTGGGTAAGC | This study |
| R184A-K187A-M197E-3R | ttgacatagctacgcagcggcgcatgcacggcAAAGAAACTGGCATAATGACTGCTG | This study |
| L165A-I168A-5F | AGTgctgcgctggctTTACTGCCTGTGGTGGCGTT | This study |
| L165A-I168A-3R | TAAagccagcgcagcACTTGCCACGATCAAGCCC | This study |
| L64F-I65Y-5F | atgctaTtCTAcaccacgctgttatcatcgtatcg | This study |
| L64F-I65Y-3R | gcgtggtgTAGaAtagcatcgcgccaaagagc | This study |
| L64A-I65A-L68A-L69A-5F | gcggccaccacggcggcaTCATCGTATCGCTATGTGCTAAAGC | This study |
| L64A-I65A-L68A-L69A-3R | tgccgccgtggtggccgcTAGCATCGCGCCAAAGAGC | This study |
| L64A-I65A-5F | ATGCTAgcggccACCACGCTGTTATCATCGTATCG | This study |
| L64A-I65A-3R | CGTGGTggccgcTAGCATCGCGCCAAAGAGC | This study |
| K204E-K211A-5F | gagcttgccagtattgagtatgcaaaagccagtgcgccaaaa | This study |
| K204E-K211A-3R | ctcaatactggcaagctCacccaccgagtagattggca | This study |

**Table S10. Synthetic peptides used in this study**

| Peptides | Sequence | Description | Reference/source |
|---|---|---|---|
| 24AA-WT | KPLRSYVNPIMPIYSVGKLASIEY | A 24AA synthetic peptide derived from linker domain of MCR-1 | This study |
| 24AA-2M | KALRSYVNSIMPIYSVGKLASIEY | A 24AA synthetic peptide derived from linker domain of M6 | This study |
| 19AA-2M-tag | KALRSYVNSIMPIYSVGKLWWWWW | A 19AA synthetic peptide derived from linker domain of M6 with five-trytophan modification at C-terminal | This study |
| Peptide MCR-1 | KPLRSYVNPIMPIYSV | A 16AA synthetic peptide derived from linker domain of MCR-1 carrying biotin label at C-terminal | This study |
| Peptide M6 | KALRSYVNSIMPIYSV | A 16AA synthetic peptide derived from linker domain of M6 carrying biotin label at C-terminal | This study |
